# KLF6 in Pulmonary Hypertension: The Dual Role of Friend and Foe

**DOI:** 10.1101/2024.07.08.602619

**Authors:** Rehab Alharbi, Merve Keles, Nadia Fernandes, Hannah Maude, Adam Fellows, Richard D Williams, Chien-Nien Chen, Nathalie Lambie, Nik Matthews, May Al-Sahaf, Minzhe Guo, Lan Zhao, Allan Lawrie, Jeffrey A. Whitsett, Inês Cebola, Beata Wojciak-Stothard

## Abstract

**Background:** Pulmonary arterial hypertension (PAH) is a severe lung condition with unmet clinical needs, marked by endothelial damage, excessive repair, and arterial narrowing, though mechanisms remain unclear.

**Methods:** This study investigates Krüppel-like transcription factor 6 (KLF6), known for its role in tissue injury response and cancer onset, in PAH through functional and expression analyses in human pulmonary artery endothelial cells (HPAECs) and human and rodent PAH lung tissues.

**Findings:** KLF6 expression increased in early experimental PAH in response to hypoxia and inflammation, while the expression of endothelial homeostasis regulators KLF2 and KLF4, previously linked to PAH, decreased. KLF6 overexpression enhanced pulmonary endothelial survival and angiogenesis through broad transcriptomic remodelling, including changes in genes governing endothelial homeostasis and arterial identity (e.g., SOX17, ERG, BMPR2) and promoted human pulmonary artery smooth muscle cells (HPASMCs) proliferation, which was inhibited by bosentan and imatinib. KLF6 functional and transcriptomic responses differed from those of KLF2 and KLF4. Comparative analysis of RNA-seq PAH databases and spatial transcriptomic analysis of human idiopathic PAH (IPAH) tissues highlighted strong association of KLF6 with vascular remodelling, especially with the formation of angioproliferative (plexiform) lesions. High KLF6 expression was observed in IPAH vascular endothelium and IPAH blood-derived endothelial progenitor cells. Single nucleus RNA-seq in PAH associated with Alveolar Capillary Dysplasia confirmed disease-related elevated KLF6 expression in arterial endothelial cells.

**Interpretation:** Accumulation and reorganization of KLF6+ endothelial cells characterize human PAH. KLF6 drives endothelial repair and an apoptosis-resistant, angioproliferative endothelial phenotype. Targeting KLF6 could be a novel therapeutic approach for PAH.

**RESEARCH IN CONTEXT:** *Evidence before this study:* Pulmonary arterial hypertension (PAH) is a progressive and life-shortening lung disease with no cure. In PAH development, endothelial damage is believed to initiate an abnormal repair process, leading to extensive vascular remodelling and the formation of complex angio-proliferative (plexiform) lesions. We conducted a systematic search of the PubMed database to identify transcription factors potentially involved in driving endothelial repair and promoting an apoptosis-resistant, angio-proliferative vascular phenotype. Previous research has linked the loss of endothelial homeostasis in PAH to the inhibition of transcription factors KLF2 and KLF4. While KLF6 is known to play a vital role in vascular development and supports endothelial repair, its specific role in PAH remains unexplored.

*Added value of this study:* This study is the first to establish a connection between KLF6 and PAH pathogenesis. Our findings reveal that KLF6 activation is a key feature of an apoptosis-resistant, angio-proliferative endothelial phenotype characteristic of human PAH-associated plexogenic arteriopathy. Furthermore, we identify both overlapping and unique activation patterns and transcriptional programs regulated by KLF2, KLF4, and KLF6 in lung endothelial cells, highlighting KLF6’s unique role in driving endothelial dysfunction in PAH.

*Implications of all the available evidence:* Targeting KLF6 offers a promising therapeutic strategy to counteract excessive vascular repair and prevent the vascular remodelling in PAH.

**FUNDING:** PhD studentship from the University of Hafr Al Batin, KSA, and the Saudi Cultural Bureau in London (UKSACB) (Rehab Alharbi). Spatial transcriptomic reagents were funded by the British Heart Foundation Centre of Research Excellence Award and Senior BHF Fellowship FS/18/52/33808 (Allan Lawrie). Human samples used in this research project were obtained from the Imperial College Healthcare Tissue Bank (ICHTB) supported by the National Institute for Health Research (NIHR) Biomedical Research Centre based at Imperial College Healthcare NHS Trust and Imperial College London.

## INTRODUCTION

Pulmonary arterial hypertension (PAH) is a progressive and life-limiting vascular lung disease with no cure, despite targeted treatment with multiple therapies^1^. Pathologically, PAH is characterized by sustained pulmonary vasoconstriction and progressive obliteration of resistance pulmonary arteries and arterioles due to medial thickening, intimal fibrosis, and formation of angioproliferative (plexiform) lesions^2^. These changes result in the loss of small distal vessels, often described as “vascular pruning”^3^.

PAH is a multifactorial condition where genetic predisposition^4^, hypoxia^5^, and inflammation^6^ play contributory roles. The most commonly recognized molecular mechanism underlying PAH is the loss of function of bone morphogenetic protein receptor 2 (BMPR2), a TGF-β receptor, either due to mutations or increased protein degradation and reduced membrane recycling^7^. In recent years, mutations in other genes, mostly from the TGF-β superfamily, have also been implicated in PAH^8^.

In the process of PAH development, endothelial damage caused by genetic, epigenetic or environmental factors triggers exaggerated repair, during which clonal expansion of apoptosis-resistant endothelial cells gives rise to complex angio-proliferative (plexiform) lesions^9,10^. Plexiform lesions are a hallmark of human disease, documented in 90% of PAH cases^11^ but are poorly replicated in experimental PAH^12^. Loss of endothelial barrier function, pro-inflammatory and pro-thrombotic activation, endothelial-to-mesenchymal transition (EMT) as well as the proliferation and migration of pulmonary arterial smooth muscle cells (PASMCs) and adventitial fibroblasts are thought to drive pulmonary vascular remodelling^2^. The mechanisms that initiate the formation of plexiform lesions are not known, but a contribution of circulating endothelial progenitor cells^13^ or bronchopulmonary anastomoses^14^ have been suggested.

Endothelial dysfunction in PAH has been attributed to reduced activity of two closely related, flow-activated Krüppel-like transcription factors, KLF2 and KLF4^15,16^. Both KLF2 and KLF4 maintain endothelial homeostasis by controlling expression of anti-proliferative, anti-inflammatory, anti-angiogenic and anti-thrombotic genes^17,18^. Recent data show that expression of KLF6, another member of this family, is significantly upregulated during EMT in HPAECs^19^ and is reduced under conditions of *BMPR2* knockdown in an organ-on-a-chip model of PAH^20^.

KLF6 plays a critical role in vascular development^17^ and aids endothelial repair in wire-denuded arteries ^21^. Changes in the expression of KLF6 have been linked to cancer as well as inflammatory and fibrotic diseases^22^. Of potential relevance to PAH, KLF6 activates endoglin and activin receptor-like kinase 1 (ALK1)^23^, two TGF-β receptors involved in the regulation of pulmonary vascular remodelling^8^. However, the role of KLF6 in PAH or its relationship with the structurally and functionally related transcription factors KLF2 and KLF4 has not been investigated.

This study is the first to provide evidence of KLF6 dysregulation in PAH. We demonstrate that KLF6 is activated in the early stages of the disease in experimental PAH, and that KLF6 and its regulated genes are overexpressed in PAH blood-derived endothelial progenitor cells and PAH lung endothelium. We show that activation of KLF6 is a key feature of an apoptosis-resistant, angio-proliferative endothelial phenotype in plexogenic arteriopathy in PAH. Furthermore, we identify both similarities and differences in the activation patterns and transcriptional programs regulated by KLF2, KLF4, and KLF6 in lung endothelial cells, emphasizing the specific role of KLF6 in promoting endothelial dysfunction in PAH.

## METHODS

Detailed description of experimental protocols is provided in Supplemental Materials and Methods.

### Cell Culture

Human Pulmonary Artery Endothelial Cells (HPAECs) were obtained from PromoCell (Germany, Cat. No. C-12241) and Human Pulmonary Artery Smooth Muscle Cells (HPASMCs) were obtained from Lonza (Walkersville, USA, Cat. No. CC-2581). The cells were used between passages 5-10. Donor information is provided in Table M1 in Supplemental Materials and Methods. Cells were cultured, as described in^15^.

### Peripheral blood mononuclear cells (PBMCs) and endothelial colony forming cells (ECFCs)

Venous blood samples were obtained from healthy volunteers (n=5) and from heritable PAH (HPAH) patients with rare pathogenic BMPR2 variants (n=5)^20^ following informed written consent (REC Ref 17/LO/0563). Demographic and clinical characteristics of HPAH patients and healthy volunteers are shown in Table M2 in Supplemental Materials and Methods.

### Adenoviral overexpression KLF2, KLF4 and KLF6 in HPAECs

The overexpression of most common (long) isoforms of KLF2, KLF4 and KLF6 was achieved by adenoviral gene transfer (AdKLF2-GFP (Cat. No. ADV-213187), AdKLF4-FLAG (Cat. No. ADV-213191), AdKLF6-HA (Cat. No. ADV-213194); Vector Biolabs (Pennsylvania, USA). Adenoviral control (AdCTRL-GFP) for AdKLF2-GFP was AdGFP (Vector Biolabs, Cat. No. 1060), while AdTet-off^24^was kind gift from Professor Stuart Yuspa (National Cancer Institute, NIH, Bethesda, USA) and was used as adenoviral control (AdCTRL) for AdKLF4 and AdKLF6.

### KLF6 silencing

The knockdown of KLF6 expression was performed using silencer select siRNA targeting KLF6 (siRNA-KLF6, Thermo Fisher Scientific, UK, Cat. No. s3376). Control siRNAs included a scrambled siRNA control (siRNA-CTRL Cat. No. 4390846) and a positive siRNA control targeting GAPDH (siRNA-GAPDH, Cat. No. 4390849).

### HPAEC culture under flow

HPAECs were cultured in Nunc Slide Flaskettes (Thermo Fisher Scientific, UK, Cat. No. 170920) until 95% confluent. The bottom slides with the cells were then detached from the flask and placed inside the flow chamber in a parallel flow apparatus^25^. The cells were then subjected to a laminar flow at 4 dynes/cm^2^, physiological for lung arteries^26^, for different periods (30 mins,1h, 4h, 6h and 24h). After the flow exposure, the cells were used for RNA extraction and RT-qPCR to measure mRNA expression levels of KLF2, KLF4 and KLF6.

### Real-time quantitative PCR (qPCR)

Protocols for RNA extraction and RTq PCR are provided in Supplemental Materials and Methods.

### RNA Sequencing

RNA sequencing (RNAseq) (75bp paired-end reads) was performed using an Illumina NextSeq 2000 sequencer (Illumina, USA) at the Imperial BRC Genomics Facility (Imperial College London) to assess the whole transcriptomic effects of KLF2, KLF4, and KLF6 overexpression by adenoviral gene transfer (AdKLF2, AdKLF4 and AdKLF6 conditions, respectively).

The quality of the paired end reads within the fastq files were assessed using FastQC (https://www.bioinformatics.babraham.ac.uk/projects/fastqc/) and MultiQC. To determine transcript abundance values, Salmon (v1.5.2) was used to map sequencing reads in the fastq files to the human genome (GENCODE v44) (https://www.gencodegenes.org/human/) and to obtain gene counts. To identify differentially expressed genes (DEGs), the gene expressions of the AdKLF2, AdKLF4 and AdKLF6 conditions (n=4 biological replicates/condition) were compared with the corresponding negative control conditions using the R package DESeq2 (v1.38.3)^27^, with default settings. Enhanced volcano (v1.16.0) (https://github.com/kevinblighe/EnhancedVolcano 2020) and heatmaps were generated using the R package pheatmap (v1.0.12). Gene annotation was performed using biomaRt (2.46.3). The list of PAH genes was derived from ^28^.

### Gene ontology (GO) and Kyoto Encyclopaedia of Genes and Genomes (KEGG) enrichment analysis

GO and KEGG pathway enrichment analyses were carried out separately on the sets of up- and down-regulated genes from each comparison using the Metascape (https://metascape.org) online tool^29^, the WEB-based GEne SeT AnaLysis Toolkit (WebGestalt) (https://www.webgestalt.org)^30^ and the clusterProfiler package (v3.14.3) in R. The DisGeNET enrichment analysis and network of enriched terms of up- and down-regulated genes was performed using Metascape. Genes that passed the thresholds of FDR <0.01, absolute value of log_2_ (fold change)>2 were used for GO and KEGG pathway enrichment analyses, unless indicated otherwise.

### Comparative analysis with published RNA-seq datasets and disease-gene databases

Comparative analysis was carried out using RNA-seq datasets from HPAECs and ECFCs from the “double hit” microfluidic model of PAH^20^ and from PAECs that had been isolated from IPAH patients^31^ or with known PAH and IPAH-related genes from DisGeNET databases (https://www.semanticscholar.org/paper/GeneOverlap%3A-An-R-package-to-test-and-visualize-Shen/117e12840af966176bbc348db6edf034b0ea479c). Overlapping genes were visualized as UpSet plots using the ComplexUpset package (v1.3.3) in R. The significance of the overlaps between gene sets in comparison to the genomic background was calculated by one-tailed Fisher’s exact test using the GeneOverlap package (v.1.22.0) in R (https://www.semanticscholar.org/paper/GeneOverlap%3A-An-R-package-to-test-and-visualize-Shen/117e12840af966176bbc348db6edf034b0ea479c)

### Spatial transcriptomics

To assess the transcriptomic changes in PAH lungs, spatial transcriptomics of lung tissues from healthy controls and PAH patients were carried out using NanoString GeoMx® to enable high-plex spatial profiling of RNA or protein within specific areas of interest in the tissue ^32^

Formalin fixed and paraffin embedded (FFPE) lung tissue sections (3-4μm) from PAH patients (n[=[6) and age- and sex-matched healthy volunteers (n[=[5) were obtained from the Royal Papworth Hospital NHS Foundation Trust Tissue Bank (Cambridge, UK) with informed written consent (Imperial College Healthcare Tissue Bank Human Tissue Act licence: 12275; REC Wales approval: 22/WA/0214). Donor information for FFPE lung tissue samples is shown in Table M8 in Supplemental Material and Methods.

### Spatial transcriptomics data analysis and visualization

The raw counts were processed using NanoString’s GeoMx NGS pipeline software V2.2 (Nanostring Technologies, USA), where they were converted into digital count conversion (DCC) files. The GeomxTools package (V3.5.0) was used for quality control (QC), and the downstream analysis of the DCC files was performed using R (https://bioconductor.org/packages/GeomxTools/). Differential gene expression analysis was performed using a linear mixed model (LMM) as recommended in the GeomxTools manual. The adjusted p values were calculated using the Benjamini-Hochberg multiple test correction, and differentially expressed genes in all analyses were defined as FDR <0.05 and absolute value of log_2_ (fold change) > 0.25. For spatial deconvolution, the SpatialDecon R package (https://www.nature.com/articles/s41467-022-28020-5) was used to identify the cell type composition of each ROI using the IPF Lung Cell Atlas (https://www.science.org/doi/10.1126/sciadv.aba1983) as the cell profile matrix, which is composed of 36 cell types.

Gene ontology and KEGG pathway analyses were performed using the WEB-based Gene SeT AnaLysis Toolkit (WebGestalt) (https://www.webgestalt.org)

^30^ The ComplexUpset R package (v1.3.3) was used to visualize the number of overlapping genes between different DE gene datasets (https://zenodo.org/records/7314197).

The GeneOverlap R package (v.1.22.0) was used to test the degree and significance of overlap between two gene lists in comparison with a genomic background using Fisher’s exact test (https://www.semanticscholar.org/paper/GeneOverlap%3A-An-R-package-to-test-and-visualize-Shen/117e12840af966176bbc348db6edf034b0ea479c)

### Analysis of endothelial *KLF6* expression in single nucleus RNA sequencing of alveolar capillary dysplasia with misalignment of pulmonary veins (ACDMPV) and control lungs

Integrated single nucleus RNA-seq (snRNA-seq) of ACDMPV and control lungs were obtained from Guo et al. ^33^, which contains snRNA-seq of five ACDMPV lungs (2 weeks to 3.5 years), three 3-year-old control lungs, and three preterm neonate lungs (1-4 days old, 29-31 weeks of gestational age). Detail donor information can be found in Guo et al^33^. We subset the integrated data to the identified endothelial cells (ECs). Gene expression was SoupX-corrected and normalized by Seurat NormalizeData function. Differential expression test was performed in each EC cell type for *KLF6* between ACDMPV vs. 3-yr control lung cells and between ACDMPV vs. preterm neonate lung cells.

Tests were performed using the Seurat 4 FindMarkers function using Wilcoxon Rank Sum test. Genes with the following criteria were considered up in an EC subtype in ACDMPV: p_val<0.05, log2FC>=log2(1.5), pct.1>=0.2, “ident1.pct>=20.nSample” >=2 (i.e., pct>=20% in at least 2 ACDMPV samples) in the cell type. Genes with the following criteria were considered down in an EC subtype in ACDMPV: p val<0.05, log2FC <= -log2(1.5), pct.2>=0.2, “ident2.pct>=20.nSample” >=2 (i.e., pct>=20% in at least 2 Control or 2 Preterm samples) in the cell type.

### Immunocytochemistry (ICC) and Immunohistochemistry (IHC)

Protocols and reagent lists are provided in Supplemental Materials and Methods.

### NF-**κ**B Luciferase Reporter Assay

HPAECs were left untreated or were infected with adenoviral NF-κB luciferase reporter^34^ (AdNFkB-luc, Vector Biolabs, Cat. No.1740), AdCTRL and AdKLF6 or were transfected with siRNA-KLF6 and scrambled-siRNA control, as required. Since the basal KLF6 expression in unstimulated, quiescent HPAECs was insufficient to produce a measurable effect, the effects of KLF6 silencing were studied in cells cultured under flow. 3 hours after adenoviral infection or 48 hours after siRNA exposure, the culture media were replaced with a fresh EGM2 medium with or without 10 ng/mL of TNF-α (R&D Systems, USA, Cat. No. 210-TA-020). The cells were then incubated under either normoxic or hypoxic conditions for 24 hours. A luciferase reporter assay (Promega, USA, Cat. No. E1500) was then performed according to the manufacturer’s instructions. The intensity of luminescence, proportional to the level of NF-κB–driven luciferase activity, was measured in GloMax® luminometer (Promega, UK).

### Transwell permeability assay

Endothelial permeability assay was measured in cells grown in Transwell plates (Corning, USA, Cat. No. 3413). Passage of 40 kDa FITC-Dextran (Sigma-Aldrich, Dorset, UK, Cat. No. FD40S) following cell treatments was measured at excitation/emission 490/525 nm using a GloMax® luminometer (Promega, UK)^15^.

### Angiogenesis assays

Angiogenesis was assessed in a tube formation assay *in vitro* and in an *ex vivo* pulmonary arterial explants sprouting assay. For the tube formation assay, HPAECs infected with either AdCTRL or AdKLF6 were seeded into a 96-well plate pre-coated with growth factor-reduced Matrigel (Scientific Laboratory Supplies, UK, Cat. No. 354230). After 20 hours of incubation, endothelial tube formation was imaged using a phase*-*contrast microscope. Images were then analysed using ImageJ software (Fiji, version 2.14.0) with an Angiogenesis Analyzer plugin to quantify the number of nodes and meshes as well as the total tube length.

For the arterial explants sprouting assay, surgical specimens of human pulmonary arteries were cleaned of blood and attached connective tissues in PBS. Donor characteristics – including age, gender, height, weight and diagnosis – are summarized in Table M11. AdCTRL or AdKLF6 was added on top of the endothelial layer, and the explants were then incubated for 3 hours in a humidified incubator. Afterwards, the arterial tissue was washed, cut into roughly 1 mm^2^ fragments which were then embedded in Matrigel and incubated for 3 weeks in a humidified incubator under normoxic conditions. Endothelial sprouts were fluorescently labelled with Vybrant™ CFDA SE Cell Tracer (Invitrogen, USA, Cat. No. V12883) before being imaged under a fluorescent microscope. The total area of sprouting was measured using ImageJ software (Fiji, version 2.14.0).

### EdU Cell Proliferation Assay

Cell proliferation assay was performed using an EdU Cell Proliferation Assay Kit (EdU-594, EMD Millipore Corp, USA, Cat. No. 17-10527), according to the manufacturer’s instructions.

### Proliferation of PASMCs in co-culture with HPAECs

Control (AdCTRL) and AdKLF6-overexpressing HPAECs were grown to confluence in 6.5 mm Transwell inserts with 1.0 µm pore size transparent PET membrane (Corning, Cat. No 353104) in EGM2 culture media containing serum and growth factors. HPASMCs were seeded at the bottom of 24-well plates (50 x10^3^ cells/well) or on the other side of empty Transwell inserts or inserts containing endothelial cells (20 x 10^3^ cells/insert) in EGM2 medium containing 5% FBS and no growth factors. Following 2-hour incubation in a humidified culture incubator, unattached PASMCs were washed away with PBS and Transwell inserts containing HPAECs or co-cultures of HPAECs and HPASMCs were inserted into the wells of 24 well plate. Culture media were replaced with EGM2 medium containing 0.2% FBS, antibiotics and EdU (for further detail please see EdU Cell Proliferation Assay), with no growth factors. In some wells, ET-1 receptor antagonist bosentan^35^ and tyrosine kinase inhibitor, imatinib^36^ were added at clinically relevant doses, previously shown to inhibit ET-1-induced PASMC proliferation (bosentan, Stratech S3051; 20 µmol/L)^35^ and PDGF-β-induced PASMC proliferation (imatinib, Enzo ALX-270-492; 10 µmol/L)^20,36,37^. PASMCs cultured in full PASMC medium served as a positive control.

### Wound healing assay

Endothelial cell migration was assessed in a wound healing assay using ibidi Culture-Insert 2 Well in µ-Dish 35 mm (ibidi, Germany, Cat. No. 81176) or in a scratch assay under flow.

The wound area was measured using a custom-made Wound Healing Tool ImageJ macro (written by Stephen Rothery, Facility for Imaging by Light Microscopy (FILM), Imperial College London).

### Apoptosis Caspase-Glo 3/7 Assay

Apoptosis was assessed by measuring caspase 3/7 activity using a Caspase-Glo 3/7 Assay Kit (Promega, Southampton, UK, Cat. No. G8091), according to the manufacturer’s instructions.

### TUNEL Apoptosis Assay

TUNEL Assay was performed using a Click-iT Plus for In Situ Apoptosis Detection Kit with Alexa Fluor 647 dye (Invitrogen, Cat. No. C10619), according to the manufacturer’s instructions.

### Study approval

Venous blood samples were obtained with the informed written consent and approval of the local ethics committee (REC Ref 17/LO/0563) from healthy volunteers and from HPAH patients with a rare pathogenic variant of *BMPR2* gene.

Human pulmonary artery tissue samples were obtained following ethical approval from the Imperial College Healthcare Tissue Bank (ICHTB HTA licence: 12275; REC approval: 17/WA/0161).

Formalin fixed and paraffin embedded lung tissue sections from healthy volunteers and PAH patients were obtained from the Royal Papworth Hospital NHS Foundation Trust Tissue Bank (Cambridge, UK) with informed written consent and ethical approval by (ICHTB HTA licence: 12275; REC Wales approval: 22/WA/0214).

### Statistical Analysis

All graphs and statistical analyses were performed using either GraphPad Prism software 9 (GraphPad Software, USA) or Rstudio (RStudio Inc, version 1.2.5042).

All experiments were performed in at least four biological replicates, with 3 technical replicates performed per experiment, unless stated otherwise. All data were tested for normal distribution using the Shapiro-Wilk test. An unpaired student’s t test was used to analyse the normally distributed data from two sample groups, while a one- or two-way ANOVA test was used to analyse three or more sample groups as appropriate.

Statistical significance was accepted when p*-*values were less than 0.05. All error bars are representative of mean (*±* SEM).

## RESULTS

### *KLF6* expression is increased under PAH-related vascular stress conditions

Given its key role in the regulation of vascular repair, we examined whether hypoxia and/or inflammation - two important triggers of PAH^38^ - modulate KLF6 levels in rodent models of PAH, specifically in chronically hypoxic mice and monocrotaline (MCT) rats and cultured human pulmonary artery endothelial cells (HPAECs). In mice, development of PAH was confirmed by elevated right ventricular systolic pressure (RVSP), right ventricular hypertrophy (RVH), and increased pulmonary arterial muscularization (Supplementary Figure S1). The PAH phenotype of the MCT rats is reported in our previous study^39^.

Mice exhibited a 4-fold increase in lung *KLF6* expression after 24 hours of hypoxic exposure (p<0.0001), which returned to control levels 2 weeks later, when the disease was established (Figure 1A). An early rise in *KLF6* expression was also observed in monocrotaline (MCT) rats (Figure 1B), which typically display a more severe phenotype and greater inflammatory response, compared to chronically hypoxic animals^12^ (Figure 1C).

**Figure 1.**
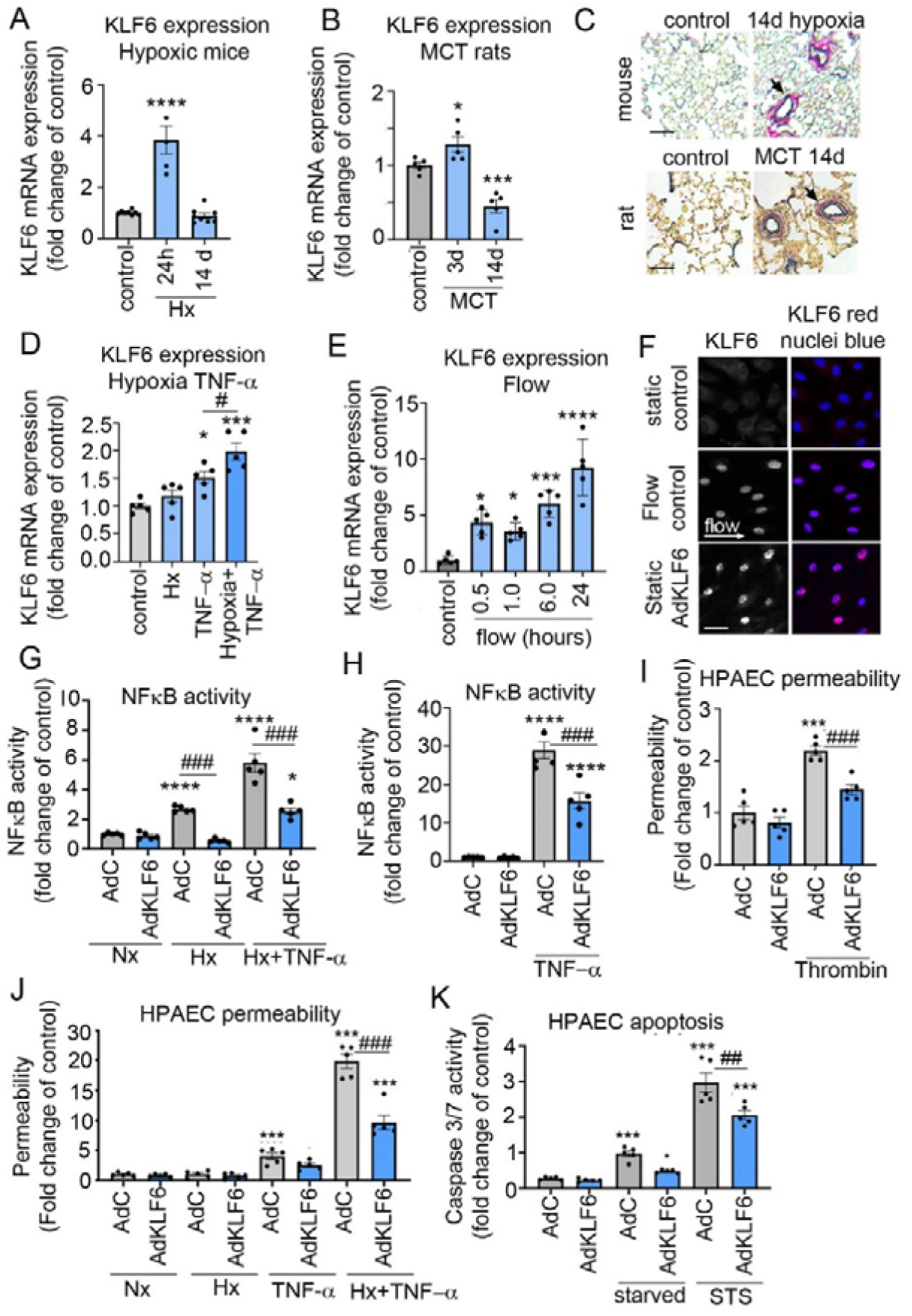
KLF6 in endothelial responses to PAH-associated stress conditions. **(A)** KLF6 mRNA expression changes in hypoxic mice (10% O_2_) and **(B)** MCT rats at different experimental time points, as indicated. **(C)** Corresponding representative images of healthy and PAH mouse and rat lung tissues; arrows point to muscularised vessels (EVG); Bar=25 µm. **(D**) KLF6 mRNA expression in HPAECs exposed to hypoxia (2% O_2,_ 24h) with, or without TNF-α (10 ng/ml, 24h) or **(E)** in HPAECs exposed to flow (4 dynes/cm^2^ 0-24h). n=5. **(F)** Localization of KLF6 in HPAECs under flow or HPAECs infected with AdKLF6. In merged images, nuclei are blue and KLF6 is red, confocal microscopy. Scale bar = 10 μm. **(G)**, (**H**) and **(I)** show effects of KLF6 overexpression on NF-κB activity in HPAECs treated with TNF-α (10ng/ml, 24h), hypoxia (2% O2, 24h) or thrombin (1U/mL, 1h), as indicated. AdCTRL: adenoviral control; AdKLF6: adenoviral KLF6. **(J)** Effect of KLF6 overexpression on endothelial permeability in hypoxia (Hx) and TNF-a-treated HPAECs, as indicated. **(K)** Effect of KLF6 on starvation- and staurosporine (STS)-induced endothelial apoptosis. In all graphs, bars show mean fold-changes of control±SEM. KLF6 expression was normalised to B2M. *P<0.05, ***P<0.001, ****P<0.0001, comparisons with untreated or viral controls, as appropriate. #P<0.05, ##P<0.01, ###P<0.001, comparison with corresponding treatment controls, as indicated. One-way ANOVA with Tukey’s post-hoc test; (A) n=4-7, (B-K) n=5.

To mimic the PAH-associated vascular stress conditions *in vitro*, human pulmonary artery endothelial cells (HPAECs) were exposed to hypoxia (2% O[) and TNF-α, either separately or in combination. The combined treatment was performed to establish the “double hit” conditions, which have been shown to impose an additional burden that predisposes to the disease and induce a more severe vascular phenotype in experimental PAH models^40,41^. When combined, hypoxia and TNF-α induced a more pronounced increase in *KLF6* expression in HPAECs than either factor alone (Figure 1D). Notably, while *KLF6* expression was upregulated, the expression of *KLF2* and *KLF4* was reduced under the same experimental conditions, both *in vivo* and *in vitro* (Supplementary Figure S2A-D).

KLF2- and KLF4 regulate endothelial adaptation to flow^42^ and their expression is reduced in PAH, likely to contribute to endothelial damage^43^. Flow exposure increased *KLF2, KLF4* and *KLF6* expression in HPAECs, but expression pattern of *KLF6* differed from those of *KLF2* and *KLF4*: the peak response for KLF6 occurred at later time points following the onset of flow (Supplementary Figure S2E and F), likely indicating distinct roles for these transcription factors in the endothelial response to mechanosensory stimulation.

Altogether these analyses reveal that KLF6 is over-expressed in both *in vitro* and *in vivo* models of PAH and follows a distinct activation pattern from the previously implicated in PAH KLF2 and KLF4.

### KLF6 improves endothelial survival and stimulates angiogenesis

To investigate the role of stress induced KLF6 activation, KLF6 was overexpressed in HPAECs via adenoviral gene transfer. KLF6 exhibited nuclear localization, both when exogenously expressed and when induced by the onset of flow (Figure 1F and Supplemental Figures S2G, H).

The overexpression of KLF6 inhibited the activation of the pro-inflammatory transcription factor NFκB in HPAECs exposed to hypoxia and TNF-α (Figure 1G and H), while silencing of KLF6 had an opposite effect (Supplemental Figure S3A-C).

KLF6 overexpression enhanced endothelial resilience and improved endothelial survival. KLF6 attenuated hypoxia-, TNF-α- and thrombin-induced breakdown in endothelial barrier function (Figure 1I, J) and inhibited HPAEC apoptosis induced by serum starvation and treatment with staurosporine (STS)^44^ (Figure 1K). Conversely, KLF6 silencing markedly enhanced inflammatory activation and apoptosis in HPAECs (Supplementary Figure S3D).

In PAH, endothelial damage is thought to trigger exaggerated repair, which leads to the emergence of highly proliferative, angiogenic, cancer-like cells, contributing to intimal hyperplasia and the formation of plexiform lesions^45^.

In addition to its pro-survival effects, KLF6 markedly enhanced endothelial angiogenesis, as evidenced by increased capillary network formation in KLF6-overexpressing HPAECs (Figure 2A-D). Increased angiogenic sprouting was also observed in human pulmonary artery explants transduced with AdKLF6 (Figure 2E, F). Adenoviral transduction efficiency in the arterial endothelium exceeded 80%, similar to the transduction efficiency observed in cultured HPAECs (Supplementary Figure S4).

**Figure 2.**
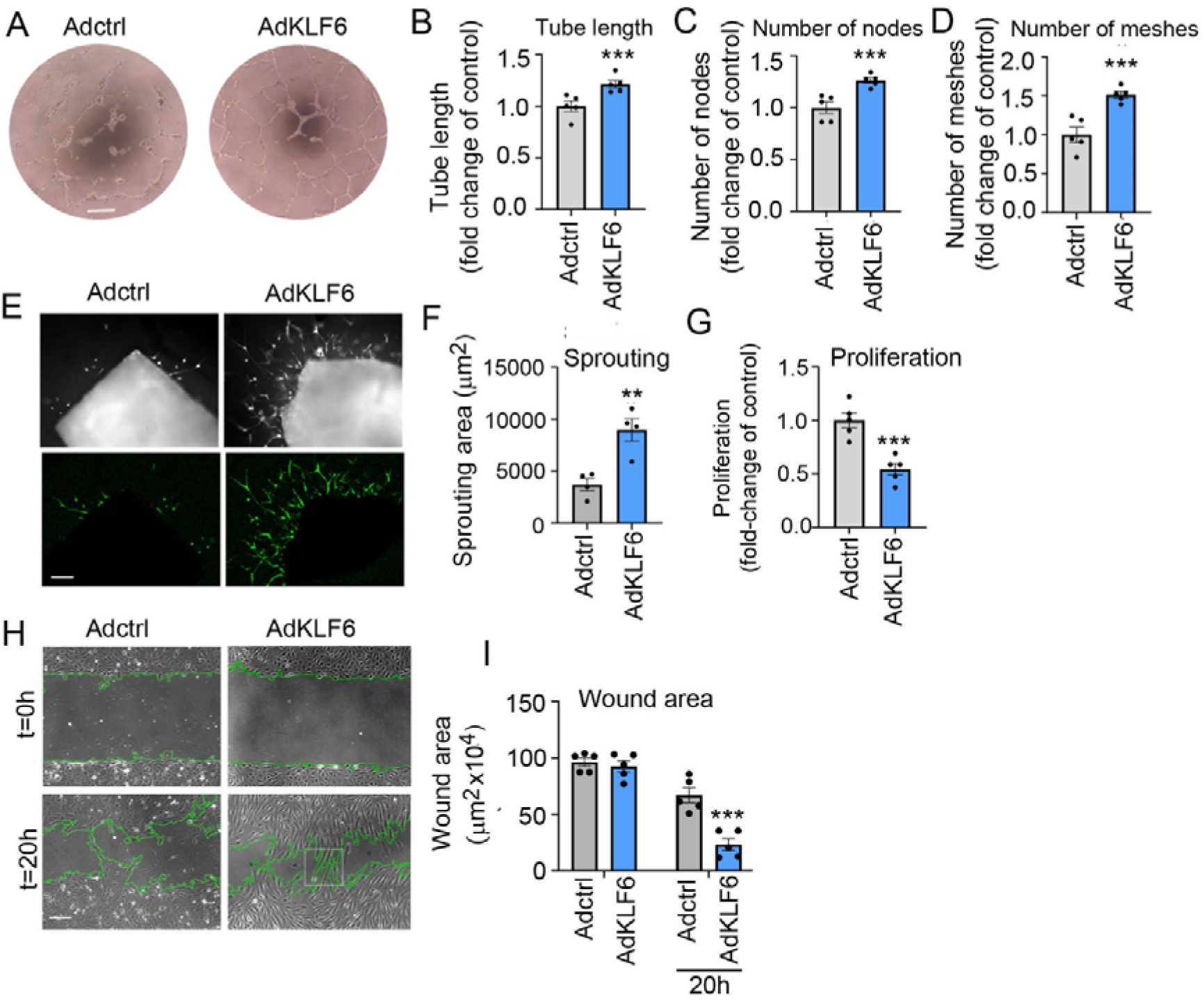
Effects of KLF6 on endothelial angiogenesis, proliferation and migration. **(A)** Images of tube formation in AdCTRLand AdKLF6-treated HPAECs; phase contrast microscopy. Scale bar = 50μm. **(B)** Total tube length, **(C)** number of nodes and **(D**) meshes in HPAECs treated, as indicated. ImageJ with Angiogenesis Analyzer plugin. **(E)** Sprouting angiogenesis in human pulmonary artery explants treated with AdCTRLor AdKLF6. Explants were labelled with Vybrant CFDA SE cell tracker green, fluorescent microscopy. Scale bar=50μm. **(F)** Sprouting area in explants treated, as indicated. n=4 donors; 8 explants/donor were analysed. **(G)** HPAEC proliferation; EdU incorporation assay. Images in **(H)** and corresponding graph in **(I)** show endothelial wound healing in control and KLF6-overexpressing HPAECs. Boxed area shows well polarised AdKLF6-overexpressing cells migrating into the wound area. Wound edges are traced in green. In (B, D, F, G) ***P<0.001, comparison with adenoviral controls, unpaired t-test; In (I) **P <0.01, ***P<0.001, comparison with adenoviral controls; two-way ANOVA with Tukey’s post-hoc test; n=4-5. Bars show mean ±SEM.

Angiogenesis is a complex process involving endothelial cell proliferation, migration, and junctional remodelling. To gain further insights into the mechanisms involved, we studied endothelial proliferation and migration following manipulation of KLF6 expression in HPAECs. KLF6 inhibited HPAEC proliferation (∼ 2-fold decrease, p < 0.001) (Figure 2G), but markedly increased endothelial migration, resulting in accelerated endothelial wound closure *in vitro*. Enhanced endothelial wound repair was observed under both optimal and serum- and growth factor-reduced culture conditions (Figure 2H-I and Supplementary Figure S5 A, B). Conversely, KLF6 silencing markedly reduced endothelial wound repair (Supplementary Figure S5 C, D).

In summary, functional studies suggest that KLF6 is induced under vascular stress conditions to reduce endothelial damage and stimulate endothelial repair and angiogenesis.

### KLF6-regulated endothelial transcriptome HPAECs under healthy and disease conditions

KLFs can act as either transcriptional activators or repressors^46^, and the functional assays described above suggest a key role for KLF6 in regulating endothelial gene programs. Thus, we conducted RNA-seq analysis on HPAECs infected with AdCTRL or AdKLF6 (n=4 biological replicates/condition), to characterise the set of endothelial genes regulated by KLF6 (Figure 3A). The viral transduction was optimised to induce a ∼6-fold increase in KLF6 protein (Figure S2F), comparable with changes seen during placental differentiation^47^ and endothelial injury^21^. We detected a total of 4,732 differentially expressed genes (DEGs) in HPAECs overexpressing *KLF6* (p-value < 0.05, log_2_ |FC| <-0.25 or >0.25), including 2,332 downregulated and 2,400 upregulated genes. This included 204 downregulated and 157 upregulated genes passing a more stringent criteria of FDR <0.01 and log_2_ |FC|>2 (Supplementary File S1). Heatmap with top 50 KLF6-upregulated and downtegulated genes is shown in Figure S6.

**Figure 3.**
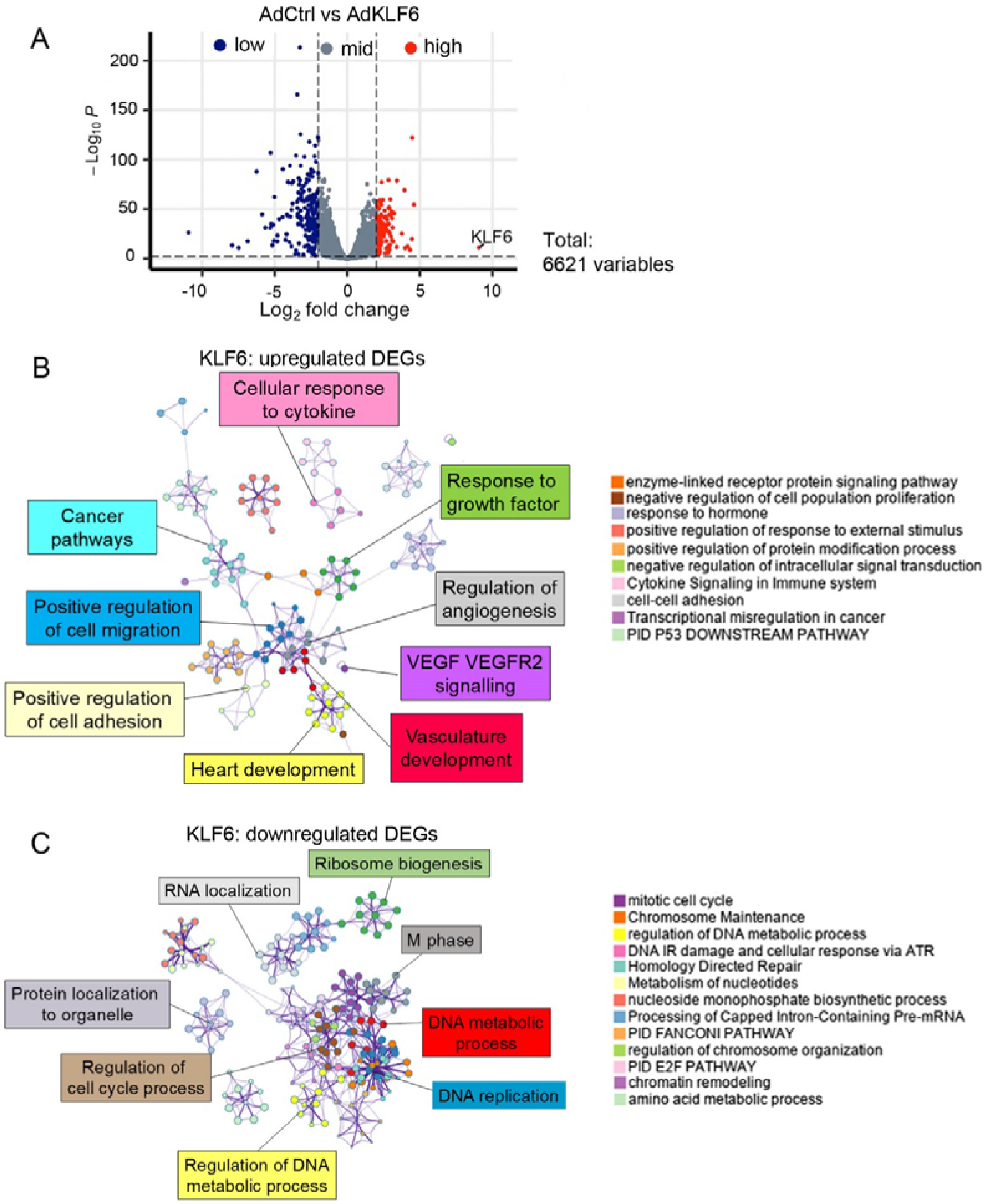
Enrichment analysis of KLF6 DEGs. **(A)** Volcano plot showing differentially expressed genes (DEGs) in AdKLF6 and AdCTRL treated HPAECs. Downregulated genes are blue and upregulated genes are red. p<0.05 and Log_2_ fold change <-1 or >1. Metascape network of enriched terms in **(B**) upregulated and **(C)** downregulated DEGs coloured by cluster ID.

Metascape^29^ gene annotation analysis showed that the genes upregulated upon *KLF6* overexpression were significantly enriched in annotation terms associated with endothelial homeostasis and function, including angiogenesis, cell adhesion and migration, vascular development and VEGF signalling (Figure 3B). Notably, KLF6 enhanced expression of several key regulators of arterial identity, repair and angiogenesis, including *ERG, BMPR2, SOX17, KDR, TIE1, TIE2, HES1*, *CDH5, PECAM1* and *FLT1*. In contrast, the downregulated genes were mainly enriched in generic terms, including cell cycle progression, DNA replication and DNA metabolic process (Figure 3C). Changes in the expression of selected genes involved in endothelial homeostasis and function were replicated by qPCR in an additional set of HPAECs (Supplementary Figure S7).

Our initial functional analyses suggested that KLF6 modulates the response of HPAECs to hypoxia and TNF-α (Figure 1G, H, Supplemental Figure S3A-C). We therefore set out to investigate at the transcriptional level the impact of KLF6 on the endothelial response to these two key PAH contributory factors.

RNA-seq analysis showed that HPAEC exposure to either hypoxia or TNF-α (10 ng/mL, 24 hours) alone resulted in detectable changes in gene expression. Two hundred genes were modulated by hypoxia (FDR <0.05, log_2_|FC<-0.25 or >0.25; comparison AdCTRL vs AdCTRL+ hypoxia), enriched, as expected, in HIF-1 and TGF-β signalling pathways (Supplementary Figure S8A and Supplementary File S2). The response to TNF-α exposure showed 2,788 DEGs (FDR <0.05, -0.25 >log_2_|FC|>0.25; comparison AdCTRL vs AdCTRL+ TNF-α), significantly enriched for TNF-α and NF-κB signalling pathways (Supplementary Figure S9A and Supplementary File S3).

We then compared the transcriptional signatures of HPAECs exposed to hypoxia or TNF-α-induced inflammation with, or without of KLF6 overexpression. KLF6 overexpression under hypoxic conditions revealed 1,576 DEGs (FDR <0.05, log2|FC|<-0.25 or >0.25) which showed significant association with inflammatory responses, autophagy, pathways controlling cell proliferation, growth factor and oestrogen signalling (Supplementary Figure S8B and Supplementary File S4).

Upon overexpression of KLF6, TNF-α-treated HPAECs showed differential expression of 9,395 genes (FDR < 0.05, log_2_|FC|> 0.25, Supplementary File S5). The upregulated genes were enriched in FOXO, p53, PI3K-AKT, JAK-STAT, TNF-α and NF-κB signalling pathways. The inhibitory effect of KLF6 on TNF-α-induced NF-κB activity in cultured HPAECs is consistent with previously reported anti-inflammatory actions of KLF6^48-50^ and may result from increased expression of NF-κB Inhibitor Alpha (*NFKBIA*). However, KLF6-induced expression of IL-6, typically seen during acute response to endothelial injury, may augment immune responses under sustained vascular stress conditions *in vivo*.^51^

In both treatment groups, KLF6-downregulated genes were enriched in cell cycle and DNA replication, consistent with anti-proliferative effects of KLF6 overexpression seen in cultured HPAECs (Supplementary Figures S8B and S9B).

In summary, transcriptomic profiling and pathway analysis of KLF6-overexpressing HPAECs reaffirms the role of KLF6 as a regulator of arterial identity, endothelial survival, repair and angiogenesis and an upstream regulator of numerous critical genes and pathways implicated in the development of PAH.

### Comparative analysis of KLF2-, KLF4 -and KLF6-induced transcriptomic changes in HPAECs reveals unique effects of KLF6

KLF2, 4 and 6 are expressed in the pulmonary endothelium and share numerous structural and functional similarities but their individual roles have not been well delineated ^18,22^. To identify shared as well as unique genes regulated by these three transcription factors, RNA-seq was conducted on control HPAECs, as well as HPAECs overexpressing either one of these KLF factors. Adenoviral infection was optimised to ensure a comparable ∼6-fold increase in protein expression in all KLFs (Supplementary Figure S10 A-D).

KLFs are major regulators of broad gene expression programs. Indeed, the extent of perturbation of the endothelial transcriptome by overexpression of either KLF2 or KLF4 was similar to that observed in KLF6-overexpressing cells (4,732 DEGs), with 4,256 and 4,622 DEGs for KLF2- and KLF4-overexpressing HPAECs, respectively (p-value <0.05, log_2_|FC|>0.25) (Supplementary Figure S10 E, F and Supplementary Files S6 and S7).

We compared all three transcriptomic datasets, focussing on the set of genes showing the strongest differential expression (FDR<0.05, log_2_|FC|>2). The analysis revealed individual and shared contributions of KLFs to regulating endothelial transcription. Namely, 86 DEGs unique to KLF6 were enriched in key functions promoting endothelial repair such as wound healing, vasculature development, TGF-B signalling, response to DNA damage (Figure 4A, B). The 104 DEGs shared among KLF2, KLF4 and KLF6 (Figure 4C) showed enrichment in processes controlling cell proliferation such as DNA metabolism, DNA replication and cell cycle regulation. Lists of unique and shared KLF2, KLF4 and KLF6-regulated DEGs are provided in Supplementary Tables S1 and S2.

**Figure 4:**
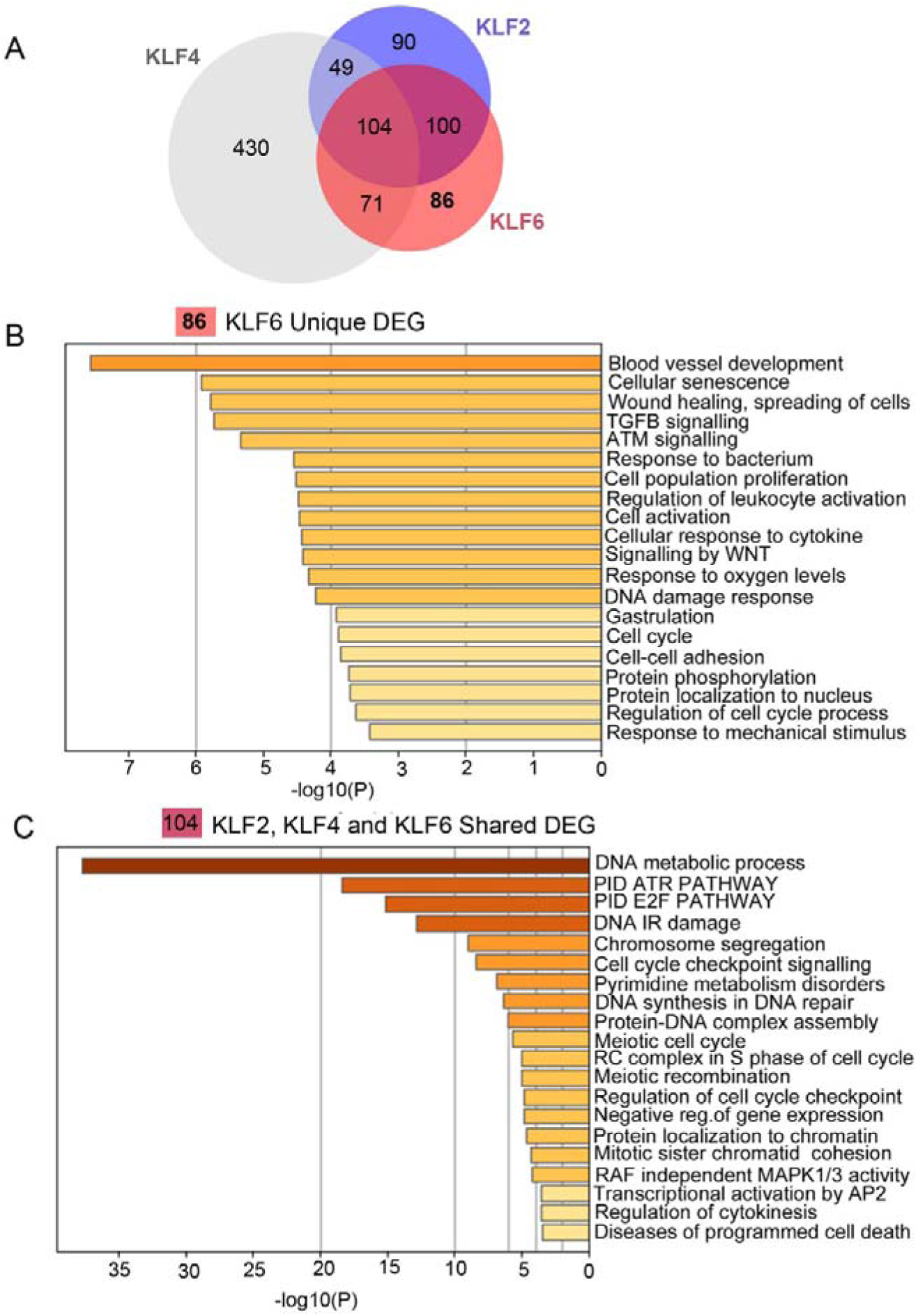
Metascape pathway and functional enrichment analysis of shared and unique KLF6-regulated DEGs. **(A)** Venn diagram shows the number of KLF2-, KLF4- and KLF6-regulated DEGs, with DEGs unique to KLF6 marked in red. **(B)** A corresponding bar graph shows Metascape pathway and process enrichment analysis of unique KLF6-regulated DEGs. **(C)** Venn diagram shows the number of shared KLF2-, KLF4- and KLF6-regulated DEGs in HPAECs, where shared DEGs are marked in red. **(D)** Metascape pathway and process enrichment analysis of shared DEGs. DEG selection criteria were FDR<0.05, log2|FC|>2 or <-2

Transcription factors are often part of complex regulatory networks whereby they can regulate each other via feedback loops and/or work within a specific hierarchy. We therefore harnessed the transcriptomic datasets corresponding to overexpression of different KLFs to investigate their relationships in HPAECs. The analysis of mutual regulatory interactions revealed that KLF6 increased expression of KLF2, but KLF2 inhibited expression of KLF6 (Suppl. Fig S11 A, B). This result suggests that KLF2, which promotes endothelial quiescence and homeostasis, may act to restrict the actions of KLF6. No significant regulatory relationship was observed between the expression of KLF6 and KLF4 even though KLF4-overexpressing cells appeared to express more KLF6 (p=0.056, Supplementary Figure S11 C, D).

Collectively, these analyses suggest that KLF6 plays a unique role in orchestrating pulmonary endothelial response to injury. KLF6 can function in conjunction with KLF2 and KLF4 to inhibit endothelial proliferation, apoptosis, and inflammatory activation. The net effect of KLF6 activation, supported by functional and transcriptomic analyses, is twofold: minimizing injury and promoting endothelial regeneration.

### KLF6-regulated genes in PAH

The transcriptomic analyses described above suggest a core role for KLF6 in lung endothelial homeostasis. This prompted us to investigate the relationship between KLF6 and PAH-relevant genes. DisGeNET disease enrichment analysis^52^ revealed significant associations of KLF6-upregulated genes with vascular diseases, including PAH, whereas the down-regulated genes showed links with mitochondrial diseases and hypertrophic cardiomyopathy (Figure 5A and B). Interestingly, KLF6 altered the expression of numerous genes with known PAH-linked mutations, including *BMPR2, ENG, THBS1, SOX17, KDR, CAV1, ACVRL1, SMAD1, SMAD4, EDN1, ATP13A3, KLK1, NOTCH3, TOPBP1* and *BMPR1B* (Figure 5C). KLF6 DEGs also showed significant overlaps with publicly available sets of differentially expressed genes in PAH and IPAH, including PAH HPAECs and ECFCs from microfluidic models of PAH ^20^, IPAH PAECs^31^ and a previously curated list of PAH-associated genes (named herein as “known PAH genes”)^53^ (Supplementary Figure S12 and Supplementary Table S3).

**Figure 5.**
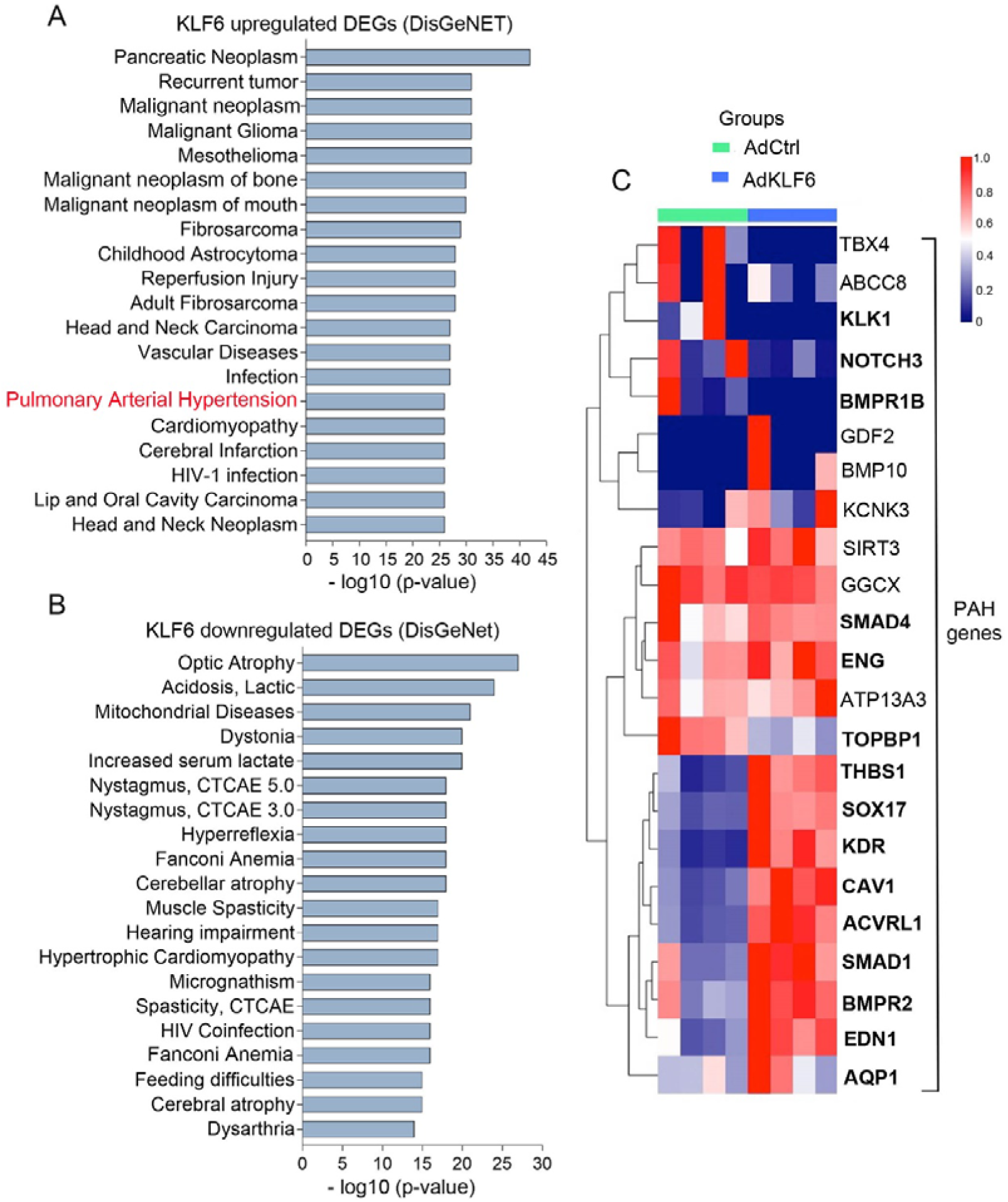
DisGeNET enrichment analysis of KLF6-regulated DEGs. Disease gene enrichment analysis of **(A)** upregulated and **(B)** downregulated KLF6 DEGs. Pulmonary hypertension terms are highlighted in purple **(C)** Heatmap shows colour-coded changes in expression of PAH-associated genes, with hierarchical clustering in control and KLF6-overexpressing HPAECs. DEGs are marked in bold. Each column represents one experimental repeat; n= 4/group.

Considering that KLF6-induced increase in expression of endothelin-1^54^ and TGF-β 1family genes^55^ in pulmonary arterial endothelial cells may potentially affect the behaviour of underlaying arterial smooth muscle cells, we studied HPASMC proliferation in co-culture with AdCTRL- and KLF6-infected HPAECs (Supplementary Figure S13). Endothelial KLF6 overexpression significantly stimulated PASMC proliferation and this effect was prevented by a combined treatment with of ET-1 receptor antagonist, bosentan^35^ and PDGF-β receptor inhibitor, imatinib^36,37^ at clinically relevant doses (Supplementary Figure S13).

To further investigate the contribution of KLF6 to vascular remodelling in PAH and to analyse site-specific changes in gene expression, spatial transcriptomic analysis of remodelled human PAH vasculature was carried out, with two separate categories, one for the concentric intimal lesions with different degrees of medial and adventitial thickening and the other one for vessels located within the complex plexiform lesions (Figure 6A) [30 ROIs/lung in sections from patients with (n=6)] and equivalent sixed pulmonary arterioles from age- and sex-matched healthy controls (n=5).

**Figure 6.**
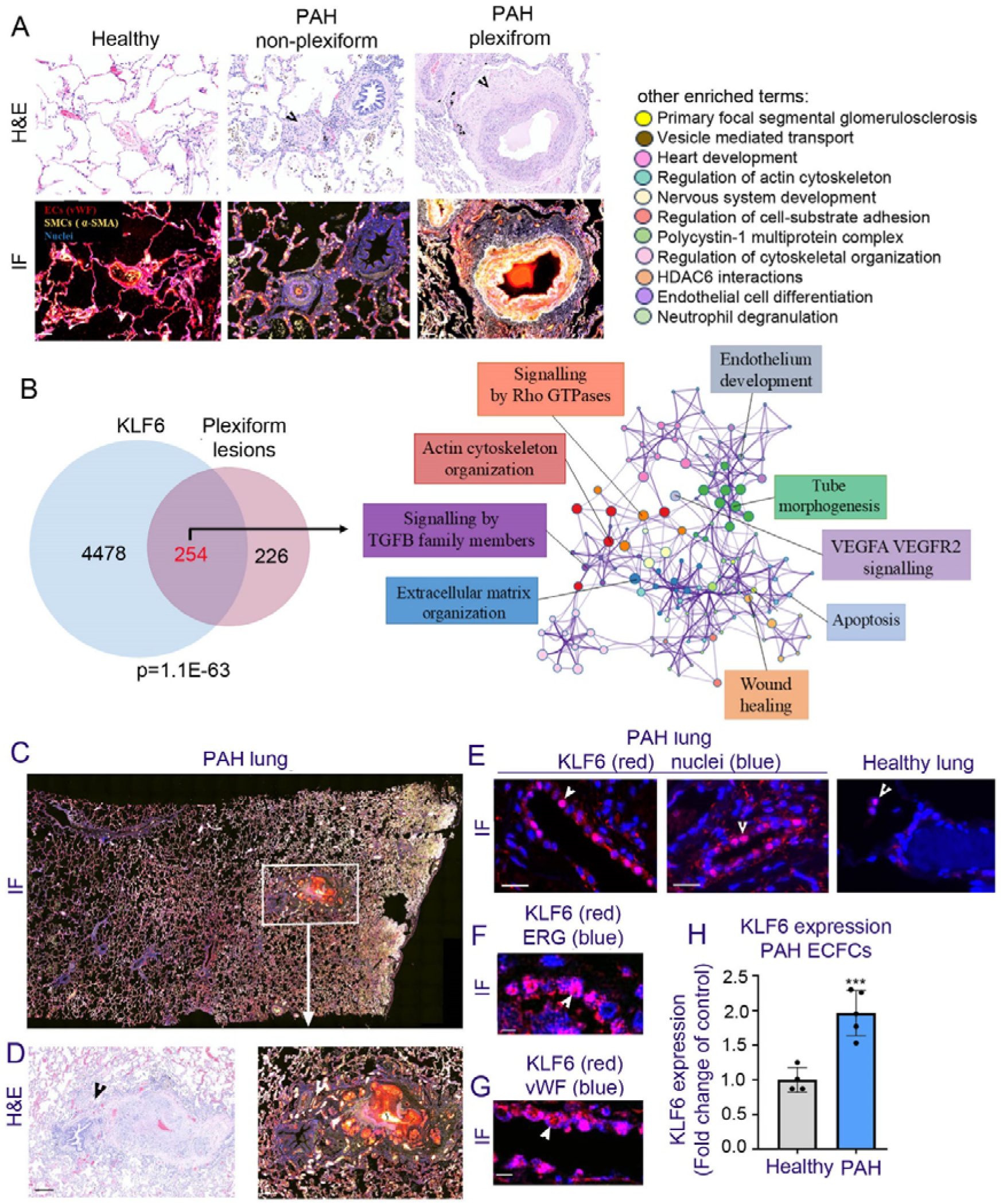
KLF6 in PAH endothelium. **(A)** Representative examples of healthy lung tissues, non-plexiform remodelled vessel and plexiform lesion in PAH lung used for spatial transcriptomics. H&E and IF staining was used to guide the selection of ROIs in GeoMx Digital Spatial Profiler. Von Willebrand factor (vWF; red), α-smooth muscle actin (α-SMA; yellow) and (SYTO 13, blue). 30-45 ROIs were selected in lung sections from healthy (n=5) and PAH (n=6) individuals, with separate annotation for non-plexiform plexiform lesions. **(B**) Venn diagram showing an overlap between KLF6-regulated DEGs in HPAECs and plexiform lesions DEGs identified by spatial transcriptomics; P= 9.7E-53, Fisher’s exact test in the GeneOverlap package in R. The associated Metascape network of enriched terms in the shared pool of DEGs coloured by cluster ID, is shown on the right. FDR<0.05, log2|FC|<-0.25 or >0.25 **(C)** PAH lung section (Immunofluorescence) with a plexiform lesion within the boxed area. **(D)** Enlarged images of the plexiform lesion (H&E and IF staining), with arrowheads pointing to the area enlarged in (E). **(E)** Localization of KLF6 in in PAH and healthy lung, as indicated. KLF6 is red and nuclei are blue; Arrowheads point to nuclear localization of KLF6 (pink). **(F, G)** show KLF6, vWF and ERG staining, as indicated. Scale bars in (D) =200 µm, in (E)= 50µm and in (F, G)=10 µm. **(H)** KLF6 mRNA expression in healthy and PAH ECFCs. Bars are means ±SEM, ***P<0.001, n=4-5, Student t-test.

A total of 480 genes were detected as differentially expressed in plexiform lesions (comparison of vessels found within plexiform lesions with controls; p-value<0.05, log_2_|FC|>0.25 and <-0.25) (Figure 6B and Supplementary File S8). Remarkably, more than half of plexiform lesion DEGs (254/480 genes) were identified as KLF6-regulated in HPAECs (p= 1.1E-63) (Figure 6B and Supplementary Table S4). These KLF6-regulated plexiform lesion DEGs displayed significant enrichment in several pathways and functions related to endothelial development, wound healing, tube morphogenesis, VEGF signalling, TGF-β signalling, extracellular matrix organization, Rho GTPase signalling and actin cytoskeleton organization (Figure 6B). In non-plexiform, remodelled PAH vessels (displaying thickened intima and media) KLF 6-regulated DEGs showed significant associations with growth factor signalling and extracellular matrix organization (Supplementary Figure S14 and Supplementary File S9).

Comparison of our data with a recently published spatial transcriptomic analysis of IPAH vasculature^56^ showed that KLF6-regulated genes constituted 53% of top plexiform-specific DEGs, consistent with our findings described in this study (Supplementary Figure S15 and Supplementary Table S4).

Immunostaining of human PAH lung showed that KLF6 expression was increased predominantly in the cells lining the vascular channels in plexiform lesions and displaying endothelial identity markers ERG and vWF. These cells were also noted, albeit less frequently, in and around the remodelled vessels (Figure 6C-G and Supplementary Figures S16 and S17). A marked upregulation of KLF6 expression was also found in PAH ECFCs, compared with the corresponding healthy controls (Figure 6H). PAH blood-derived endothelial colony forming cells (ECFCs) display numerous characteristics of PAH endothelium, including increased proliferative and angiogenic responses ^57,58^ and are thought to play a contributory role in the formation of angioproliferative lesions in human PAH.

In the healthy human lung, single KLF6^+^ cells were sparsely dispersed in the lung parenchyma (Supplementary Figure S16 C) but did not exhibit any particular pattern of localization. Human placental tissue, known to express high levels of KLF6^59^, served as a positive control (Supplementary Figure S16 E). Lung tissue from a patient with tuberculosis, taken as disease control, showed increased numbers of KLF6^+^ cells compared with healthy controls (Supplementary Figure S18 C), however, in contrast to PAH lung tissue, these cells did not show a collective pattern of localization. Lung tissues from pulmonary hypertensive MCT rats and hypoxic mice did not show an accumulation of KLF6^+^ cells (Supplementary Figure S18 A, B), whilst Sugen/hypoxia rats exhibited increased number of KLF6^+^ cells in the lung (Supplementary Figure S18 A). These cells however did not display the collective organization observed in human PAH lung tissues.

Alveolar capillary dysplasia (ACD) is a rare lung developmental disorder leading to persistent pulmonary arterial hypertension and fatal outcomes in newborns^60^. Recently, Guo et al. reported an integrated analysis of single nucleus RNA-seq (snRNA-seq) of ACDMPV (n=5) and control lungs (three 3-year-old and three preterm neonate lungs) and identified six EC subtypes, including capillary 1 (CAP1) cells, capillary 2 (CAP2) cells, systemic vascular ECs (SVECs; also known as bronchial vessels), venous ECs (VECs), arterial ECs (AECs), and lymphatic ECs (LECs), in the integrated data ^33^. Given the pervasive role of KLF6 in regulating transcriptional programs essential for endothelial homeostasis, and its involvement in PAH-associated vascular remodelling, we performed a single cell comparative analysis of *KLF6* expression in ACDMPV vs. control lungs within each EC subtype using this integrated snRNA-seq dataset. The analysis revealed a disease-related increase in KLF6 expression in arterial ECs, and to a lesser extent in venous ECs, when compared to age-matched controls and only in arterial ECs when compared with preterm neonate controls (n=3/group) (Supplementary Figure S19 and Supplementary Table S5).

These results collectively demonstrate that the accumulation and structural reorganization of KLF6+ endothelial cells are characteristic of human PAH. The elevated expression of KLF6 in arterial endothelial cells in ACDMPV lungs implies a wider relevance of KLF6 upregulation in pulmonary vascular disease.

## DISCUSSION

This study identifies KLF6 as a potential driver of the pulmonary endothelial transcriptional response to injury in early PAH and as a significant feature of plexogenic arteriopathy in human disease. The proposed involvement of KLF6 is illustrated in Figure 7.

**Figure 7.**
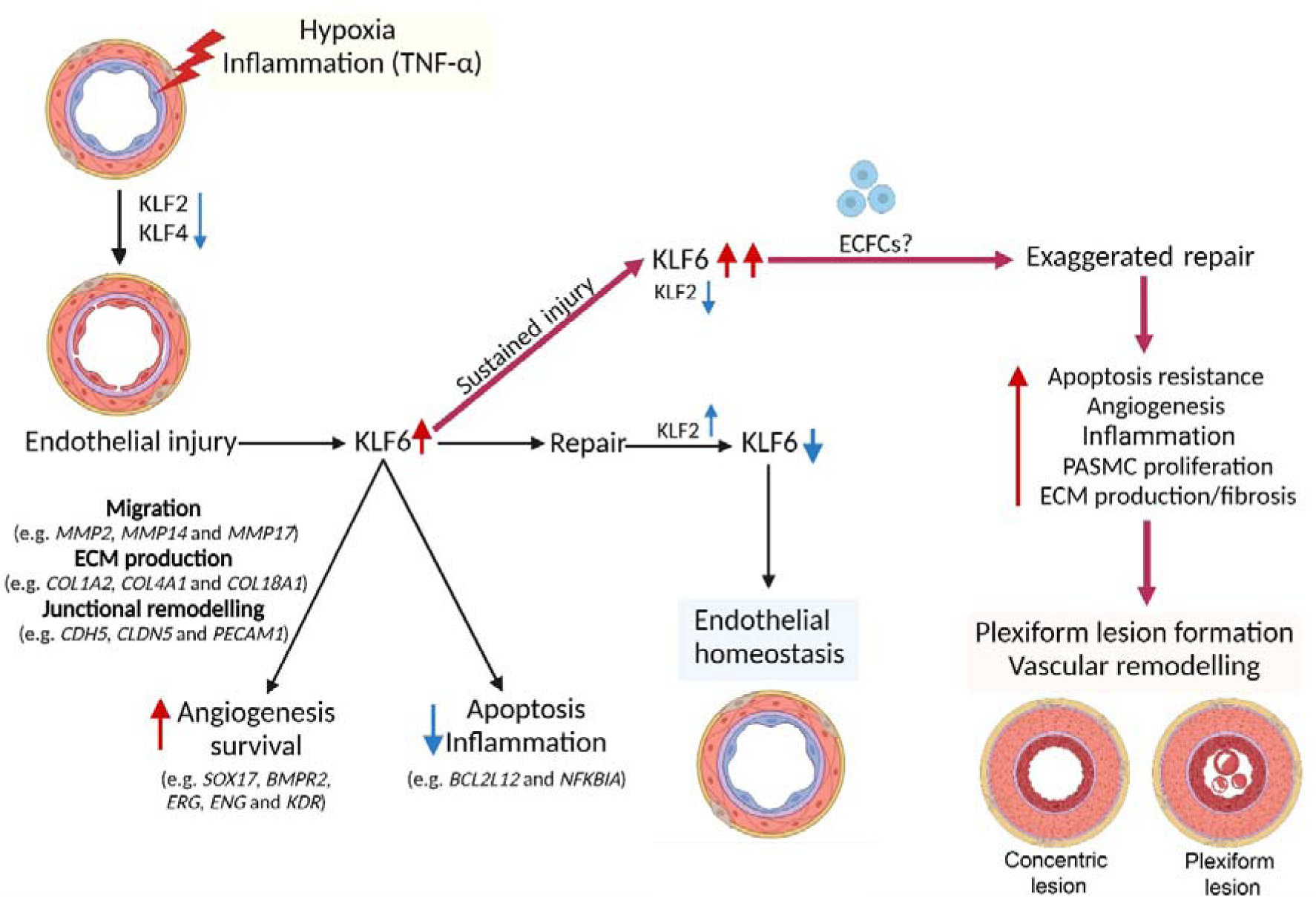
Schematic diagram of the proposed role of KLF6 in PAH. KLF6 is activated in response to endothelial injury to orchestrate the process of endothelial repair. This process is characterized by increased endothelial cell survival, angiogenesis, reduced apoptosis, and reduced inflammatory response. Under normal, healthy conditions, KLF6 activity is downregulated once the process of endothelial repair is complete. However, its continued activation under chronic hypoxic and inflammatory conditions induces endothelial apoptosis resistance, angiogenesis, fibrosis, inflammation and PASMC proliferation, augmenting vascular remodelling. Examples of KLF6-regulated genes are shown in italics. Further detail can be found in Discussion

KLF6, much like its target gene VEGF, is likely to play a dual role in PAH, offering early protective effects but contributing to pathogenic processes at later stages of the disease^61^. The pro-repair and pro-angiogenic effects of KLF6 can be linked with increased expression of arterial identity and blood vessel development genes^21,62^, such as SRY-box 17 (*SOX17*), ETS-related gene (*ERG*), VEGF receptors *FLT1* and *KDR*, angiopoietin receptor TIE2 (*TEK*), endoglin (*ENG*), activin receptor-like kinase 1 (ACVRL1), and bone morphogenetic protein receptor 2 (*BMPR2*) as well as increased expression of matrix metalloproteinases (*MMP2* and *MMP14*) and junctional proteins VE-cadherin (*CDH5*) and PECAM-1 (*PECAM1*).

DisGeNET disease enrichment analysis^29^ confirmed a significant association between KLF6 and PAH, with KLF6-regulated genes enriched in critical PAH pathways such as FOXO, p53, TNF-α, NF-kB, PI3K-Akt, and Ras signalling ^63-67^.

In addition to aiding endothelial repair, endothelial KLF6 may also increase the proliferation of PASMCs, leading to greater vascular remodelling. Supporting this idea, endothelial KLF6 has been shown to promote neointima formation in injured mouse femoral arteries^21^. We also observed increased PASMC proliferation when co-cultured with KLF6-overexpressing HPAECs. The mediators of this response still need to be identified, but the inhibitory effects of drugs used in PAH therapy, bosentan^35^ and imatinib^36^, suggest that ET-1 and PDGF-β, both downstream targets of KLF6, play a role in promoting growth in PAH PASMCs.

Spatial transcriptomic analysis of PAH lung vascular tissues has uncovered a significant enrichment of KLF6-regulated genes in processes that drive vascular repair and angiogenesis. Our data, along with recently published spatial transcriptomic analysis of IPAH vasculature^56^ show that KLF6-regulated genes account for more than half of all DEGs found in plexiform lesions, and that the expression of genes mutated in hereditary PAH is significantly upregulated. Although this may seem paradoxical, both gain and loss of function in these genes, primarily from the TGF-β family, can be associated with vascular pathology in PAH, likely reflecting the complexity of signalling events in the context of additional environmental or genetic factors^68^.

Human PAH lung sections showed accumulation of KLF6^+^ cells within vascular channels in plexiform lesions and, to a lesser extent, in other parts of the remodelled lung. These KLF6^+^ vWF^+^ ERG^+^ cells may potentially represent a subpopulation of endothelial progenitor cells, known to accumulate in lung tissues in the end-stage PAH ^57,69,70^. Consistent with this premise, we observed a significantly elevated expression of KLF6 in PAH blood-derived endothelial progenitor cells (also called the late outgrowth colony forming cells; ECFCs).

ECFCs are often used as endothelial surrogates in PAH and other cardiovascular diseases ^71^. High expression levels of KLF6 in these cells is likely to account for their pro-migratory, pro-angiogenic behaviour^72^ ^73^, potentially contributing to the formation of angio-proliferative vascular lesions in PAH lung. Selective overexpression of KLF6 in PAH endothelium is corroborated by recent data showing that only 2 out of 8 endothelial cell clusters in PAH lung display a pro-angiogenic phenotype^74^. Interestingly, amongst the animal models tested, only Sugen/hypoxia rats exhibited increased numbers of KLF6+ cells in the end stage disease, although without the collective organization, as seen in human PAH lung. It is well accepted that most rodent models poorly reproduce vascular pathology seen in human PAH, with only Sugen/hypoxia rats replicating, to a limited degree, plexiform-like arteriopathy^12^. The observed variations are likely to stem from interspecies differences in regenerative mechanisms as well as innate and adaptive immunity^75,76^.

By employing an alternative analytical approach, we also analysed single nucleus RNA sequencing data of samples from a distinct condition - alveolar capillary dysplasia characterized by severe pulmonary hypertension^77^. Through this analysis, we verified an elevation in KLF6 expression within pulmonary arterial endothelial cells. The KLF6-induced activation of VEGF/KDR signalling, prominent in both PAH and ACD^33^ lungs, may contribute to the disruption of the alveolar microvasculature, observed in both conditions.

KLF6 expression was induced concurrently with the inhibition of KLF2 and KLF4 in PAH lungs and cultured cells. Functional analyses and transcriptional profiling revealed that KLF2, KLF4, and KLF6 share anti-proliferative effects, but KLF6 uniquely stimulates endothelial repair and angiogenesis. Our results suggest that the regulatory relationship between KLF2 and KLF6 is crucial for the timely execution of endothelial repair. In the proposed scenario, the KLF6-induced increase in KLF2 expression suppresses KLF6 once endothelial repair is complete, thereby restoring endothelial homeostasis. This regulatory feedback response is unlikely to operate in PAH, where the expression and activity of KLF2 are chronically suppressed. Though we did not see a direct relationship between KLF4 and KLF6 expression, these two transcription factors are likely to interact via their transcriptional targets. The N-terminal domain of KLF6 is responsible for recruiting and interacting with different transcription factors and cofactors, including Sp1, KLF4, p53, hypoxia-inducible factor 1 alpha (HIF1α), runt related transcription factor 1 (RUNX1), E2F1, GTF3C1 and histone deacetylase 3 (HDAC3), to regulate the transcription process in a context-dependent manner^22^. Characterizing the complex crosstalk between KLF2, KLF4, and KLF6 in healthy and diseased endothelium will require further investigation.

Targeting KLF6 could represent a new therapeutic approach to mitigate the exaggerated repair processes in PAH, akin to existing strategies that pharmacologically inhibit DNA repair. ^78,79^. Whilst selective pharmacological inhibitors or activators of KLF6 have not been identified, in cancer cells KLF6 expression was regulated by a number of factors, including large HIF2α-activated and H3 acetylation-dependent super enhancer^80^, miR-191-5p, miR-200c-3p, miRNA-543 ^81-83^, long non-coding RNA CR749391^84^, transcription factor SP2 and HIF-185 86,87.

In summary, we identify KLF6 as potential important regulator of endothelial dysfunction in PAH. We show that KLF6 activation is a hallmark of apoptosis-resistant, angio-proliferative pulmonary vascular endothelial phenotype in human PAH. Dysregulation of KLF6 signalling may have broader implications for pulmonary vascular disease.

## ACKNOWLEDGEMENTS

RA received a PhD studentship from the University of Hafr Al Batin, KSA, and the Saudi Cultural Bureau in London (UKSACB). We are very grateful to Research Tissue Bank of Papworth Road, Cambridge Biomedical Campus, in particular to Marie Chet Malgapo, for providing lung sections of healthy and PAH lungs. We also thank Professor Martin Wilkins for his continuous support and helpful advice throughout the project. Mr Steve Rothery (Imperial FILM facility) for his assistance and advice with image acquisition and analysis. All widefield/confocal imaging analysis was performed at the Facility for Imaging by Light Microscopy (FILM) at Imperial College London. Human samples used in this research project were obtained from the Imperial College Healthcare Tissue Bank (ICHTB). ICHTB is supported by the National Institute for Health Research (NIHR) Biomedical Research Centre based at Imperial College Healthcare NHS Trust and Imperial College London. ICHTB is approved by Wales REC3 to release human material for research (17/WA/0161). We also thank the Imperial College London High Performance Computing Service.

## AUTHOR CONTRIBUTIONS

RA designed and performed experiments, analysed data, and wrote the manuscript; NF, HM analysed RNA-seq data, IC analysed RNA-seq data and critically evaluated manuscript; NM, NL, AL and MK analysed spatial transcriptomics data and critically evaluated manuscript, LZ and CNC provided lung tissues from pre-clinical PAH and critically evaluated manuscript, MAS provided surgical samples of pulmonary arteries, JAW and MG provided and analysed ACD snRNA-seq data patient data and critically analysed evaluated manuscript; B.W.S. conceived the study, secured funding, performed experiments and wrote the manuscript.

## DATA AVAILABILITY

The RNA-sequencing data generated and analysed during this study are currently being uploaded to European Genome-phenome Archive (EGA). All data generated or analysed during this study are included in this published article (and its supplementary information files) or are available from the corresponding author upon request.

