## Supplemental Methods for "KLF6 in Pulmonary Hypertension: The Dual Role of Friend and Foe"

Alharbi et al.

**SUPPLEMENTAL MATERIALS AND METHODS**

**Cell Culture**

Human Pulmonary Artery Endothelial Cells (HPAECs) were obtained from PromoCell (Germany, Cat. No. C-12241) at passage 3. The cells were used between passages 5-10. Information on the HPAEC donors – including age and gender population doubling time – are provided in Table M1.

**Table M1. Donor information for HPAECs used in cell culture experiments.**

| **Lot #** | **Age** | **Gender** | **Doubling Time** | **Company** |
| --- | --- | --- | --- | --- |
| 431Z031 | 23 | Female | 48.3h | PromoCell |
| 433Z034.1 | 34 | Female | 55.4h | PromoCell |
| 433Z034.2 | 34 | Female | 58.1h | PromoCell |
| 458Z016.12 | 51 | Female | 28.5h | PromoCell |
| 458Z016.13 | 51 | Female | 28.5h | PromoCell |
| 483Z021.6 | 49 | Male | 43.3h | PromoCell |
| 451Z031.14 | 68 | Male | 25.4h | PromoCell |
| 18TL155113 | 42 | Male | 25.0h | Lonza |

Cells were cultured in T75 culture flasks (Sarstedt, Germany, Cat. No. 83.3911.002) that had been pre-coated with bovine plasma fibronectin (10 µg/mL, EMD Millipore Corp, USA, Cat. No. 341631) in an Endothelial Cell Basal Medium 2 (PromoCell, Germany, Cat. No. C- 22211). This medium was supplemented with Endothelial Cell Growth Medium2 (ECGM2 SupplementPack; PromoCell, Germany, Cat. No. C-39211), 1% penicillin (100 U/mL)-streptomycin (100 μg/mL, Invitrogen, UK, Cat. No. 15140- 122) and 1% MycoZap™ Prophylactic (Lonza, Cat. No. VZA-2032). The cells were then incubated in a humidified incubator at 37 °C with 21% O_2_, 5% CO_2_, and the endothelial cell growth medium was replaced with a fresh medium every 2-3 days. Only HPAECs displaying endothelium-specific cobblestone morphology at confluence were used for experimentation. The cells were used between passages 5 and 8.

Human Pulmonary Artery Smooth Muscle Cells (HPASMCs) were obtained from Lonza (Walkersville, USA, Cat. No. CC-2581) at passage 3. Cells were cultured in T75 tissue culture flasks pre-coated with 0.2% porcine gelatin (Sigma, Cat. no. G1890) in smooth muscle cell growth medium 2 (SmGM2, PromoCell, Cat. no. C-22062), supplemented with 5% FBS and growth factor supplement (PromoCell, Cat. no. C-39267) and antibiotics. The cells were used between passages 4-6. For HPAEC and HPASMC co-culture, ECGM2 medium was supplemented with 10% foetal bovine serum (FBS). VEGF, heparin, ascorbic acid, and hydrocortisone, known to inhibit HPASMC proliferation, were excluded.

**Peripheral blood mononuclear cells (PBMCs) and endothelial colony forming cells (ECFCs).**

Venous blood samples were obtained from healthy volunteers (n=5) and from heritable PAH (HPAH) patients with rare pathogenic BMPR2 variants (n=5)^1^ with the informed written consent and approval of the local ethics committee (REC Ref 17/LO/0563). Demographic and clinical characteristics of HPAH patients and healthy volunteers are shown in Table M2.

Peripheral blood mononuclear cells (PBMCs) were isolated using a method described by Ormiston et al. 2015^2^ and peripheral blood samples were collected into EDTA coated tubes to avoid coagulation. 50 mL of blood was diluted 1:1 with PBS (Mg^2+^/Ca^2+^free), and 22 mL of the diluted blood was carefully applied over 15 mL of Ficoll-Paque Plus gradient solution (GE Healthcare, Amersham, UK, Cat. No. 17-1440-03) in a 50 mL falcon tube and then centrifuged at 400 g for 30 minutes at room temperature with the brake off. After centrifugation, the buffy coat layer containing the PBMCs was carefully collected into a new 50 mL falcon tube and then diluted 1:1 with PBS and centrifuged at 300 g for 20 minutes with the brake on. Pelleted PBMCs were then either immediately seeded in a 24*-*wells plate pre-coated with 1% gelatin or resuspended and cryopreserved in a freezing medium containing 10% dimethyl sulfoxide (DMSO; Sigma-Aldrich, UK, Cat. No. D2650) and 90% foetal bovine serum (FBS).

The cells (3-5 × 10^7^ cells in 4 mL of EGM2 medium per well) were seeded in 6 well-plates which had been pre-coated with type 1 rat tail collagen (BD Biosciences, Bedford, MA, USA). The culture medium was changed every 24 hours for the first week to remove non*-*adherent cells and every 48 hours thereafter. Colonies typically appeared between days 7 and 28 and were isolated using an 8 mm or 10 mm in diameter attaching cloning cylinder (Millipore, Watford, UK) with vacuum grease. All the cells from the inside of the cylinder were trypsinised and resuspended in an EGM2 medium and then plated onto 24-well plates which had been precoated with fresh 1% gelatin.

After reaching 70-80% confluence, the colonies were trypsinised and subcultured into 6 well plates and T75 flasks. These late outgrowth endothelial cells (ECFCs) showed stable population doubling times and a typical endothelial cobblestone morphology. The cells were immunostained with endothelial-specific markers VE*-*cadherin (CDH5), PECAM1 (CD31) and von Willebrand Factor (vWF) to confirm endothelial cell lineage.

ECFCs were cultured in endothelial cell growth medium2 (EGM2, Lonza, Cat. No. CC-3156) containing 20% FBS (HyClone; GE Healthcare, USA, Cat. No. SH30070.03), 1% penicillin (100 U/mL)-streptomycin (100μg/mL, Invitrogen, UK*,* Cat. No. 15140- 122) and growth factors (EGM-2 bullet kit, Lonza, Cat. No. CC- 4176) in T75 culture flasks pre-coated with 1*%* gelatin. They were then incubated at 37 °C, 21% O_2_, 5% CO_2_. ECFCs were used for experiments between passages 3 and 6.

**Table M2. Demographic and clinical characteristics of HPAH patients and healthy volunteers.** Data represented as median (range). mPAP=mean Pulmonary Arterial Pressure; PDE5=Phosphodiesterase type 5; ET-1R=Endothelin-1 Receptor.

|  | | Control (n=5) | HPAH (n=5) |
| --- | --- | --- | --- |
| Females | | 4/5 | 3/5 |
| Age (years) | | 29 (23- 56) | 43 (30-58) |
| Time from diagnosis (months) | | - | 84 (29-171) |
| mPAP (mmHg) | | - | 58 (58-70) |
| Six-minute walk distance (m) | | - | 366 (261-441) |
| WHO Functional Class | **I** | - | 3 |
|  | **II** | - | 1 |
| PDE5 Inhibitors | | - | 5 |
| ET-1R Antagonists | | - | 5 |
| Prostanoids | | - | 2 |
| Anticoagulants | | - | 3 |
| Statins | | - | 1 |

**Adenoviral overexpression KLF2, KLF4 and KLF6 in HPAECs**

The overexpression of most common (long) isoforms of KLF2, KLF4 and KLF6 was achieved by adenoviral gene transfer (AdKLF2-GFP (Cat. No. ADV-213187), AdKLF4-FLAG (Cat. No. ADV- 213191), AdKLF6-HA (Cat. No. ADV- 213194); Vector Biolabs (Pennsylvania, USA). Adenoviral control (AdCTRL-GFP) for AdKLF2-GFP was AdGFP (Vector Biolabs, Cat. No. 1060), while AdTet-off^3^was kind gift from Professor Stuart Yuspa (National Cancer Institute, NIH, Bethesda, USA) and was used as adenoviral control (AdCTRL) for AdKLF4 and AdKLF6.

HPAECs were cultured in 6-well cell culture plates or 8-well ibidi μ-slides (Thistle Scientific, UK, Cat. No.IB-80829). At 80% confluency, the cells were infected with AdCTRL, AdGFP, AdKLF2, AdKLF4, AdKLF6 at multiplicity of infection (MOI) of 1:100. 3 hours post infection, the EGM2 media were replaced with a fresh medium, and the cells were then incubated at 37°C, 21% O_2_, 5% CO_2_ for 24 hours to allow for target gene overexpression before further experimentation.

**KLF6 silencing**

The knockdown of KLF6 expression was performed using silencer select siRNA targeting KLF6 (siRNA-KLF6, Thermo Fisher Scientific, UK, Cat. No. s3376). Control siRNAs included a scrambled siRNA control (siRNA-CTRL Cat. No. 4390846) and a positive siRNA control targeting GAPDH (siRNA-GAPDH, Cat. No. 4390849).

HPAECs were cultured in 6-well plates in EGM2 media until confluence reached 90%. After 24 hours of incubation, 6 μL of Lipofectamine RNAiMAX reagent (Thermo Fisher Scientific, UK, Cat. No. 13778075) was diluted in a 100 μLOpti-MEM medium (Thermo Fisher Scientific, UK, Cat. No. 31985070). 2 μL(20 pmol) of the appropriate siRNA (10 μM) was diluted in a 100 μL Opti-MEM medium. Following this, 100 μLof Lipofectamine RNAiMAX/OPTIMEM mix was combined with 100 μLof the siRNA/OPTIMEM mix and the mixture was incubated at room temperature for 5 minutes. During this time, the EGM2 medium was removed from the wells, and the cells were then washed with 1 mLof phosphate buffered saline (PBS; Sigma-Aldrich, UK, Cat. No. D8537).

1 mL of OPTIMEM medium and 100 μL of the siRNA /Lipofectamine RNAiMAX/OPTIMEM mix were added to each well. After 6 hours, the cells were washed with 1 mL of PBS, and the EGM2 media (containing 10% FBS) were added to each well. After 24 hours, the medium was removed, the cells were washed with 1 mL PBS, and 1mL of reduced serum EGM2 (containing 0.2% FBS) was added to each well to synchronise the cells for maximal response to stimuli in further experiments. At 48 hours after transfection, the transfected cells were used for flow experiments.

**HPAEC culture under flow.**

HPAECs were cultured in Nunc Slide Flaskettes (Thermo Fisher Scientific, UK, Cat. No. 170920) until 95% confluent. The bottom slides with the cells were then detached from the flask and placed inside the flow chamber in a parallel flow apparatus^4^. The cells were then subjected to a laminar flow at 4 dynes/cm^2^, physiological for lung arteries^5^, for different periods (30 mins,1h, 4h, 6h and 24h). After the flow exposure, the cells were used for RNA extraction and RT-qPCR to measure mRNA expression levels of KLF2, KLF4 and KLF6.

**Real-time quantitative PCR (qPCR), RNA sequencing (RNA-seq) and spatial transcriptomics**

**RNA extraction from lung tissues**. Frozen lung tissue samples from MCT rats were cut into small pieces and placed in a BioMasher tube (Takara Bio Europe, France, Cat. No. 9791A) containing 100 μL of TRIzol Reagent (Life Technologies, UK, Cat. No. 15596026). After grinding the samples 10-15 times, 600 μL of TRIzol Reagent was added and the homogenized solution was transferred into 1.5 mL Eppendorf tubes and vortexed for ~30 seconds. 140 µLRNase-free chloroform (Thermo Fisher Scientific, UK, Cat. No. 327155000) was added and the sample was incubated at room temperature for 2-3 minutes and then placed on ice. The mixture was centrifuged at 12,000 g for 15 minutes at 4°C. The upper aqueous phase was transferred into a new Eppendorf tube. 4 µLof glycogen (5 mg/mL; Thermo Fisher Scientific, UK, Cat. No. AM9510) and 450 µL of Isopropanol (Thermo Fisher Scientific, UK, Cat. No. 327272500) were then added and inverted to mix for 10 seconds. The mixture was placed in a fridge at 4°C for 10 minutes to enhance RNA precipitation, and was then centrifuged at 12,000 g for 10 minutes at 4°C. The supernatant was aspirated, and the pellet was air dried for about 10 minutes. The RNA pellet was then resuspended in 20 µL RNase-free water. The total RNA was quantified using a NanoDrop^TM^ ND-2000 spectrophotometer (Thermo Scientific, UK).

**RNA extraction from cultured cells.** HPAECs were washed with pre-warmed PBS, then trypsin solution (Gibco, UK, Cat. No. 25200072) was added, followed by incubation at 37°C, 21% O_2_, 5% CO_2_ for ~5 minutes. Dulbecco’s Modified Eagle’s Medium (DMEM; Sigma-Aldrich, UK, Cat. No. D6429), supplemented with 10% foetal bovine serum (FBS; Thermo Fisher Scientific, UK, Cat. No. 10270098), was then added to neutralize the enzymatic reaction of the trypsin. The cell suspension was transferred to a 1.5 mLRNase-free microcentrifuge tube and centrifuged at 500 g for 5 minutes at room temperature. The supernatant was discarded, and the cell pellet was washed once with PBS and pelleted again. The cell pellet was either stored at -80 °C to be used later, or was used immediately for RNA extraction. Before starting the RNA extraction procedure, the bench surfaces, pipettes and all the equipment was sprayed with RNaseZap™ RNase Decontamination Solution (Invitrogen, USA, Cat. No. AM9780).

Total RNA was extracted using Monarch Total RNA Miniprep Kit (New England Biolabs UK, Cat. No. T2010S) according to the manufacturer’s instructions. All centrifugation steps were performed at (8,000-16,000 x g). Briefly, 300 μL of RNA Lysis Buffer containing 1% beta-mercaptoethanol (β-ME) were added to the cell pellet and the sample was vortexed vigorously for 30 seconds to lyse the cells. The mixture was then transferred to a gDNA removal column and centrifuged for 30 seconds to remove the genomic DNA. After centrifuging, the gDNA removal column was discarded and the flow-through in the collection tube which contained the RNA was saved. 300 μL of ethanol (≥ 95%) were added to 300 μL of the flow-through and mixed thoroughly by pipetting up and down. The mixture was then transferred to an RNA Purification Column and centrifuged for 30 seconds. After discarding the flow-through, 500 μL of RNA Wash Buffer were added to the RNA Purification Column and centrifuged for 30 seconds. The flow-through was then discarded, and 500 μL of RNA Priming Buffer were added and centrifuged again for 30 seconds. For the second wash, another 500 μL of RNA Wash Buffer were added and centrifuged for 1 minute. After discarding the flow-through, the RNA Purification Column was placed in a RNase-free microcentrifuge tube and the RNA was eluted in 50 μL nuclease-free water and centrifuged at maximum speed of 16,000 x g for 1 minute. The elution step was repeated twice by pipetting the eluted RNA back into the same RNA purification column and centrifuging it for 1 minute to maximize total RNA yield per sample.

RNA concentration and purity were determined using a NanoDrop^TM^ ND-2000 spectrophotometer (Thermo Fisher Scientific, UK), and the samples were stored at -80°C for later analyses. Ratio of absorbance at 260 nm and 280 nm (A260/A280) with values greater than 1.8 considered good quality RNA. For the RNA sequencing experiment, RNA quality was assessed using a TapeStation 2200 system (Agilent Technologies, UK) at the Imperial BRC Genomics Facility at Imperial College London. All samples had RNA Integrity Numbers (RIN) of 8 or higher.

**Reverse Transcription (RT).** 100 ng of the total RNA was reverse-transcribed into complementary DNA (cDNA) using a LunaScript RT SuperMix Kit (New England Biolabs, UK, Cat. No. E3010L) according to the manufacturer’s instructions. 20 μL reaction volume were prepared in 96-well PCR plates (Thermo Fisher scientific, UK, Cat. No. 4306737), as shown in Table M3. A negative No-RT control was added to each plate to verify the absence of DNA contamination in the RNA samples. Before the reverse transcription, the PCR plates were sealed, centrifuged and placed in a SimpliAmp^TM^ Thermal Cycler (Applied Biosystems, USA). Thermal conditions are shown in Table M4. cDNA samples were then diluted to 1 ng/μL using nuclease-free water and stored at -20°C.

**Table M3. Components of reverse transcription reaction mix.** Modified from New England Biolabs (<https://uk.neb.com>).

| Component | 20 μL Reaction | Final Concentration |
| --- | --- | --- |
| LunaScript RT SuperMix (5x) | 4 µL | 1X |
| RNA sample | variable | 100ng |
| Nuclease-free Water | up to 20 µL | - |

**Table M4. Thermal conditions for reverse transcription.** Modified from New England Biolabs (<https://uk.neb.com>).

| **Cycle Step** | **Temperature** | **Time** | **Cycles** |
| --- | --- | --- | --- |
| Primer Annealing | 25°C | 2 minutes | 1 |
| cDNA Synthesis | 55°C | 10 minutes | 1 |
| Heat Inactivation | 95°C | 1 minute | 1 |
| Hold | 4°C | ∞ | 1 |

**Real-time quantitative PCR (qPCR)**.

A qPCR master mix solution was prepared for each gene of interest as described in Table M5 below.

**Table M5. Components of qPCR master mix.** Modified from New England Biolabs (<https://uk.neb.com>).

| **Component** | **10 μL Reaction** | **Final Concentration** |
| --- | --- | --- |
| Luna Universal qPCR Master Mix | 5 μL | 1X |
| 10 μM forward primer | 0.5μ L | 0.25 μM |
| 10 μM reverse primer | 0.5 μL | 0.25 μM |
| Nuclease-free water | 3 μL | - |
| cDNA products | 1 μL | 1ng |

9 μL of qPCR master mix solution were added in a 384-well PCR plate (StarLab, UK, Cat. No. E1042-3840). After preparing the qPCR master mix solution, 1 ng/μL of the cDNA sample was first added in a 384-well PCR plate (StarLab, UK, Cat. No. E1042-3840), and 9 μL of qPCR master mix solution was then added to each well of the plate. A negative control was included for each gene of interest in which the cDNA was replaced by Nuclease-free water. The qPCR was carried out using a QuantStudio^TM^ 6 Flex Real-Time PCR System (Applied Biosystems, USA). Amplification steps for qPCR reactions are described in Table M6.

**Table M6. Cycling conditions for qPCR.**

| **Cycle Stage** | **Temperature** | **Time** | **Cycles** |
| --- | --- | --- | --- |
| Hold Stage | 50°C | 2 minutes | 1 |
|  | 95°C | 5 minutes |  |
| PCR Stage | 95°C | 15 seconds | 40 |
|  | 62°C | 30 seconds |  |
| Melt Curve Stage | 95°C | 15 seconds | 1 |
|  | 60°C | 1 minute |  |
|  | 95°C | 15 seconds |  |

Gene-specific primers for qRT-PCR were designed using FASTA sequences (PubMed, NCBI) and PrimerBlast (NCBI, USA), using default parameters. At least one primer in each pair spanned the exon–exon junction. All primer sequences are listed in Table M7.

The relative expression level was calculated using the 2*^–^*^∆∆CT^ method with a target gene that had been normalized to the expression of appropriate housekeeping gene, which was actin beta (*ACTB*) for static experiments or beta-2 microglobulin (*B2M*) for flow experiments.

**Table M7Primer sequences used for qPCR.**

| **Gene** | **Forward Primer (5’-3’)** | **Reverse Primer (5’-3’)** |
| --- | --- | --- |
| **Human** | | |
| ***KLF2*** | GCAAGACCTACACCAAGAGTTCG | CATGTGCCGTTTCATGTGC |
| ***KLF4*** | TGGACCCCCTCTCAGCAATG | CTCTTGGTAATGGAGCGGCG |
| ***KLF6*** | GGGTGTGGCTCTTTGCTTTA | AGCGTTAGTCACTGCTCATTTC |
| ***BMPR2*** | CTGCGGCTGCTTCGCAGAAT | TGGTGTTGTGTCAGGAGGTGG |
| ***SOX17*** | GGACCGCACGGAATTTGAAC | GGACACCACCGAGGAAATGG |
| ***ERG*** | GGAGTGGGCGGTGAAAGA | AAGGATGTCGGCGTTGTAGC |
| ***MKI67*** | TGCTCCCCACCTCAGAGAGTT | GACCTTCCTGACCTGTTTGCAG |
| ***PCNA*** | GTGAACCTCACCAGTATGTCCAA | ACTAGCGCCAAGGTATCCGC |
| ***BRD4*** | GATTGCCGGCTCCTCCAAGAT | ATGGTGCTTCTTCTGCTCCCTC |
| ***PECAM1*** | TGGAAGGAGTGCCCAGTCCCA | CGGAAGGATAAAACGCGGTCCTG |
| ***CDH5*** | CCTCCGATACATGAGCCCTCC | TCGGAAGAACTGGCCCTTGT |
| ***KDR*** | CACCAGAAATGTACCAGACCATGC | AGTCCAGAATCCTCTTCCATGCTC |
| ***HES1*** | AGAAAGATAGCTCGCGGCATT | CGGAGGTGCTTCACTGTCAT |
| ***FLT1*** | GTTTCCAGACCCGGCTCTCT | TGGTTACAGGGGTGCCAGAA |
| ***TIE2*** | CCCTGAATGCCCCAAACGTG | ACGGACCAGTTGCACACAGA |
| ***SLC2A1*** | GGGCCAAGAGTGTGCTAAAGA | GGTGACCTTCTTCTCCCGCA |
| ***ACTB1*** | GCACCACACCTTCTACAATGA | GTCATCTTCTCGCGGTTGGC |
| ***B2M*** | CCAGCGTACTCCAAAGATTCAGG | TCAATGTCGGATGGATGAAACCC |
| **Rat** | | |
| ***ACTB*** | ACCCCAGCCATGTACGTAGC | TGCCTGGGTACATGGTGGTG |
| ***KLF2*** | ACTTGCAGCTACACCAACTG | CTGTGACCCGTGTGCTTG |
| ***KLF4*** | TGAACTGACCAGGCACTACC | GCCTCTTCATGTGTAAGGCA |
| ***KLF6*** | CGGACGCACACAGGAGAAAA | CGGTGTGCTTTCGGAAGTG |
| **Mouse** | | |
| ***B2M*** | GCCTGTATGCTATCCAGAAAACC | TCAATGTGAGGCGGGTGGAA |
| ***KLF2*** | GAGCCTATCTTGCCGTCCTTT | CACGTTGTTTAGGTCCTCATCC |
| ***KLF4*** | CCAGAGGAGCCCAAGCCAAAG | CGGTAGTGCCTGGTCAGTTCAT |
| ***KLF6*** | ACTGTCTTTTCCAACCCGAC | AAGATAGCGTTCCAACTCCAG |

**RNA Sequencing**

RNA sequencing (RNAseq) (75bp paired-end reads) was performed in an Illumina NextSeq 2000 sequencer (Illumina, USA) at the Imperial BRC Genomics Facility (Imperial College London) to assess the whole transcriptomic effects of KLF2, KLF4, and KLF6 overexpression by adenoviral gene transfer (AdKLF2, AdKLF4 and AdKLF6 conditions, respectively).

The quality of the paired end reads within the fastq files were assessed using FastQC (<https://www.bioinformatics.babraham.ac.uk/projects/fastqc/>) and MultiQC. To determine transcript abundance values, Salmon (v1.5.2) was used to map sequencing reads in the fastq files to the human genome (GENCODE v44) (<https://www.gencodegenes.org/human/>) and to obtain gene counts. To identify differentially expressed genes (DEGs), the gene expressions of the AdKLF2, AdKLF4 and AdKLF6 conditions (n=4 biological replicates/condition) were compared with the corresponding negative control conditions using the R package DESeq2 (v1.38.3)^6^, with default settings. Enhanced volcano (v1.16.0) (https://github.com/kevinblighe/EnhancedVolcano 2020) and heatmaps were generated using the R package pheatmap (v1.0.12). Gene annotation was performed using biomaRt (2.46.3). The list of PAH genes was derived from ^7^.

**Gene ontology (GO) and Kyoto Encyclopaedia of Genes and Genomes (KEGG) enrichment analysis**

GO and KEGG pathway enrichment analyses were carried out separately on the sets of up- and down-regulated genes from each comparison using the Metascape (<https://metascape.org>) online tool^8^, the WEB-based GEne SeT AnaLysis Toolkit (WebGestalt) (<https://www.webgestalt.org>)^9^ and the clusterProfiler package (v3.14.3) in R. The DisGeNET enrichment analysis and network of enriched terms of up- and down-regulated genes was performed using Metascape. Genes that passed the thresholds of FDR <0.01, absolute value of log_2_ (fold change)>2 were used for GO and KEGG pathway enrichment analyses, unless indicated otherwise.

**Comparative analysis with published RNA-seq datasets and disease-gene databases.**

To investigate whether differentially expressed genes from KLF6-overexpressing HPAECs were enriched in PAH-related genes, a comparative analysis was carried out using RNA-seq datasets from HPAECs and ECFCs from the “double hit” microfluidic model of PAH^1^ and from PAECs that had been isolated from IPAH patients^10^ or with known PAH and IPAH-related genes from DisGeNET databases (<https://www.semanticscholar.org/paper/GeneOverlap%3A-An-R-package-to-test-and-visualize-Shen/117e12840af966176bbc348db6edf034b0ea479c>). Overlapping genes were visualized as UpSet plots using the ComplexUpset package (v1.3.3) in R. The significance of the overlaps between gene sets in comparison to the genomic background was calculated by one-tailed Fisher’s exact test using the GeneOverlap package (v.1.22.0) in R (<https://www.semanticscholar.org/paper/GeneOverlap%3A-An-R-package-to-test-and-visualize-Shen/117e12840af966176bbc348db6edf034b0ea479c>)

**Spatial transcriptomics**

To assess the transcriptomic changes in PAH lungs, spatial transcriptomics of lung tissues from healthy controls and PAH patients were carried out using NanoString GeoMx® to enable high-plex spatial profiling of RNA or protein within specific areas of interest in the tissue ^11^

Formalin fixed and paraffin embedded (FFPE) lung tissue sections (3-4μm) from PAH patients (n = 6) and age- and sex-matched healthy volunteers (n = 5) were obtained from the Royal Papworth Hospital NHS Foundation Trust Tissue Bank (Cambridge, UK) with informed written consent and ethical approval by (ICHTB HTA licence: 12275; REC Wales approval: 22/WA/0214). Donor information for FFPE lung tissue samples is shown in Table M8.

**Table M8 List of human lung tissue donors used in this study.** The first control (a 49-year-old female) was excluded due to the abnormal appearance of the lung tissue.

| **Pair** | **Category** | **Age** | **Sex** |
| --- | --- | --- | --- |
| 1 | Control | 49 | Female |
|  | PAH | 32 | Female |
| 2 | Control | 53 | Female |
|  | PAH | 51 | Female |
| 3 | Control | 60 | Female |
|  | PAH | 57 | Female |
| 4 | Control | 21 | Male |
|  | PAH | 27 | Male |
| 5 | Control | 58 | Male |
|  | PAH | 37 | Male |
| 6 | Control | 64 | Male |
|  | PAH | 61 | Male |

.

The lung tissue slides were first hybridized overnight at 37°C with GeoMx Human Whole Transcriptome Atlas Human RNA probe mix (GeoMx Hu WTA, NanoString Technologies, USA). After incubation with the RNA probe mix, stringent washes were performed using a mixture of an equal volume of 100% (v/v) formamide and 4X saline-sodium citrate (SSC) to remove off-target probes and the slides were then blocked with 200 μL Buffer W (NanoString Technologies, USA) and incubated for 30 minutes at room temperature in a humidity chamber. The slides were stained with SYTO13 (nuclear marker), vWF (von Willebrand Factor; endothelial cell marker), and ACTA2 (Alpha-smooth muscle actin or α-SMA; smooth muscle cell marker) to allow the identification of blood vessels during the ROI selection. The slides were then washed with 2x SSC for 5 minutes twice and immediately loaded into the GeoMx Digital Spatial Profiler (DSP) instrument for scanning (x20 magnification) and ROI selection. For each lung tissue section, a total of 30 regions of interest (ROIs) were selected with the “circle” or “freehand” selection tool available on the GeoMx DSP software to profile blood vessels within the lungs, specifically focusing on plexiform lesions observed in PAH patients. After ROI selection on the GeoMx DSP instrument, the DSP barcodes in ROIs were ultraviolet (UV)-cleaved and collected into a 96-well DSP collection plate before being sequenced (paired-end reads) on an Illumina sequencer (Illumina, USA) at the Imperial BRC Genomics Facility (Imperial College London).

**Immunocytochemistry (ICC)**

Cells were cultured in ibidi μ-slide 8 well chamber slides (Thistle Scientific, UK, Cat. No. IB-80806) or on plastic coverslips (13mm diameter, Nunc Thermanox, USA, Cat. No. 174950). They were then washed once with pre*-*warmed PBS and fixed with 4% paraformaldehyde (w/v) (Sigma-Aldrich, UK, Cat. No. P6148) in PBS for 15 minutes at room temperature. After fixation, the cells were washed twice with PBS and permeabilized using 0.1% TritonX-100 (v/v) (Sigma-Aldrich, UK, Cat. No. 9002-93-1) in PBS. They were then gently washed twice with PBS.

For immunostaining of nuclear KLF6, PBS was removed, and the cells were incubated in 100% methanol for 5 minutes at -20°C, then air dried for 1 minute. The non-specific binding sites were then blocked by incubating cells with 2% bovine serum albumin (w/v) (BSA; Sigma-Aldrich, UK, Cat. No. A2058) in PBS for 15 minutes at room temperature, followed by incubation with primary antibody overnight at 4 °C, then secondary antibody for 2 hours at room temperature*.* Primary and secondary antibodies were both diluted in 0.1% BSA (v/v) in PBS. After each incubation time with the antibody, the cells were washed three times for 5 minutes in PBS. All primary and secondary antibodies used are listed in Table M10. Samples were then mounted in Vectashield  Antifade Mounting medium containing nuclear stain DAPI (Vector Laboratories, Cat. No. H-1200).

Immunofluorescence images were observed using a STELLARIS 8 inverted confocal microscope *(*Leica Microsystems*,* Mannheim*,* Germany) at 10x or 20x objectives.

**Immunohistochemistry (IHC)**

Immunohistochemical staining was carried out as described^13^, with some modifications. Briefly, formalin fixed and paraffin embedded lung tissue sections of healthy control (n = 6) and PAH patients (n = 6) (listed in Table M10) (Royal Papworth Hospital NHS Foundation Trust Tissue Bank, Cambridge, UK), sections of human placenta and lung tissue from a patient with pulmonary tuberculosis (Imperial College Healthcare Tissue Bank) were dewaxed in Histo-Clear (National Diagnostics, USA, Cat. No. HS-202) twice for 5 minutes, before being rehydrated in graded ethanol solutions (100%, 70% and 50%) (v/v) in distilled water (VWR, UK, Cat. No. 20821.330) for 5 minutes per wash, and then washed twice in distilled water for 5 minutes per wash. The sections were then subjected to antigen retrieval by boiling them at 95°C for 10 minutes (2 x 5 minutes on a hot plate to avoid overboiling) in a solution containing 40 mL of Low pH IHC Antigen Retrieval Solution (v/v) (Life Technologies, UK, Cat. No.00-4955-58) and 360 mL of distilled water. The sections were then washed in PBS three times for 5 minutes per wash. After air-drying, the tissue sections were encircled with a Hydrophobic Barrier PAP Pen (Thermo Scientific, UK, Cat. No. R3777). The tissue sections were then blocked with 5% bovine serum albumin (BSA) (v/v) in PBS with 4 drops of 2.5% Normal Horse Serum (Vector Laboratories, USA, Cat. No. S-2012) added and incubated for 1 hour at room temperature in a humidified chamber. After the removal of the blocking solution, sections were incubated overnight with primary antibodies diluted in 5% BSA in PBS in a humidified chamber at 4 °C before being washed with PBS three times for 5 minutes per wash. The sections were then incubated with secondary antibodies diluted in 5% BSA in PBS for 2 hours at room temperature in a humidified chamber before being washed with PBS three times for 5 minutes per wash. The primary and secondary antibodies used are listed in Table M12. The slides were then mounted in Vectashield  Antifade Mounting medium with nuclear stain DAPI. Immunofluorescence images of lung tissues were acquired using a STELLARIS 8 inverted confocal microscope (Leica Microsystems, Mannheim, Germany).

**Table M10. List of primary** **and** **secondary antibodies used** for **immunofluorescence staining.** vWF, von Willebrand factor; α-SMA and α-Smooth Muscle Actin. In addition to primary and secondary antibodies, phalloidin-TRITC (Bio-Techne, UK, Cat. No. 5783) was used as an affinity-stain for filamentous actin (F-actin) in endothelial cells.

| **Antibody Name** | **Species** | **Type** | **Dilution** | **Cat No.** | **Company** |
| --- | --- | --- | --- | --- | --- |
| KLF2  (Anti-Human) | Mouse | Monoclonal IgG | 1:200 | MAB5466 | R&D Systems |
| KLF4/GKLF  (Anti-Human) | Mouse | Monoclonal IgG | 1:100 | sc-393462 | Santa Cruz Biotechnology |
| KLF6  (Anti-Human) | Mouse | Monoclonal IgG | 1:100 | sc-365633 | Santa Cruz Biotechnology |
| KLF6  (Anti-Human) | Rabbit | Polyclonal IgG | 1:100 | PA5-84819 | Invitrogen |
| ERG  (Anti-Human) | Mouse | Monoclonal IgG | 1:200 | sc-376293 | Santa Cruz Biotechnology |
| VE-Cadherin/CD144 (Alexa Fluor 488)  (Anti-Human) | Mouse | Monoclonal IgG | 1:200 | 53-1449-42 | Invitrogen |
| α-SMA  (Cy3-Conjugated)  (Anti-Human) | Mouse | Monoclonal IgG | 1:300 | C6198 | Sigma-Aldrich |
| CD31/PECAM1  (Anti-Human) | Mouse | Monoclonal IgG | 1:500 | M0823 | Dako |
| HA-Tag  (Anti-Human) | Rabbit | Monoclonal IgG | 1:300 | C29F4 | Cell Signaling Technology |
| FLAG  (Anti-Human) | Rabbit | Polyclonal IgG | 1:300 | F7425 | Sigma-Aldrich |
| vWF  (Anti-Human) | Rabbit | Polyclonal IgG | 1:200 | A0082 | Dako |
| Cy5-Conjugated  (anti-Mouse) | Goat | Polyclonal IgG | 1:200 | 16855-AAT | AAT Bioquest |
| Alexa Fluor Plus 594 (Anti-Mouse) | Goat | Polyclonal IgG | 1:200 | A32742 | Invitrogen |

**NF-κB Luciferase Reporter Assay**

HPAECs that had been cultured in 96-well plates or in 6 channel μ-Slides (Thistle Scientific, UK, Cat. No. IB-80806) were left untreated or were infected with adenoviral NF-κB luciferase reporter^13^ (AdNFkB-luc, Vector Biolabs, Cat. No.1740), AdCTRL and AdKLF6 or were transfected with siRNA-KLF6 and scrambled-siRNA control, as required.

Since the basal KLF6 expression in unstimulated, quiescent HPAECs was insufficient to produce a measurable effect, the effects of KLF6 silencing were studied in cells cultured under flow.

3 hours after adenoviral infection or 48 hours after siRNA exposure, the culture media were replaced with a fresh EGM2 medium with or without 10 ng/mL of TNF-α (R&D Systems, USA, Cat. No. 210-TA-020). The cells were then incubated under either normoxic or hypoxic conditions for 24 hours. A luciferase reporter assay (Promega, USA, Cat. No. E1500) was then performed according to the manufacturer’s instructions. Briefly, 20 μLof cell lysates were combined with 100 μL of luciferase assay reagent in 96-microwell white opaque polystyrene plates (Thermo Scientific, UK, Cat. No. 136101). The intensity of luminescence, proportional to the level of NF-κB–driven luciferase activity, was measured in GloMax® luminometer (Promega, UK).

**Transwell permeability assay.**

Endothelial permeability assay was carried out using a 6.5 mm transwell plates with sterile 0.4 μm pore size polycarbonate membrane inserts (Corning, USA, Cat. No. 3413).

Cells were seeded at a density 2 x10^4^ cells per insert with 200 µLof cell culture medium in the apical compartments of transwell inserts that had been pre-coated with bovine plasma fibronectin (10 μg/mL, EMD Millipore Corp, USA, Cat. No. 341631). 500 μLof the medium was then added to the basal compartments of the transwell. Once the cells formed a confluent monolayer, they were infected with either AdCTRL or AdKLF6. After 2 hours of adenoviral infection, the media were changed and the cells were either left untreated or were treated with 10 ng/mL of TNF-α (R&D Systems, USA, Cat. No. 210-TA-020). The cells were then placed in normoxia (21% O_2_) or hypoxia (2% O_2_) at 37°C and 5% CO_2_ for 24 hours. After this period*,* the cells were incubated in fresh medium containing 1 mg/mL of 40 kDa FITC-Dextran (Sigma-Aldrich, Dorset, UK, Cat. No. FD40S) for 1 hour. After an hour of incubation, 500 µLof the medium was collected from the basal compartment, and the fluorescence intensity of FITC- dextran was measured at excitation/emission 490/525 nm using a GloMax® luminometer (Promega, UK).

To test thrombin-induced cell permeability, cells were infected with either AdCTRL or AdKLF6. After 24 hours, they were left untreated or stimulated with 1 U/mL of thrombin (Sigma-Aldrich, Dorset, UK, Cat. No. T7513) in a serum-free medium containing 1 mg/mL FITC-Dextran for 1 hour, then the rest of experiment was carried out as described above.

**Angiogenesis assays**.

Angiogenesis was assessed in a tube formation assay *in vitro* and in an *ex vivo* pulmonary arterial explants sprouting assay.

For the tube formation assay, HPAECs were infected with either AdCTRL or AdKLF6. 24h after infection, the cells were seeded into a 96-well plate pre-coated with growth factor-reduced Matrigel (Scientific Laboratory Supplies, UK, Cat. No. 354230). The cells were then incubated for 20 hours under normoxic conditions*.* After 20 hours of incubation*,* endothelial tube formation was imaged using a phase*-*contrast microscope. Images were then analysed using ImageJ software (Fiji, version 2.14.0) with an Angiogenesis Analyzer plugin to quantify the number of nodes and meshes as well as the total tube length.

For the arterial explants sprouting assay, surgical specimens of human pulmonary arteries were cleaned of blood and attached connective tissues in PBS. Donor characteristics – including age, gender, height, weight and diagnosis – are summarized in Table M11. The arteries were cut longitudinally with a scalpel, opened and pinned down on the dissection dish with the endothelium facing up. 50-100 μL of EGM2 containing AdCTRL or AdKLF6 was added on top of the endothelial layer, and the explants were then incubated for 3 hours in a humidified incubator. Afterwards, the arterial tissue was thoroughly washed in PBS and cut into roughly 1 mm^2^ fragments which were then embedded in Matrigel and incubated for 3 weeks in a humidified incubator under normoxic conditions. AdKLF6 was added to the explants again at 2 weeks to maintain the efficiency of gene transduction. Endothelial sprouts were fluorescently labelled with Vybrant™ CFDA SE Cell Tracer (Invitrogen, USA, Cat. No. V12883) before being imaged under a fluorescent microscope. The total area of sprouting was measured using ImageJ software (Fiji, version 2.14.0).

**Table M11. Pulmonary artery donor characteristics.** ADC: adenocarcinoma, AVM: arteriovenous malformation.

| **ID** | **Age** | **Gender** | **Height (cm)** | **Weight (kg)** | **Diagnosis** |
| --- | --- | --- | --- | --- | --- |
| CX22000050 | 56 | M | 170 | 79.8 | Pulmonary AVM |
| CX22000077 | 33 | F | 160 | 88.9 | Pulmonary AVM |
| CX22000115 | 70 | F | 158 | 53 | Non-mucinous lung ADC |
| CX22000116 | 21 | F | 157 | 45.7 | Pulmonary AVM |
| CX22000221 | 72 | M | 181 | 76 | Non-mucinous lung ADC |

**EdU Cell Proliferation Assay**

Cell proliferation assay was performed using an EdU Cell Proliferation Assay Kit (EdU-594, EMD Millipore Corp, USA, Cat. No. 17-10527), according to the manufacturer’s instructions. EdU solution was diluted 1:1000 in cell culture medium. 24 hours later cells were fixed with 3.7% formaldehyde in PBS for 15 minutes and permeabilized using 0.5% Triton X-100 in PBS for 20 minutes. The cells were then treated with an EdU reaction cocktail which was prepared according to the manufacturer’s instructions and were then incubated for 30 minutes.

Cells were mounted with Vectashield Antifade Mounting medium containing DAPI (Vector Laboratories, H-1200). Images of EdU positive cells were acquired under a fluorescent Zeiss Axio Observer widefield microscope (Carl Zeiss AG, Germany) at a 10x magnification and analysed with ImageJ software (Fiji, version 2.14.0). The results were quantified as a ratio of the number of EdU positive cells against the total number of cells.

**Proliferation of PASMCs in co-culture with HPAECs**.

Control (AdCTRL) and AdKLF6-overexpressing HPAECs were grown to confluence in 6.5 mm Transwell inserts with 1.0 µm pore size transparent PET membrane (Corning, Cat. No 353104) in EGM2 culture media containing serum and growth factors. HPASMCs were seeded at the bottom of 24-well plates (50 x10^3^ cells/well) or on the other side of empty Transwell insterts or inserts containing endothelial cells (20 x 10^3^ cells/insert) in EGM2 medium containing 5% FBS and no growth factors. Following 2-hour incubation in a humidified culture incubator, unattached PASMCs were washed away with PBS and Transwell inserts containing HPAECs or co-cultures of HPAECs and HPASMCs were inserted into the wells of 24 well plate. Culture media were replaced with EGM2 medium containing 0.2% FBS, antibiotics and EdU (for further detail please see EdU Cell Proliferation Assay), with no growth factors. PASMCs cultured in full PASMC medium served as a positive control.

**Wound healing assay**.

Endothelial cell migration was assessed in a wound healing assay using ibidi Culture-Insert 2 Well in µ-Dish 35 mm (ibidi, Germany, Cat. No. 81176) or in a scratch assay under flow. Cells were seeded at a density of 3×10^4^ cells per well. After overnight incubation, the cells were infected with either AdCTRL or AdKLF6. Next day, cells were serum- and growth factor starved (0.2% FBS, with no growth factors) for 6 hours*.* Culture inserts were removed after starvation, and the cells were washed with PBS to remove non-adherent cells. They were then either left in a starvation medium (0.2% FBS, with no growth factors) or were stimulated by the addition of full EGM2 containing serum and growth factors (EGM2, 2 mL) to the dish. Images of the cells were captured at 0 hour and 20 hours using a Rebel phase contrast microscope (ECHO, USA) with a 10x objective.

In the scratch assay, HPAECs were seeded in Nunc Lab-Tek Flaskettes (Thermo Fisher Scientific, UK, Cat. No. 170920) at a density of 2×10^5^ cells per Flaskette. After an overnight incubation, the cells were transfected with KLF6 siRNA or negative control siRNA. After 48 hours, the confluent endothelial cell monolayers were serum- and growth factor starved (0.2% FBS, with no growth factors) for 6 hours. The bottom slides containing the cells were then detached and placed into petri dishes (150 x15 mm standard style, VWR International Ltd, UK, Cat. No. 391-2003). A straight “wound” was created by scratching endothelial monolayers with a sterile 200 µL pipette tip. The slide was then washed in PBS and placed inside the flow chamber of a parallel flow apparatus, where the cells were exposed to laminar flow (4 dynes/cm^2^) for 20 hours. Images of the wound were taken at 0 hour and 20 hours using a phase contrast microscope (ECHO Rebel, USA) with a 10x objective.

The wound area was measured using a custom-made Wound Healing Tool ImageJ macro (written by Stephen Rothery, Facility for Imaging by Light Microscopy (FILM), Imperial College London)

.

**Apoptosis Caspase-Glo 3/7 Assay**

Apoptosis was assessed by measuring caspase 3/7 activity using a Caspase-Glo 3/7 Assay Kit (Promega, Southampton, UK, Cat. No. G8091), according to the manufacturer’s instructions. Briefly, following the cell treatments, Caspase-Glo 3/7 reagent was prepared by adding a Caspase-Glo 3/7 substrate to buffer and allowing them to equilibrate to room temperature. An equal volume of reagent (100 μL) was added to each well of the 96-well plate containing 100 μL of blank (medium only) and treated cells in a culture medium.

The plate was then placed on a plate shaker at 300-500 rpm for 30 seconds and incubated at room temperature for 1 hour. Luminescence was measured using a GloMax® luminometer (Promega, UK), according to the manufacturer’s instructions.

**TUNEL Apoptosis Assay**

TUNEL Assay was performed using a Click-iT Plus for In Situ Apoptosis Detection Kit with Alexa Fluor 647 dye (Invitrogen*,* Cat. No. C10619), according to the manufacturer’s instructions. TUNEL positive cells were identified using a fluorescent Zeiss Axio Observer widefield microscope (Carl Zeiss AG, Germany) with a 10x objective, and analysed with ImageJ software (Fiji, version 2.14.0). Results were quantified as a ratio of TUNEL positive cells against the total number of cells.

**Study approval**

Venous blood samples were obtained with the informed written consent and approval of the local ethics committee (REC Ref 17/LO/0563) from healthy volunteers and from HPAH patients with a rare pathogenic variant of *BMPR2* gene*.*

Human artery tissue samples were obtained following ethical approval from the Imperial College Healthcare Tissue Bank (ICHTB HTA licence: 12275; REC approval: 17/WA/0161).

Formalin fixed and paraffin embedded lung tissue sections from healthy volunteers and PAH patients were obtained from the Royal Papworth Hospital NHS Foundation Trust Tissue Bank (Cambridge, UK) with informed written consent and ethical approval by (ICHTB HTA licence: 12275; REC Wales approval: 22/WA/0214).

**Statistical Analysis**

All graphs and statistical analyses were performed using either GraphPad Prism software 9 (GraphPad Software, USA) or Rstudio (RStudio Inc, version 1.2.5042).

All experiments were performed in at least four biological replicates, with 3 technical replicates performed per experiment, unless stated otherwise. All data were tested for normal distribution using the Shapiro-Wilk test. An unpaired student’s t test was used to analyse the normally distributed data from two sample groups, while a one- or two-way ANOVA test was used to analyse three or more sample groups as appropriate.

Statistical significance was accepted when p*-*values were less than 0.05. All error bars are representative of mean (*±* SEM).
