## Supplemental Figures and Tables for "KLF6 in Pulmonary Hypertension: The Dual Role of Friend and Foe"

Alharbi et al.

SUPPLEMENTAL FIGURES

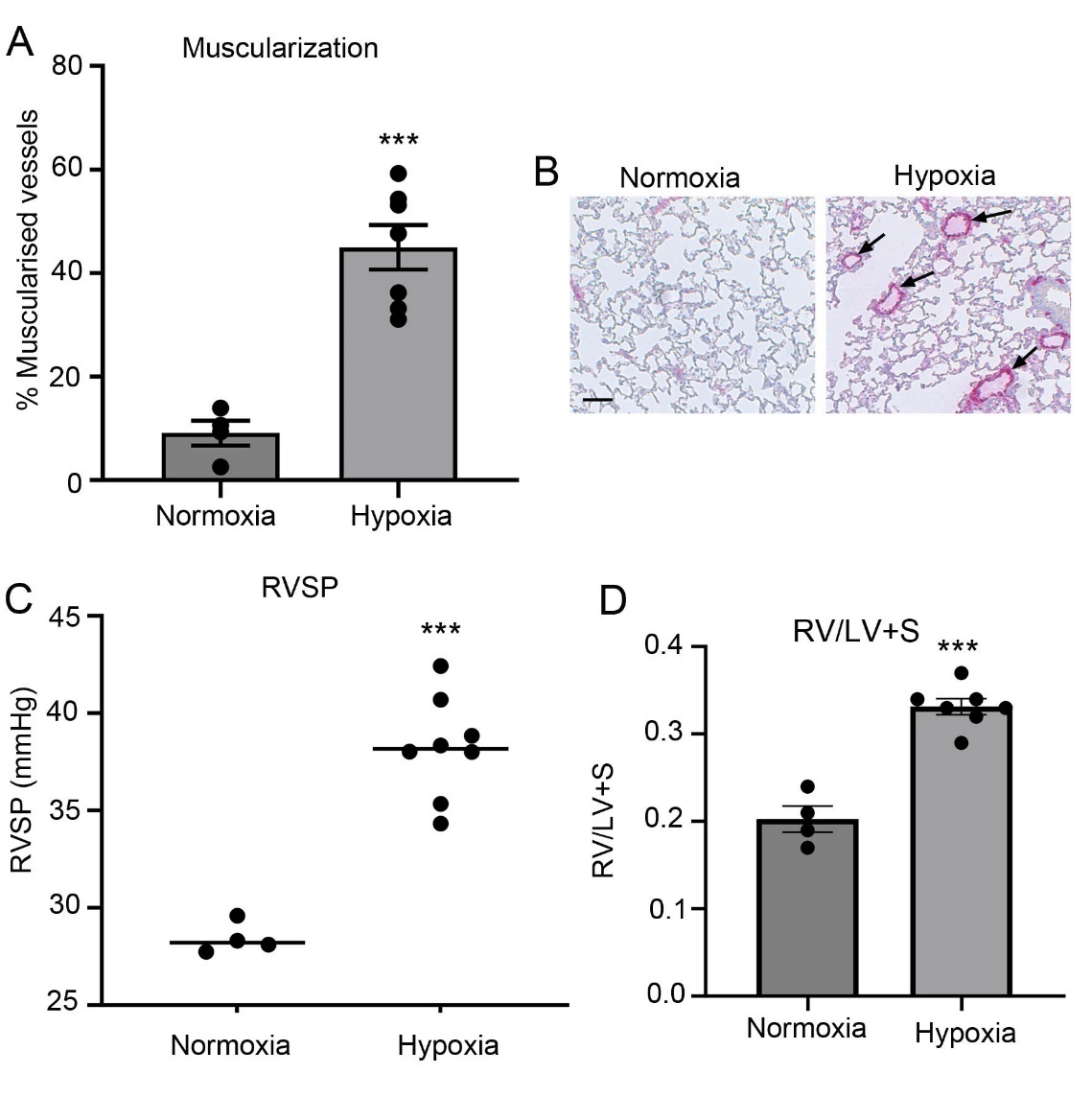

***Figure S1.*** ***Hypoxia-induced PH in mice.***

*C57/BL mice were left in normoxia (Nx) or were placed under hypoxia (Hx; 10% O_2_) for 2 weeks.* ***(A)*** *Muscularization expressed as % of α-SMA-positive vessels compared with the total number of vessels with diameter <25µm.* ***(B)*** *Representative images of lung sections from different study groups, as indicated. Alkaline phosphatase immunostaining, α-SMA is shown in pink. Arrows point to the remodelled vessels. Bar=40µm (****C****) RVSP; (****D****) RV/LV+S; ***P<0.001, comparison with normoxic controls, Student t-test; n=4 in Nx and n=8 in Hx.*

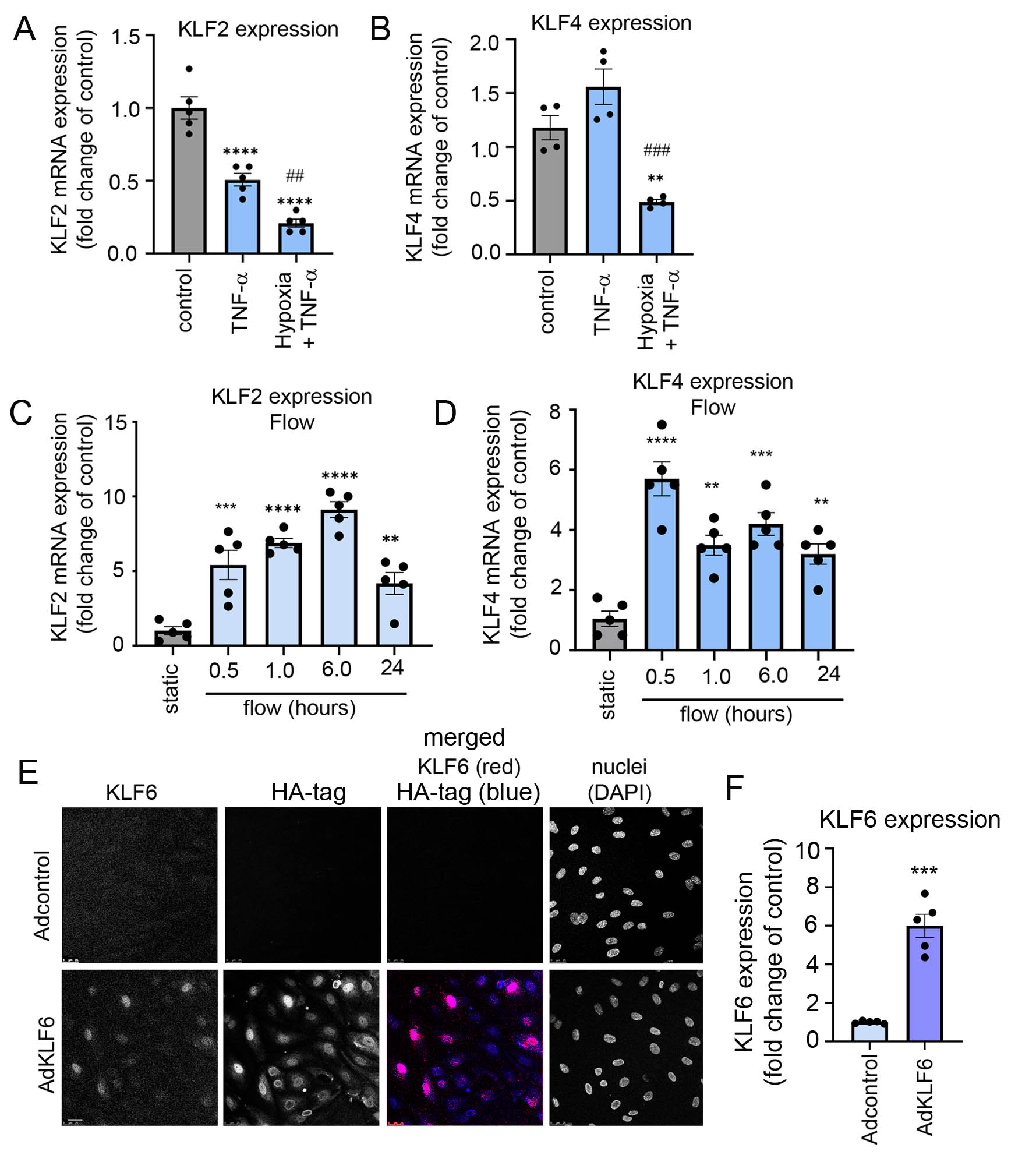

***Figure S2. KLF2, KLF4 and KLF6 overexpression in HPAECs under different experimental conditions.*** ***(A)*** *KLF2 and* ***(B)*** *KLF4 mRNA expression in HPAECs exposed to TNF-α (10 ng/ml, 24h) under normoxic or hypoxic (2% O_2_, 24h) conditions, as indicated.* ***(C)*** *and* ***(D)*** *show KLF2 and KLF4 expression in HPAECs exposed to flow (4 dynes/cm2 0-24h), respectively.* ***(E)*** *Representative confocal microscopy images showing nuclear localization of recombinant HA-tagged KLF6 (AdKLF6) in HPAECs 24 hours post-infection. Scale bar =10μm; n=3.* ***(F)*** *KLF6 protein and* ***(G)*** *KLF6 mRNA levels in HPAECs infected with AdCTRL or AdKLF6. **P<0.01, ***P<0.001, ****P<0.0001, comparison with relevant study controls. ^##^P<0.05 and ^###^P<0.001, comparisons between TNFα and TNFα+hypoxia, as indicated. In (A-D) one-way ANOVA with Tukey post-test, n=5; In* ***(F, G)*** *unpaired t-test; n=3 in (F) and n=5 in (G). Bars are means ±SEM*

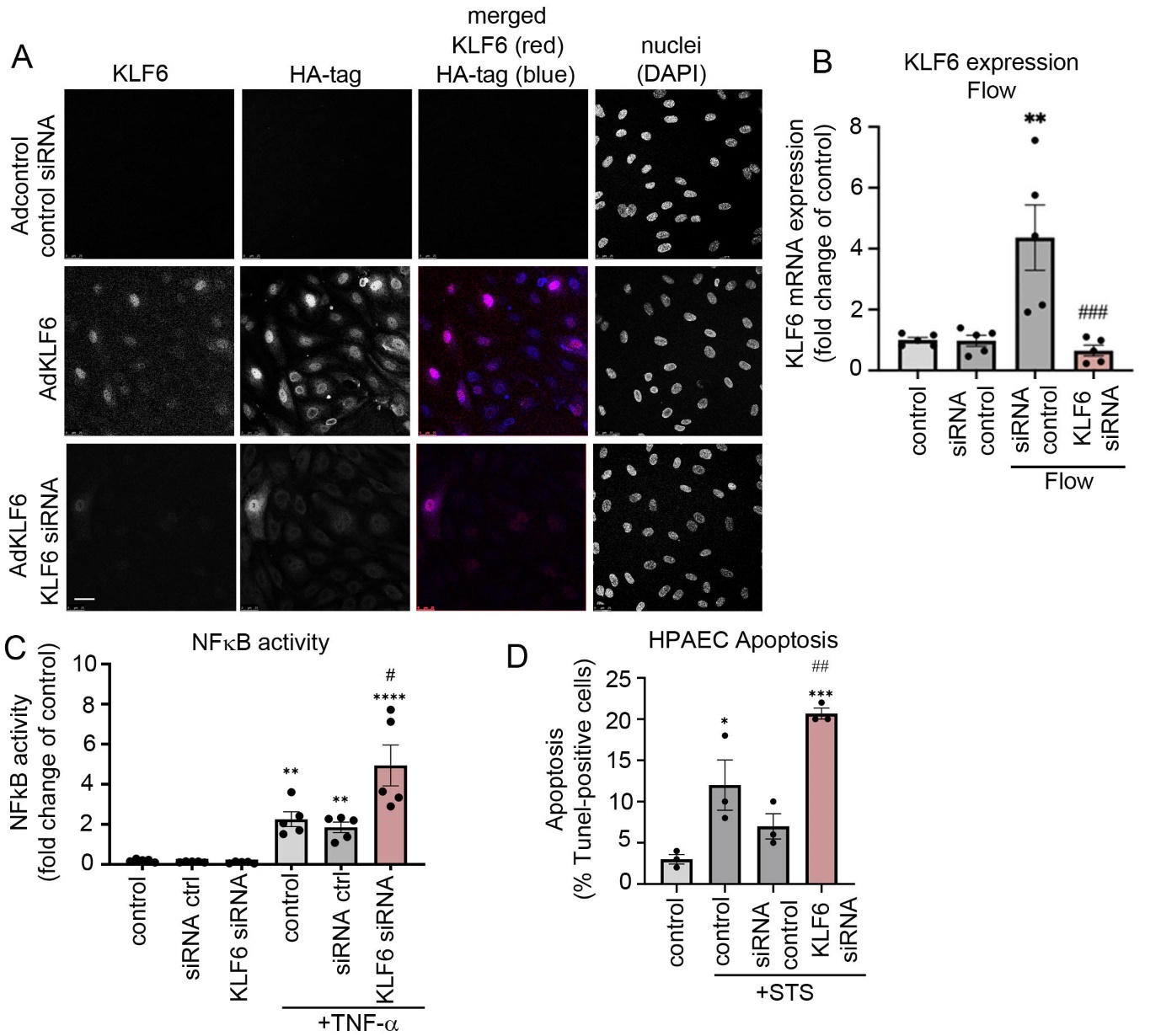

***Figure S3. Effects of KLF6 silencing in HPAECs. (A)*** *Representative confocal microscopy images showing localization of endogenous and recombinant HA-tagged KLF6 (AdKLF6) in HPAECs transfected with control, non-targeting siRNA or KLF6 siRNA, as indicated. HPAECs were transfected with siRNA 6 hours post-adenoviral infection and incubated for further 48 hours before immunostaining. Scale bar=10μm.* ***(B)*** *KLF6 mRNA levels in HPAECs transfected with control siRNA or KLF6 siRNA and exposed to flow (4 dynes/cm^2^; 24h).* ***(C)*** *NFκB activity (luciferase reporter assay) and* ***(D)*** *apoptosis in HPAECs transfected with control or KLF6 siRNA and treated with TNF-α (10 ng/ml, 24h) or staurosporine (STS), as indicated. *P<0.05, **P<0.01, ***P<0.001, ****P<0.0001, comparisons with controls; ^#^P<0.05, ^##^P<0.01, ^###^P<0.001, comparison with relevant treatment controls; one-way ANOVA with Tukey post-test. Bars are means ±SEM. In* ***(B, C)*** *n=5, in* ***(D)*** *n=4.*

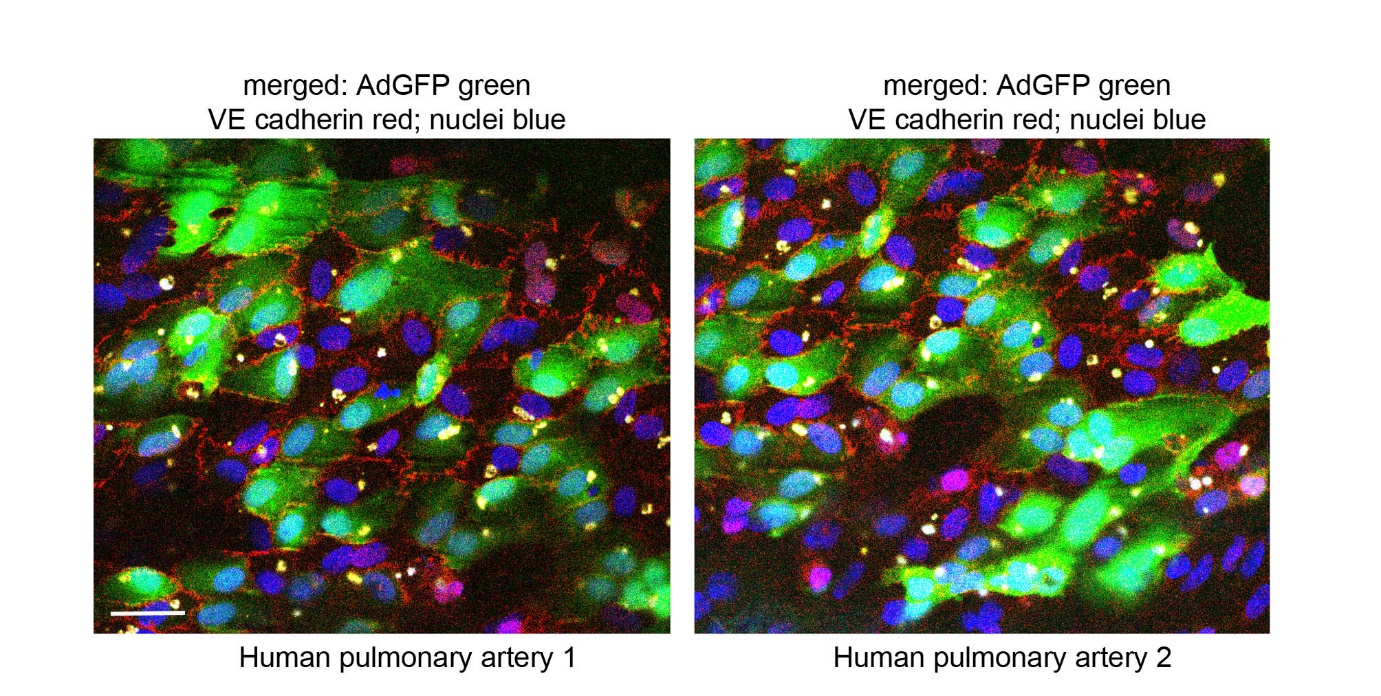

***Figure S4. Adenoviral transduction efficiency in the endothelium of human pulmonary artery explants****. Surgically removed pulmonary arteries were opened up and the endothelial side was incubated with AdGFP for 24 hours. In confocal images of the pulmonary arterial endothelium shown above, the cells overexpressing AdGFP are green, VE-cadherin is red and nuclei are blue, as indicated. Bar=20 µm.*

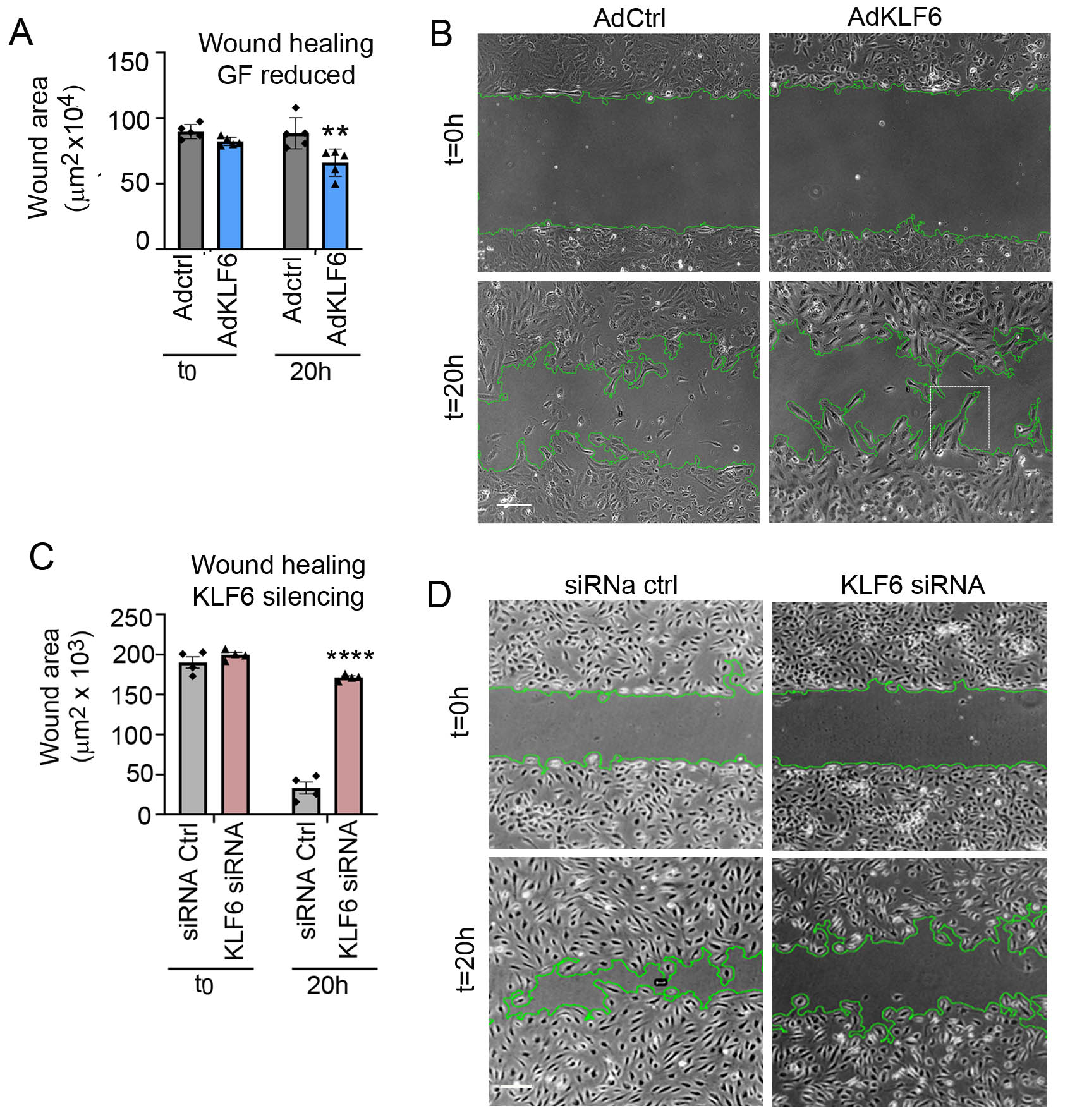

***Figure S5. Effects of KLF6 overexpression and silencing on endothelial wound healing in vitro.*** *Graph in* ***(A)*** *and corresponding light microscopy images in* ***(B)*** *show endothelial wound healing (measured as reduction in the wound area), in serum-and growth factor-starved control (AdCTRL) and KLF6-overexpressing (AdKLF6) HPAECs at t=0 and t=20h post-wounding. In* ***(B)*** *Boxed area shows well polarised AdKLF6-overexpressing cells migrating into the wound area. Graph in* ***(C)*** *and corresponding representative images in* ***(D)*** *show wound healing in HPAECs 48h post-transfection with control siRNA or KLF6 siRNA at t=0 or t=20h post-wounding. Green line demarcates wound edges. **P <0.01, ****P<0.0001, comparison with relevant treatment controls, one-way ANOVA with Tukey post-test; n=4-5. Bars show mean ±SEM.*

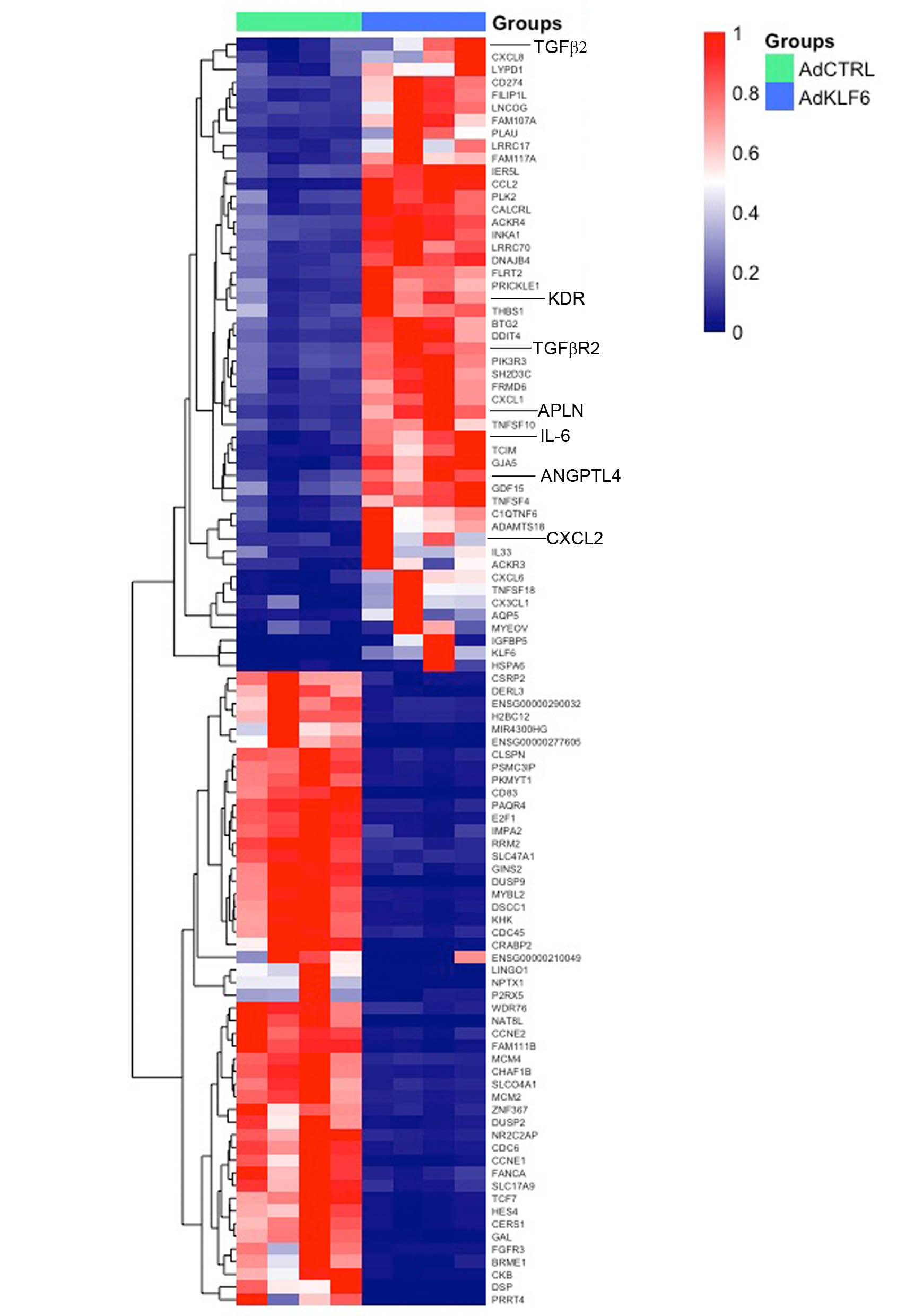

***Figure S6. Heatmap of top 50 upregulated and 50 downregulated KLF6-regulated DEGs.*** *Hierarchical clustering heatmap representing the top 50 up- and 50 down-regulated genes (FDR <0.05, log_2_*|FC| *<-0.25 or >0.25) upon KLF6 overexpression in HPAECs. The coloured scale bar represents z-score, where red represents upregulation, and blue represents downregulation.*

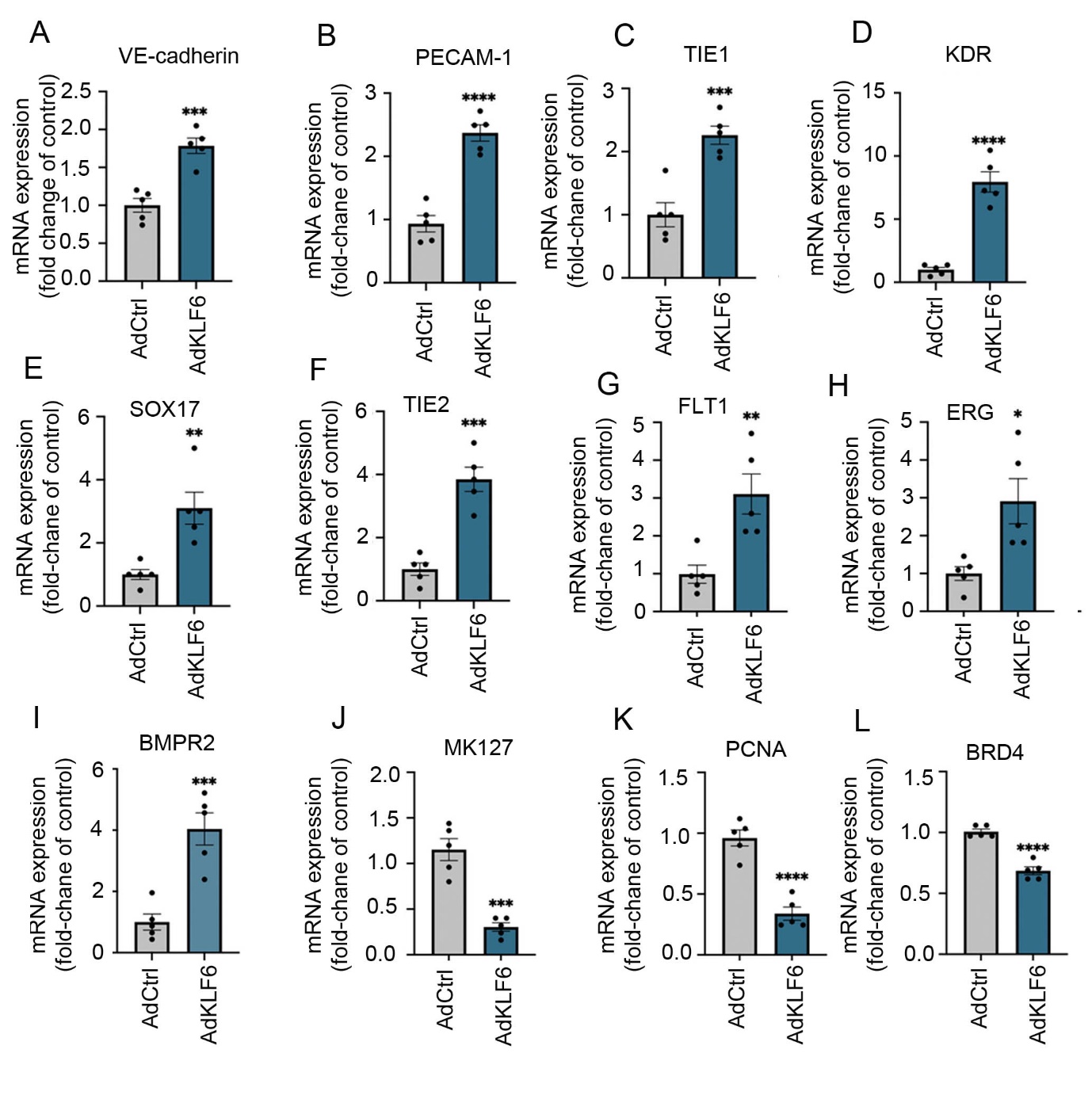

***Figure S7. Expression of selected KLF6-regulated genes.***

*Gene expression of VE-cadherin, PECAM-1, TIE1, KDR, SOX17, TIE2, FLT1, ERG, BMPR2, MK127, PCNA and BRD4 was measured by qPCR and data were normalised to β-actin. N=5, *P<0.05, **P<0.01, ***P<o,001, ****P<0.0001, comparison with AdCTRL; unpaired Student t-test, n=5.*

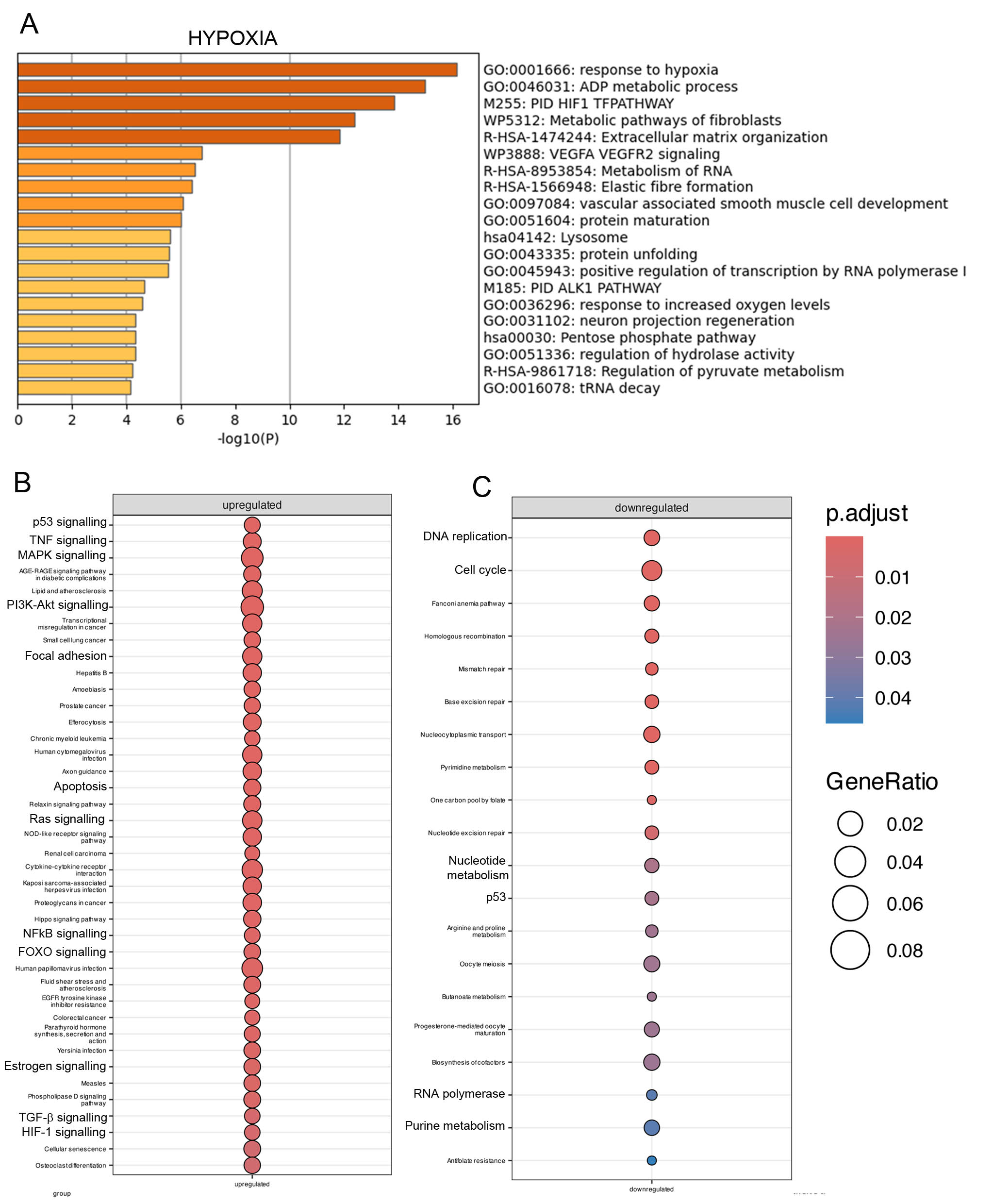

***Figure S8. Effect of KLF6 on hypoxia-induced changes in gene expression profile in HPAECs****.* ***(A)*** *Bar graph showing Metascape pathway and process enrichment analysis of differentially expressed genes following exposure of HPAECs to hypoxia (2% O2) for 24 hours. Bars are coloured by -log10 (p-value) and darker colour indicates more significant enrichment.* ***(B and C)*** *Dot plots showing the enriched KEGG pathways of DEGs (left, up-regulated genes; right, down -regulated genes) in KLF6 overexpressing HPAECs under hypoxic conditions. The colour of the dots represents the adjusted P-value, and the size of the dots represents the number of DEGs in the pathway*

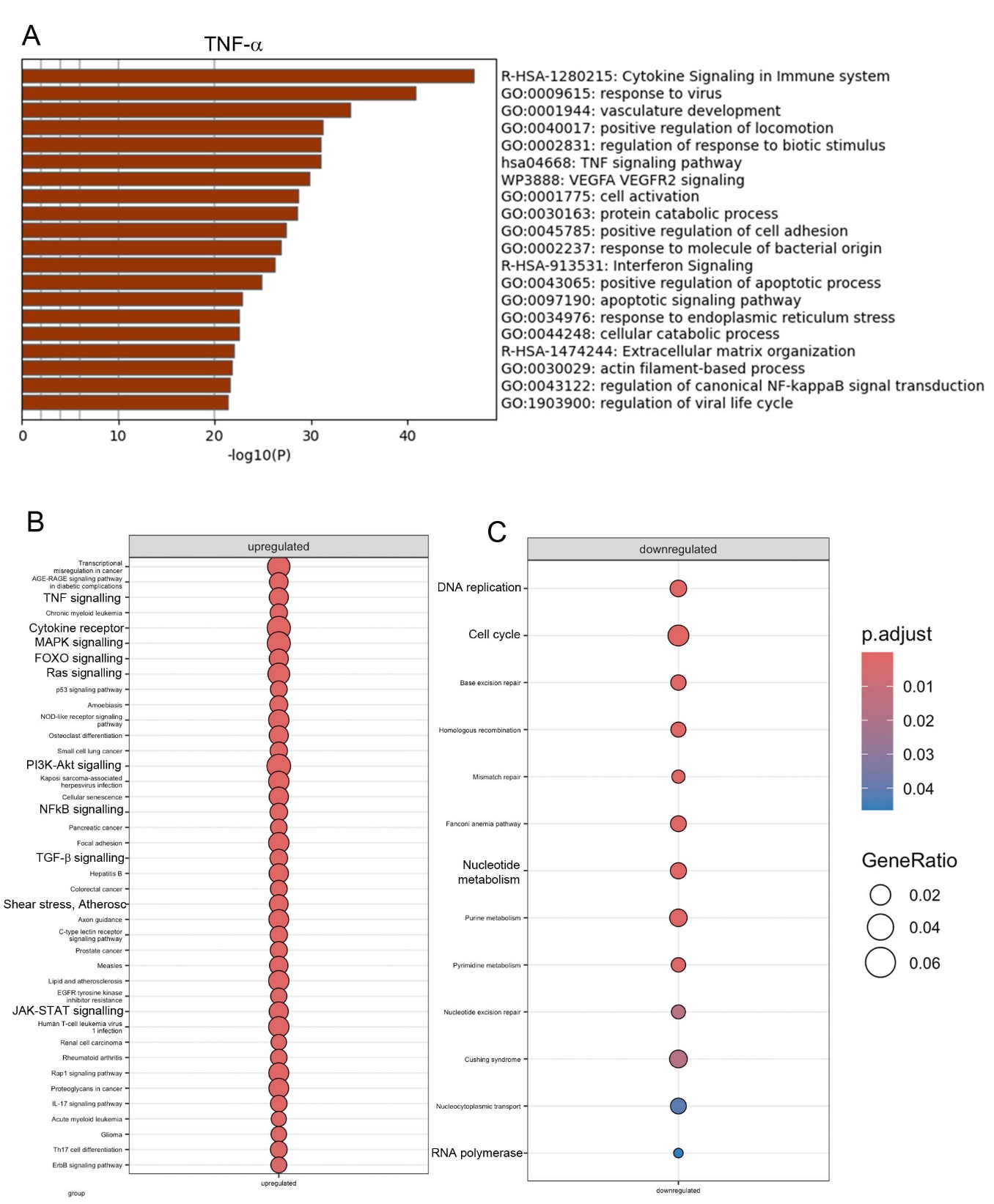

***Figure S9. Effect of KLF6 on TNF-a-induced changes in gene expression profile in HPAECs****.* ***(A)*** *Bar graph showing Metascape pathway and process enrichment analysis of differentially expressed genes following treatment of HPAECs with TNF-α (10 ng/ml) for 24 hours. Bars are coloured by -log10 (p-value) and darker colour indicates more significant enrichment****. (B, C)*** *Dot plots showing the enriched KEGG pathways of DEGs (left, up-regulated genes; right, down -regulated genes) in KLF6 overexpressing HPAECs treated with TNF-α. The colour of the dots represents the adjusted P-value, and the size of the dots represents the number of DEGs in the pathway*.

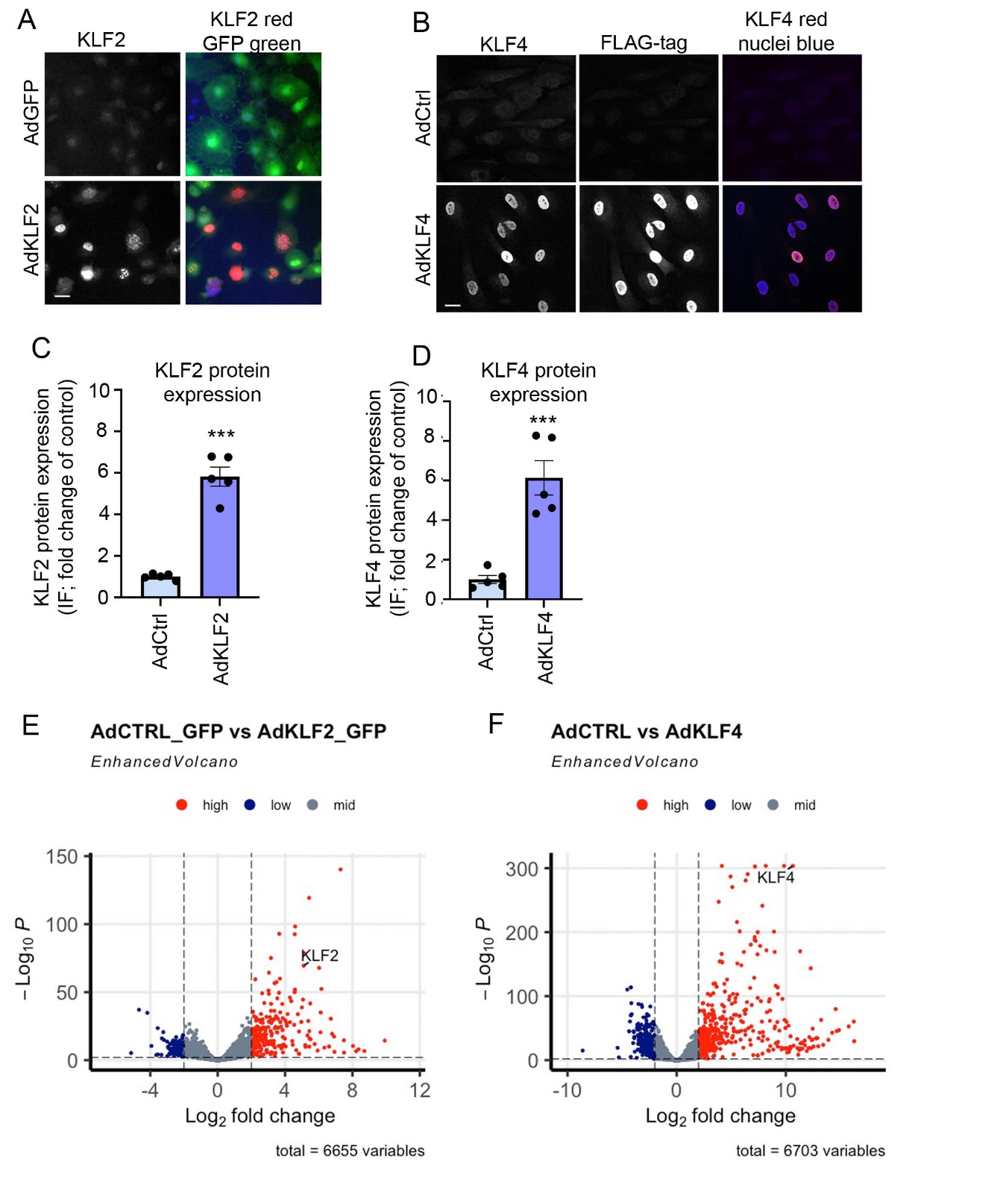

***Figure S10. Gene expression changes in HPAECs overexpressing KLF2 and KLF4. (A)*** *and* ***(B)*** *are representative confocal microscopy images showing localization of KLF2 and KLF4 in HPAECs 24h post- infection with AdGFP (adenoviral control; AdCtrl), AdKLF2-GFP or AdKLF4-FLAG, as indicated. Bar=10µm.* ***(C)*** *and* ***(D*** *show protein expression of KLF2 and KLF4 in HPAECs.* ***(E)*** *and* ***(F)*** *are volcano plots showing colour-coded changes in expression of genes induced by overexpression of KLF2 or KLF4, where downregulated genes and blue and upregulated genes are red. P<0.05, 0.25-fold cut off.*

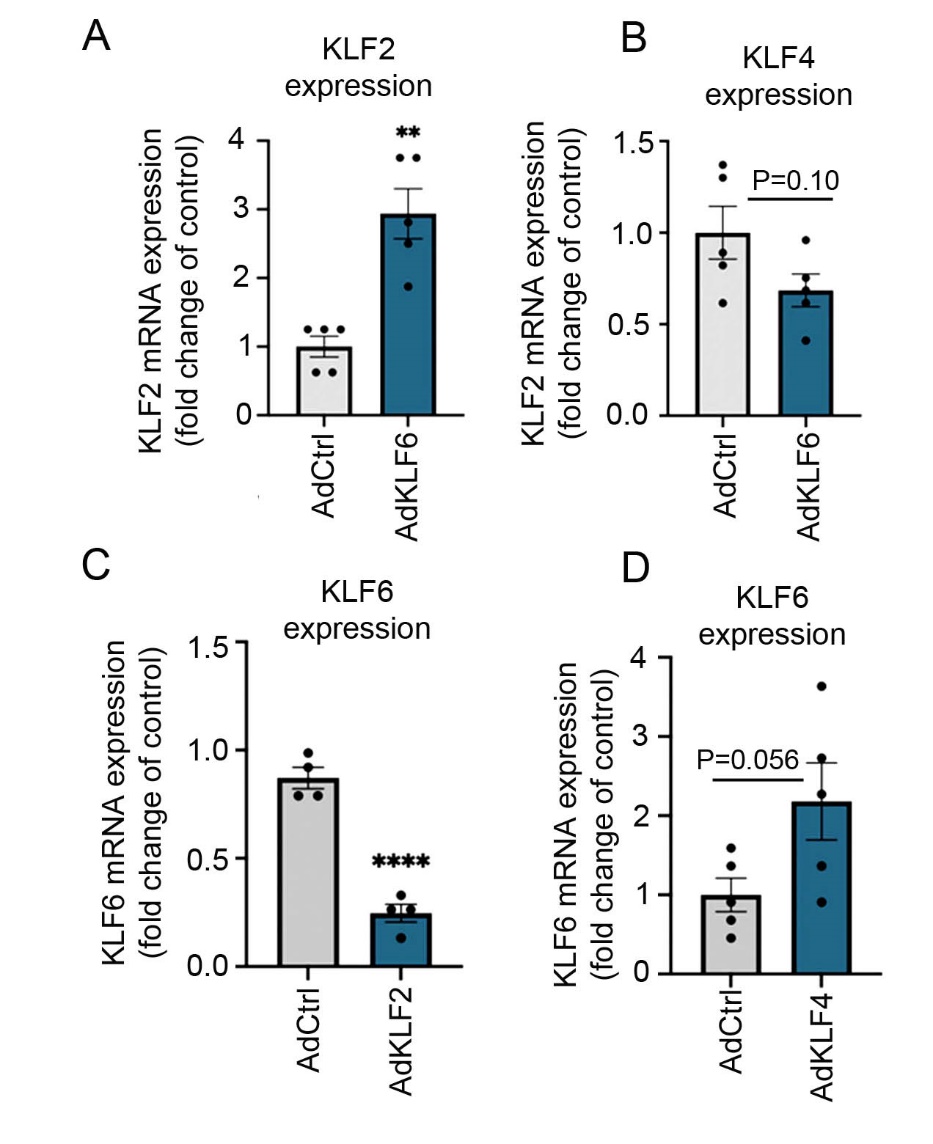

***Figure S11. Regulatory feedback relationships between KLF2, KLF4 and KLF6.***

***(A)*** *KLF2 mRNA and* ***(B)*** *KLF4 mRNA expression in HPAECs infected with AdCTRL or AdKLF6 for 24 h.* ***(C)*** *KLF6 mRNA expression in HPAECs infected with AdCTRL or AdKLF2 for 24 h.* ***(D)*** *KLF6 mRNA levels in cells infected with AdCTRL or AdKLF4 for 24 h, as indicated. In* ***(A-D)*** *data were normalized to the reference gene (β-Actin). **P<0.01; ****P<0.0001, comparison with adenoviral controls, unpaired t-test. Error bars indicate mean ±SEM; n=5.*

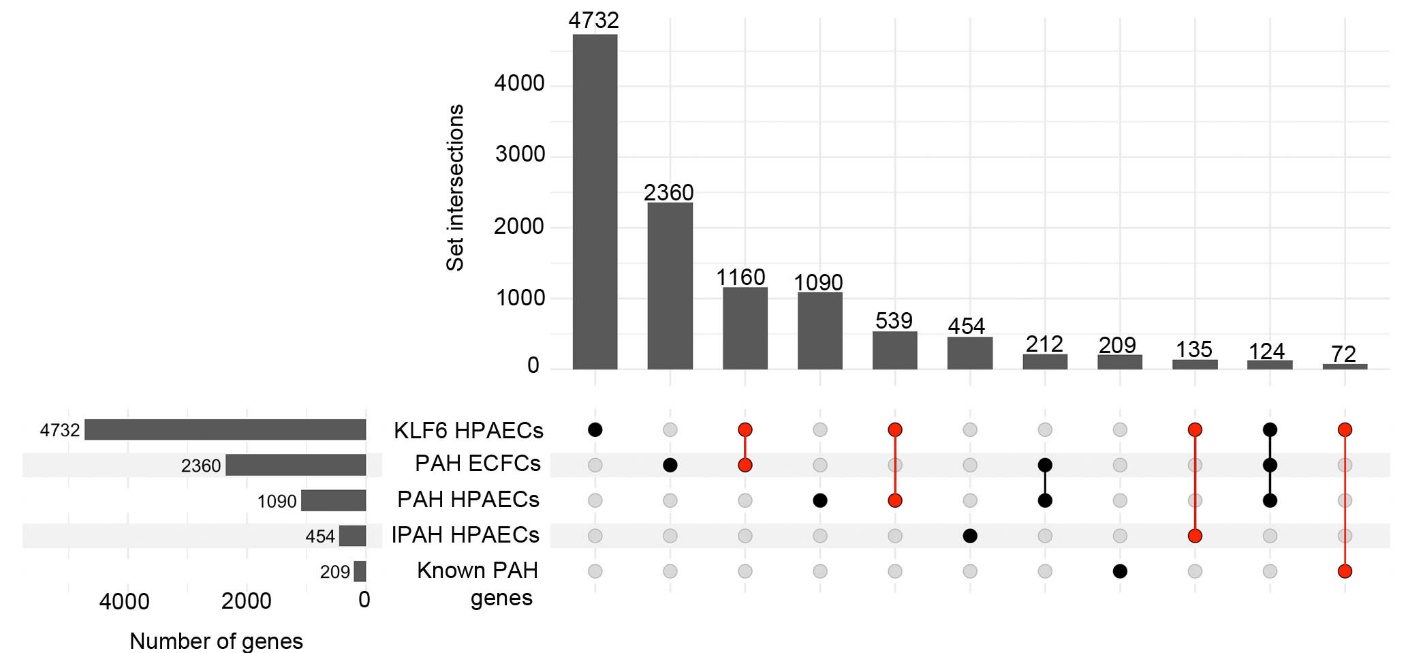

***Figure S12. Overlap analysis of differentially expressed genes in KLF6 overexpressing HPAECs and PAH DEG databases.*** *Upset plot displaying the interactions between differentially expressed gene lists of KLF6 HPAECs, PAH HPAECs, PAH ECFCs and IPAH PAECs, where connected black dots represent overlapping DEG lists and vertical bars in intersection set show number of overlapping genes; FDR<0.05. Gene lists are provided in Supplementary Table S3.*

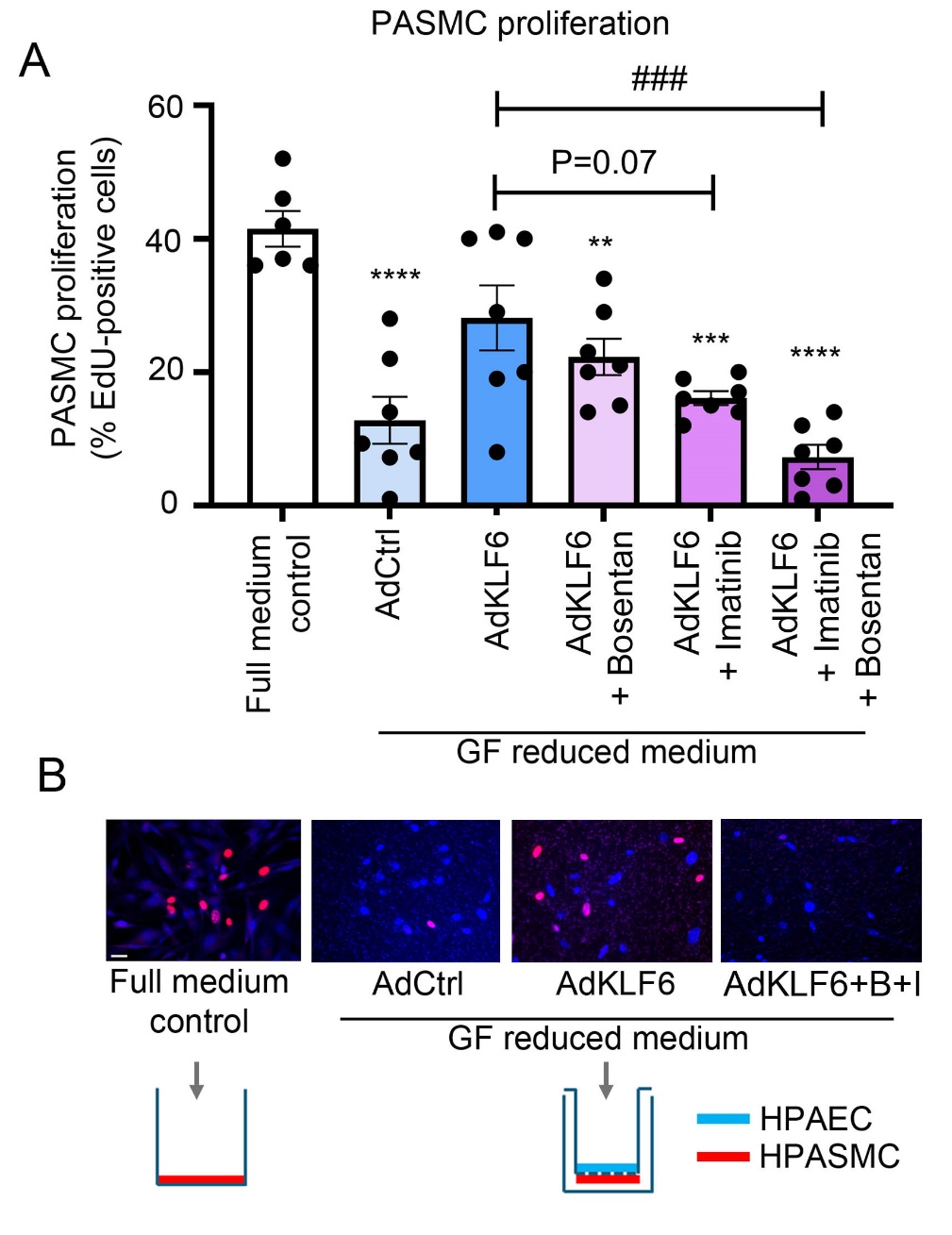

***Figure S13. Effect of endothelial KLF6 overexpression on HPASMC proliferation in vitro.***

***(A****) Graph showing proliferation of HPASMCs cultured alone or co-cultured with AdCTRL- or AdKLF6-treated HPAECs, as indicated; EdU incorporation assay. HPASMCs were cultured alone in optimal media (Full medium control) or were co-cultured with AdCTRL- or AdKLF6-overexpressing HPAECs in Transwell inserts, with the two cell types placed on either side of the porous membrane pore size 4µm in growth factor-depleted, serum-reduced (0.2% serum) medium for 16 hours. Bosentan (B; 20 µmol/L) and Imatinib (I; 10 µmol/L, were added to the cells separately, or in combination (B+I), as indicated. Error bars in (A) show means ±SEM; ^**^P<0.01,***P<0.0001, ^****^P<0.0001, comparisons with full medium control, ^###^P<0.001, comparison, as indicated; one way ANOVA, n=5 in control and n=7 in other experimental groups.* ***(B)*** *Corresponding representative fluorescent microscopy images of HPASMCs under different culture conditions, illustrated in diagrams below the images. Bar=10µm.*

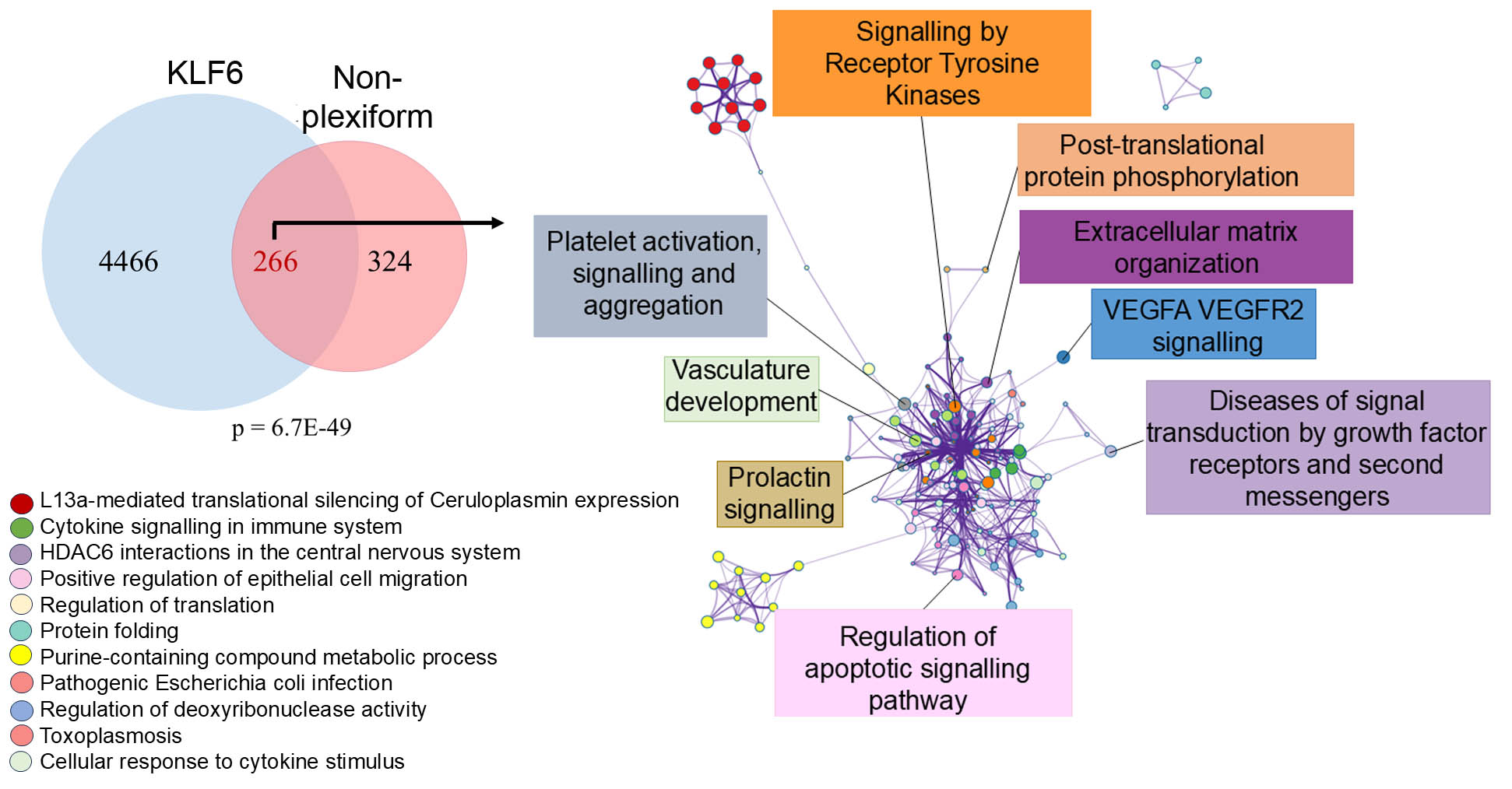

***Figure S14. KLF6-regulated DEGs in non-plexiform vascular lesions in PAH lung.*** *Venn diagram showing an overlap between KLF6-regulated DEGs in HPAECs and plexiform lesions DEGs identified by spatial transcriptomics; P= 1.1E-63, Fisher's exact test in the GeneOverlap package in R. The associated Metascape network of enriched terms in the shared pool of DEGs coloured by cluster ID, is shown on the right. FDR<0.05,*

*-0.25>log2|FC|>0.25*

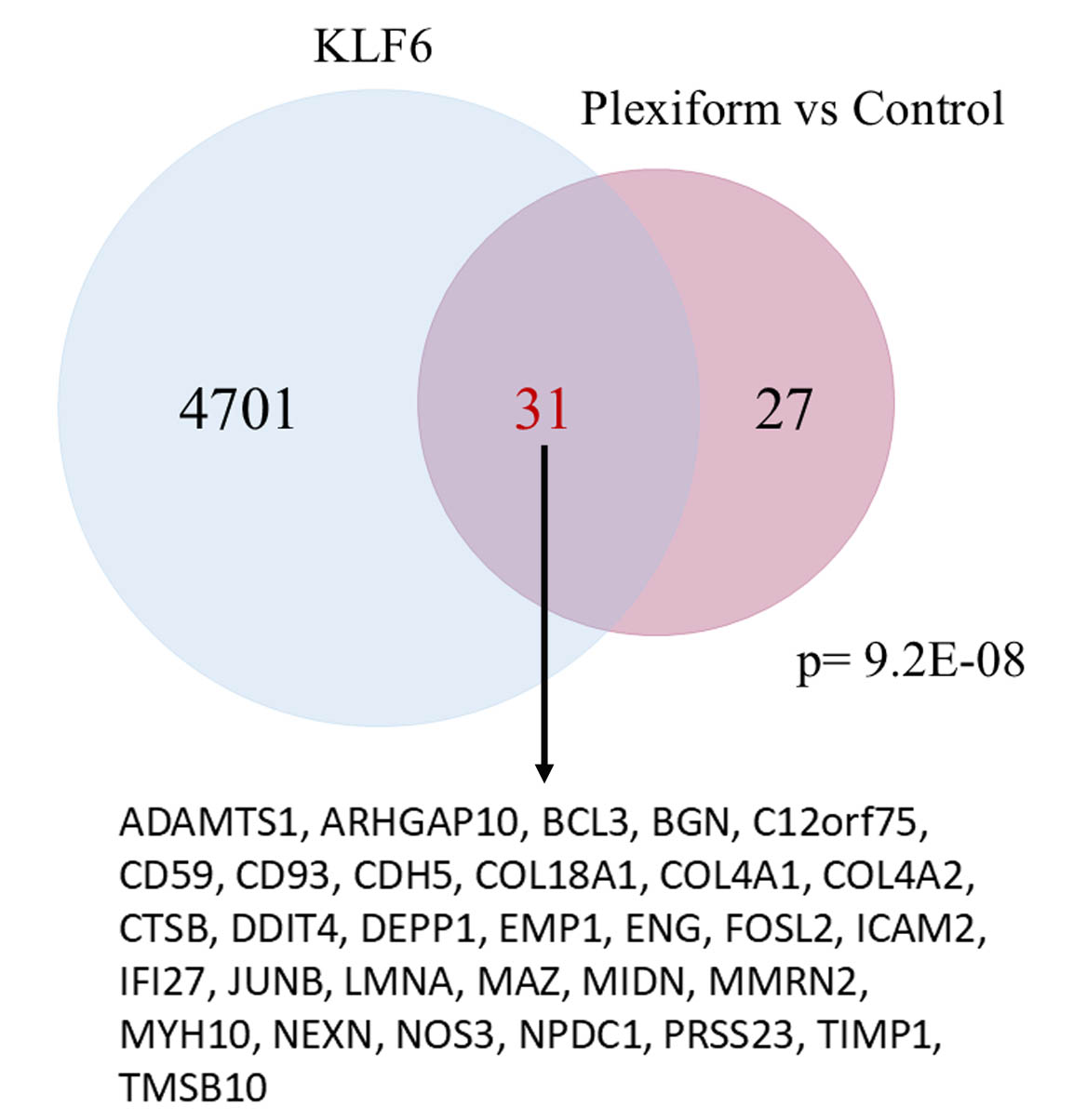

***Figure S15:*** ***Venn diagram showing an overlap between KLF6-regulated DEGs in HPAECs and plexiform lesion-specific DEGs in Tuder et al. (2024)****. The list of plexiform lesion-specific DEGs was taken from Table E6A (https://doi.org:10.1164/rccm.202307-1310OC)*

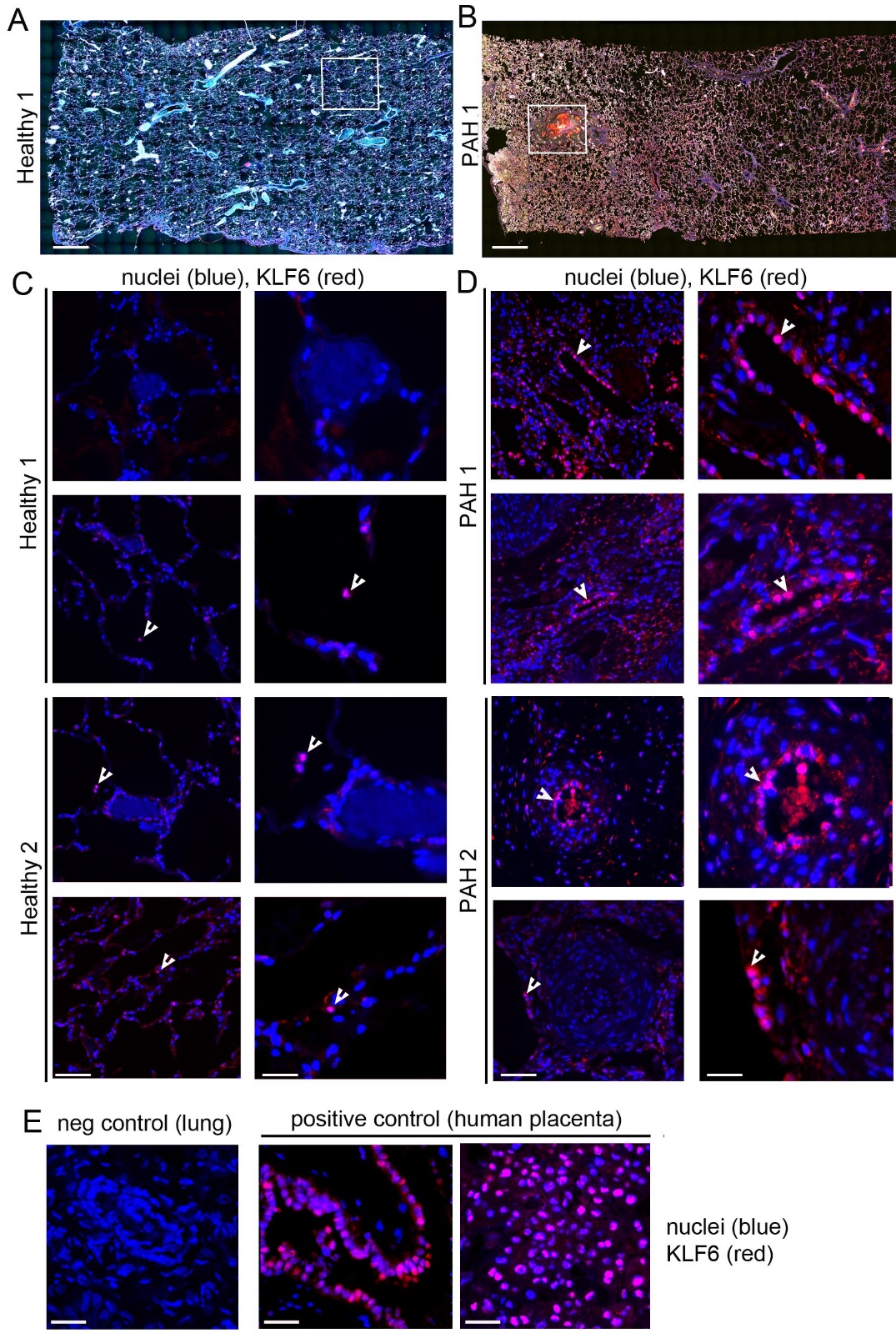

***Figure S16. Representative images of tissue sections used for spatial transcriptomic and KLF6 localization. (A)*** *Human healthy and* ***(B****) PAH lung, immunofluorescence: vWF (red), aSMA (yellow), nuclei (blue). Boxed areas indicate regions illustrated in (Healthy 1) and (PAH1) images below. Bar=500µm.* ***(C)*** *KLF6 localization in healthy and* ***(D)*** *PAH lung tissues from 2 donors; immunofluorescence: KLF6(red) and nuclei (blue); Bar=50µm. Enlarged corresponding images are shown in the right panel; Bar=20µm Arrowheads point to nuclear localization of KLF6 (pink).* ***(E)*** *negative control (secondary antibody only) and positive control showing localization of KLF6 in human placenta, as indicated. Bar=50µm.*

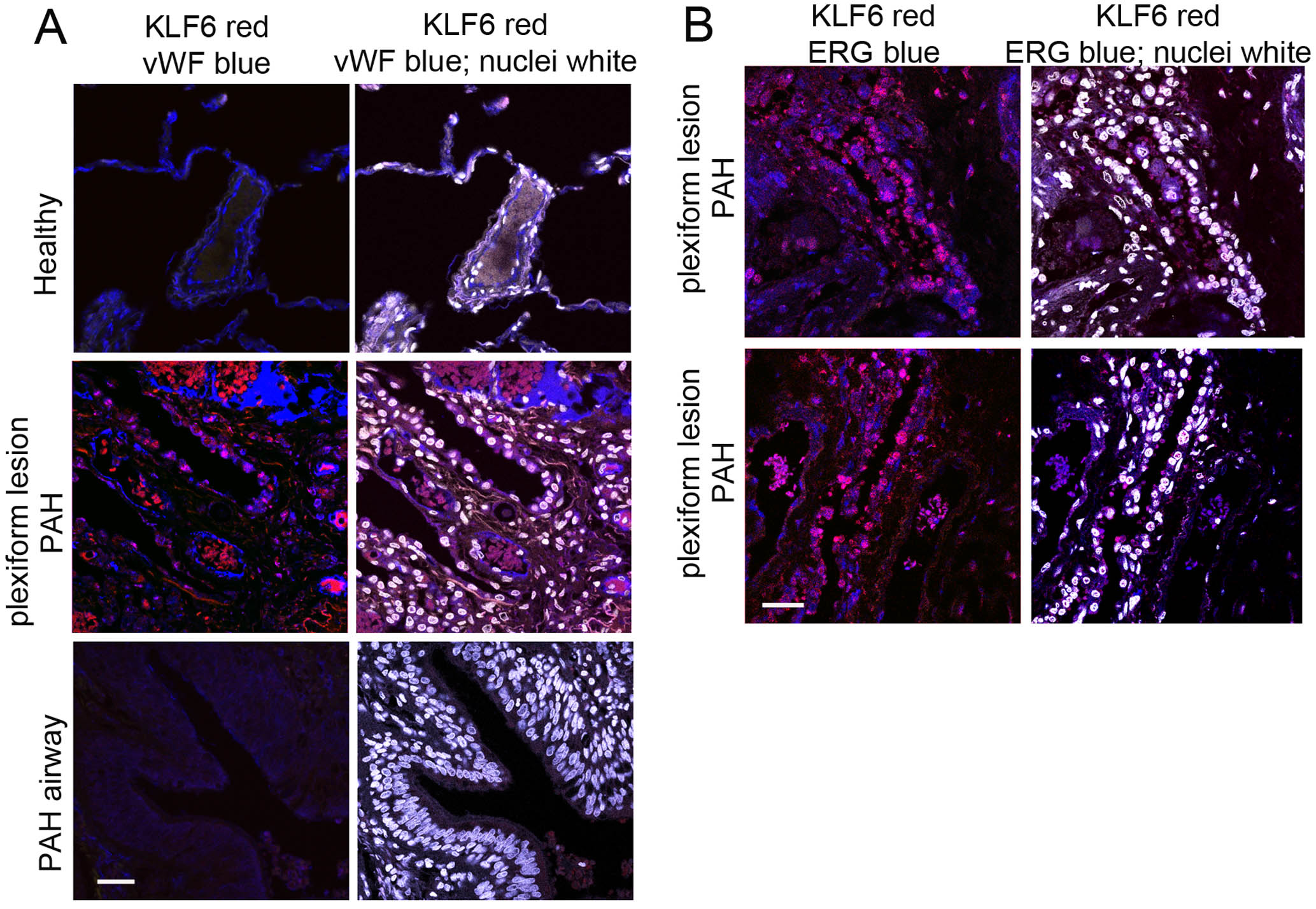

***Figure S17. KLF6 colocalization with endothelial markers Erg and vWF.*** ***(A****) vWF and* ***(B)*** *ERG immunostaining in healthy and PAH lung tissues, as indicated. KLF6 is red, vWF and Erg are blue and in merged images, nuclei are white. Immunofluorescence, pseudocolours. Bar = 50µm.*

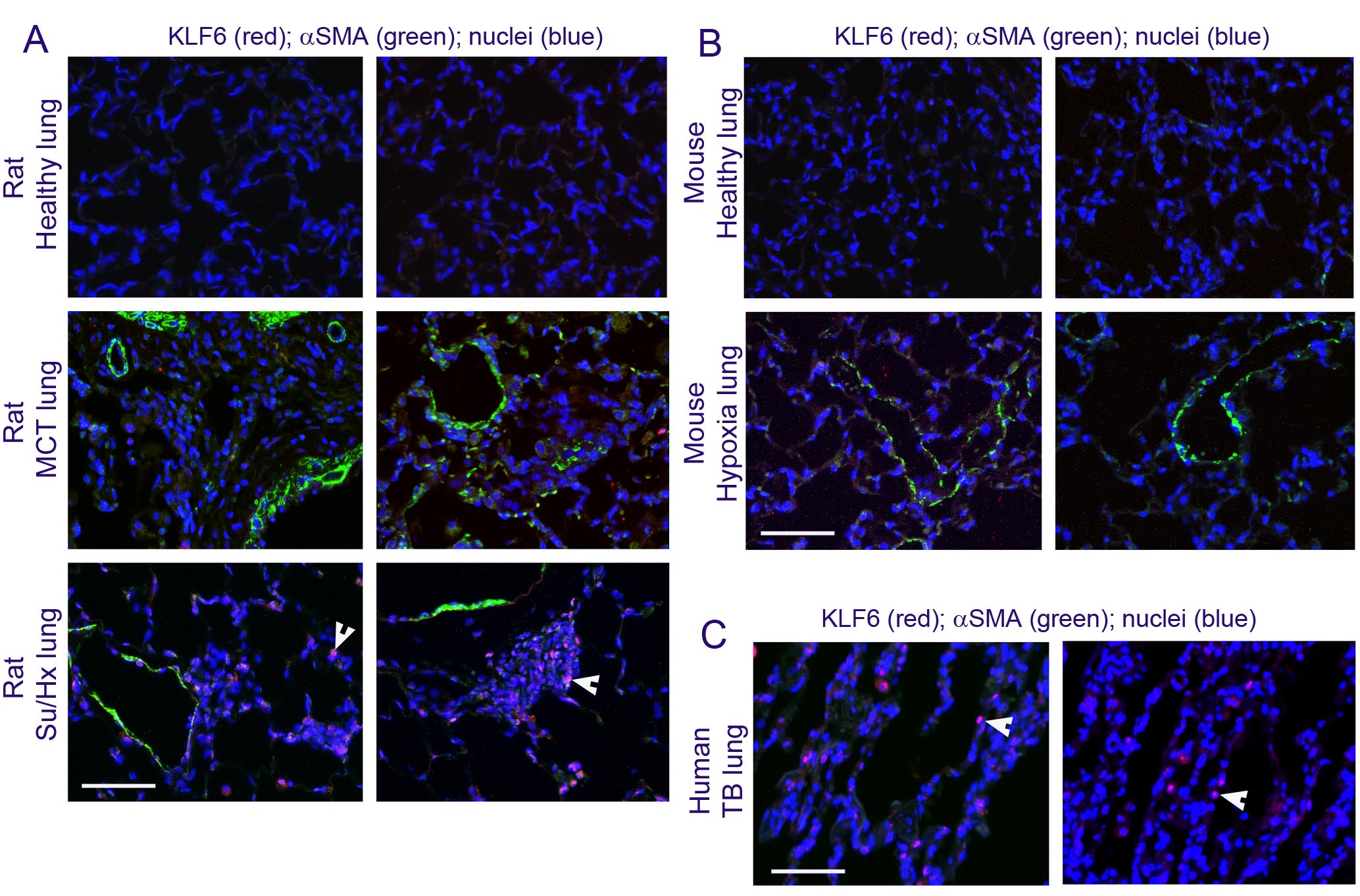
***Figure S18. KLF6 localization in rodent and human diseased lung.*** ***(A****) KLF6 localization in healthy, MCT and Sugen/hypoxia rat lung, as indicated;* ***(B)*** *KLF6 localization in healthy and chronically hypoxic mouse lung;* ***(C)*** *KLF6 localization in the lung of patient with tuberculosis. In (A-C) KLF6 is red, α-SMA is green and nuclei are blue; immunofluorescence. In (A) and (C) arrows point to nuclear localization of KLF6 (pink). Bar = 50µm.*

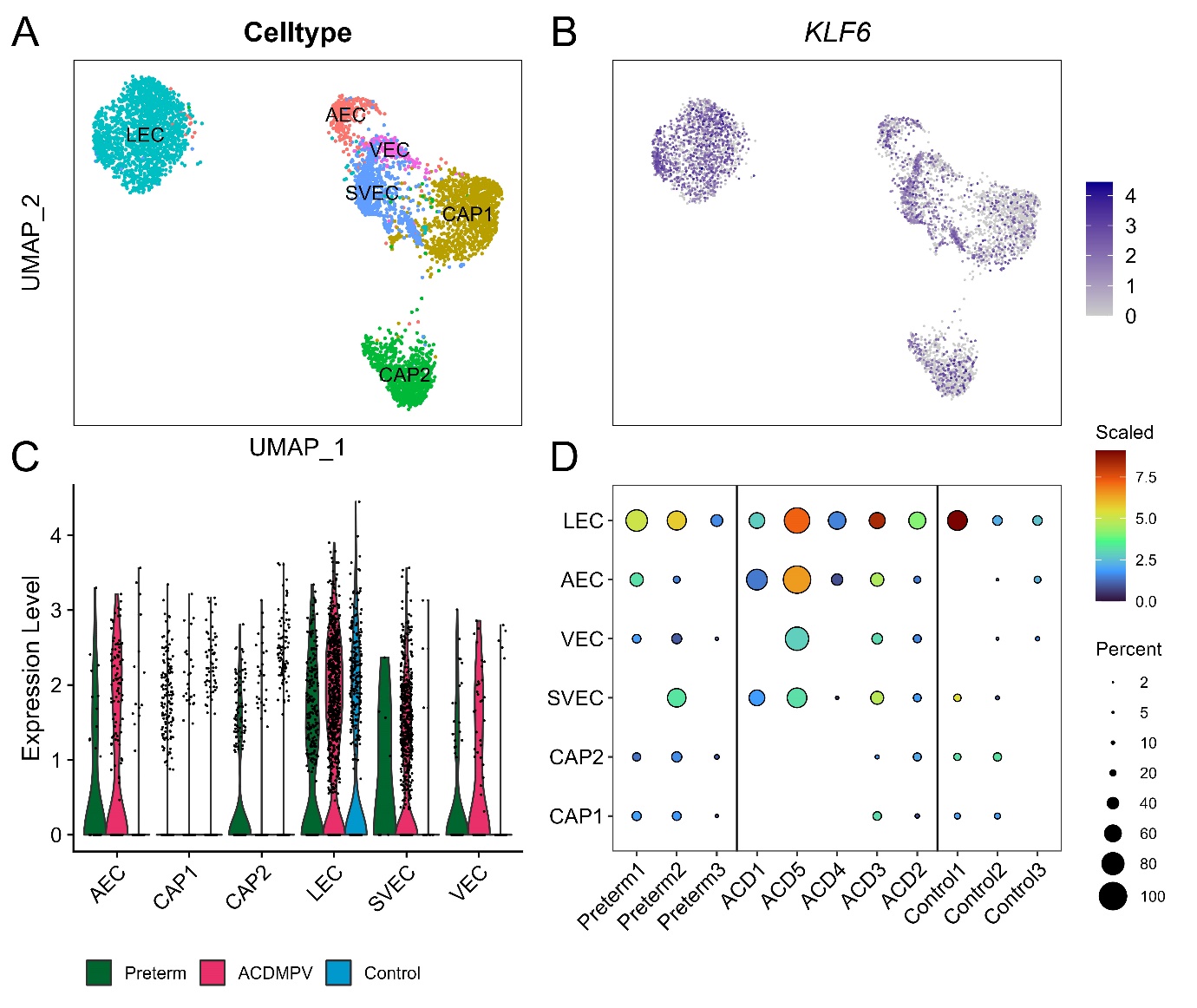

***Figure S19.*** ***Single-nucleus transcriptome analysis of KLF6 expression in endothelial cells (ECs) in alveolar capillary dysplasia with misalignment of pulmonary veins (ACDMPV) and control human lungs. (A****) UMAP of identified EC subpopulations* ***(B)*** *KLF6 expression in the identified EC populations. Cells with KLF6 expression were marked by blue colors and were plotted on top of cells without KLF6 expression (grey color).* ***(C)*** *and (****D****) show quantitative evaluation of KLF6 expression in different cell subpopulations, as indicated: AECs are arterial endothelial cells, VEC are venous endothelial cells, LECs are lymphatic endothelial cells, SVECs are systemic vascular endothelial cells and CAP1 and CAP2 are capillary cells. Differential expression was performed using Seurat 4 FindMarkers function with Wilcoxon rank sum test. Criteria were used for significance: p<0.05, FC>=1.5, pct>20%.*

SUPPLEMENTARY TABLES

**Table S1: List of unique DEGs for KLF2, KLF4 and KLF6.**

| Name | Unique genes |
| --- | --- |
| KLF2 | *ENSG00000196274, RN7SL718P, RN7SL670P, KLF2, ICAM1, ENSG00000237611, ENSG00000267834, RN7SL455P, RN7SL602P, ENSG00000210100, ENSG00000278903, MIR3648-1, EZH2, CABLES2, MIR663AHG, HSPA14, GINS3, TRAF3, MSH6, ACD, FAM210A, ENSG00000234160, SCML1, ENSG00000280800, SNHG1, PRC1, USP36, LOC122526782, SRSF7, TOP3A, NCAPD3, TEX30, RN7SL3, IMMP1L, PHGDH, ENSG00000255198, HLA-DPB1, CCDC14, TFAM, RFLNB, RRP12, RANBP1, FAM124B, SUOX, TRIM69, MX2, HHIP, TFPI, IFI6, OAS2, TNIK, NRP1, XAF1, TRIM5, ISG15, GRAP, GIMAP8, RIGI, MX1, ERG, AMIGO2, STAT2, EPSTI1, MARCKS, LYL1, ZNF561, RNF144B, IFI35, RSAD2, TM4SF1, CASP1, ENC1, DUSP6, EFNA1, JUP, SAMD9L, CD93, EDN1, IFI44L, ENSG00000289833, TP53I11, FAM43A, STAT1, IFITM1, ALDH1A1, LOC102724951, SCHIP1, CLIC1, CXCL11, CXCL10* |
| KLF4 | *SEMA3F, LASP1, ATOSB, ST3GAL1, IL32, PLAUR, DNASE1L1, SERPINB1, OSBPL5, TIMP2, VCAN, RRM2B, NEDD4L, ATP2B4, SLC2A3, TNK2, GPC1, DLX3, DGKA, SLC9A3, ELOVL1, RASGRP2, PPP2R5B, BCL3, FSTL3, GBA2, STK10, ATP2A3, MGLL, FSCN1, NMRK2, LRCH4, TP53INP2, TNS1, CHRNA3, BCORL1, WNT11, PTHLH, PPP1R13B, SMOX, SLC8B1, RCOR1, CHRD, UNC13D, TBC1D2, CBARP, PALM, MKNK2, SH3BP1, SUN2, APOL1, GSKIP, CPNE6, ELF4, TIMP1, MEDAG, MT4, MMP15, NDRG4, CEMIP, NDRG1, KCNN4, TUBB4A, CKM, DMPK, SNAPC2, EBI3, CNFN, CYTH2, CLEC11A, RASA4, LFNG, TESK1, CREB3, VSIR, SLC9A3R1, CDR2L, PMP22, ALDOC, KLF3, UGDH, SLC15A3, CD81, PITPNM1, KRT18, ART4, MGP, RAB5B, VEGFA, HES1, TNNC1, CLCN2, IL1RL1, GBP3, VAMP8, PGF, FKBP1B, TEK, FABP3, GLIPR2, DNPEP, SNAI1, PMEPA1, GRM4, LRRFIP1, WNT1, MT1G, DOK4, IRF1, TNFSF9, C3, ID1, SLURP1, ARMCX1, EDN2, FBXL12, APOL2, RAC2, FLNC, LOXL1, KLK14, ADCY4, CDKN1C, TNNI3, HELZ2, H19, CALY, ULBP2, EPS8L1, ACSS2, NFATC1, MAP1B, RARA, DNAJB1, TRIM21, CARD6, HSPA12B, CASP9, CCNA1, PLAAT4, IRAK2, BHLHE40, BIN1, KLF4, TUBB2A, IER3, IL18BP, LRRC32, SQOR, DUSP5, SHF, CERS5, RHOF, BAHD1, PSTPIP1, OSGIN1, CBX4, STAC2, COL6A1, COL6A2, ATP1B1, EFNA3, CSRNP1, SLC16A2, SLC25A25, LCN2, ADIRF, TNKS1BP1, TAGLN, HSPB8, ZFP36L2, CHD1, TRIM11, RABGEF1, ADAMTS1, KLF10, PCDH1, MYO1E, TMEM164, MRAS, CDA, SPON2, ZBTB7B, LMNA, SQSTM1, SCGB3A1, PLCD3, TAMALIN, BCL6B, HS3ST6, FGF19, LAPTM5, SYNC, KIAA1522, CAMK2N1, ATF3, IER5, IHH, MELTF, DUSP7, CSF2, MICALL2, TSC22D4, GOLGA2, CA4, MIDN, SERTAD3, PSCA, LY6D, RILP, KLK1, KLK13, SPRYD3, NSG1, LGALS9, ASMTL, NPR1, FASN, KLF13, CST1, KRT8, KISS1, PFKFB3, SERPINB9, HTRA3, SOX7, BBLN, JUNB, KRT19, KRT13, CXXC5, ZNF274, SPSB1, MAL, ORMDL3, LRRC15, ISG20, CEBPB, PRSS27, RAB43, OVOL1, RARG, HEG1, RNF213, PHLDA3, SLCO2A1, DES, MARCKSL1, DDIT3, TBC1D10C, DRAP1, BAIAP2, TPRN, LPCAT4, EPS8L2, ULK1, CASKIN2, GPR4, PNPLA2, CRACR2B, MLF1, ZFAND2A, PARP10, THBD, ZBTB7A, CDC42EP4, PJA1, PHLDA2, ADGRB1, FBXL6, TSKU, AFMID, SMTN, OAF, ALDH1A3, PTP4A3, SOCS3, TRARG1, DGAT1, MAFF, SIGIRR, IL3RA, IRF7, AHNAK2, ARAP1, PDE2A, PRR5, VSIG10L, KRT16, HEXIM1, KRT14, TRABD2A, ESPN, PLCD1, MT1X, BCAM, PEAR1, S100A3, FAM25A, FAM83G, SBSN, FAM25G, CLDN4, S100A4, HRCT1, S100A2, LAMB3, PDLIM7, SERTAD1, LYPD2, STMN3, EOLA1, SLC22A20P, AKAP17A, TPM2, HSPA1B, PSORS1C2, CDSN, C6orf15, AKT1S1, ENSG00000204791, ADGRG1, MT1M, KRT6A, RNU6-36P, RNU6-33P, GPX3, ACKR1, LTC4S, LCAT, DDAH2, PPM1N, NPTXR, PLEKHM1, CBLL1-AS1, PLEKHO2, PEG10, TNFRSF6B, GPR162, LINC00520, WFDC21P, SCRT1, LIN37, SLURP2, ENSG00000289316, ENSG00000291309, SCMH1, POLA2, BID, MRTO4, MBD3, TIPIN, SPAG5, UBE2T, CAD, BIRC5, NAT14, VRK1, NME4, BCAT2, SNX8, NCS1, NPM3, CCDC34, PTPMT1, FOXM1, OGG1, GPX7, POLR1G, DYNC2I2, CLCC1, CSE1L, PDCD2L, BCL2L12, SRD5A3, PAICS, DUT, GAMT, EXOSC2, SCLY, PCNA, NASP, ADCK2, LPIN1, DZIP1, CDK4, RNASEH2B, POLR1E, SLC37A4, ATIC, HERC5, GCSH, ARRB2, NFIC, PARP1, GINS4, SIGMAR1, GBGT1, B3GAT3, GXYLT1, SPC25, ADGRA3, AGPAT5, NDUFAF6, CIP2A, CXCL5, RPL39L, CDCA7L, FABP5, COMTD1, PACSIN3, CLMP, C19orf48, PBK, DTYMK, CDK1, FABP4, SPTBN2, SPHK1, PFAS, AURKB, RCC1, IDH2, SFXN4, WDR5, IPO4, TFDP1, JPT2, STIMATE, BOP1* |
| KLF6 | *DCBLD2, KLF6, PIR, PALMD, CAV1, SMURF2, SULT1E1, NEDD9, PLSCR4, ST3GAL5, RND3, GADD45A, PIK3R3, TNFSF18, FAM117A, CDKN1A, CXCL6, LRRC17, AJUBA, RNASE1, ITGB4, SLC38A2, MDM2, RAPGEF5, IL33, BCAR3, HECW2, ARHGAP24, PRICKLE1, ACVRL1, LPAR6, CCN1, MALL, HMGA2, ROBO4, LY96, TSC22D3, ETS2, BTG2, VCAM1, SOX17, MAMDC2, MTMR10, CXCL8, ZMAT3, CD34, C11orf68, PRR15, CITED4, FAM167B, ZFP36L1, KANK3, PDCD1LG2, LGR4, ENSG00000233818, LINC00607, RPL32P29, MIR100HG, ENSG00000256664, LNCOG, TXNIP, NCAPH2, MTHFD1, IL27RA, FBRSL1, H2BC11, XRCC3, TUBA4A, TOMM40, HAUS2, DNAJA4, SLC39A4, PDSS1, MKI67, SLC43A1, PUSL1, MCRIP2, SHMT1, POLE, SLC25A22, SEPTIN5, TOR3A, H2AX, ATAD3A, EIF4EBP3, EEF1AKMT4* |

**Table S2: List of differentially expressed genes shared between KLF2, KLF4 and KLF6**.

All have gene symbols or Ensemble ID.

| Comparison List | Overlapping genes |
| --- | --- |
| KLF2, KLF4 and KLF6 | *ANGPTL4, APOE, ASF1B, ATAD2, ATAD3B, BRME1, CCNE1, CDC25A, CDC45, CDC6, CDCA5, CDK2, CENPK, CENPM, CENPQ, CENPU, CENPV, CHAF1B, CKB, CLDN5, CLSPN, CRABP2, CX3CL1, DERL3, DNMT1, DSCC1, DSP, DTL, DUSP2, DUSP9, EEF1A2, EMP2, ENSG00000259316, ENSG00000277605, ENSG00000279573, ENSG00000290032, ENSG00000291224, ERFE, ESCO2, EXOSC5, FAM111B, FANCI, FEN1, FGFR3, FOXRED2, FRMD6, FSD1, GINS2, H2BC12, HADH, HELLS, IL6, KHK, LINC01551, LINGO1, LRP3, MAD2L1, MCM2, MCM3, MCM4, MCM5, MCM6, MCM7, MCM8, MIR4300HG, MSH2, MT1A, MYBL2, MYO19, MYRF, NAT8L, NETO2, NPTX1, NR2C2AP, OLFM2, P2RX5, PCLAF, PDGFB, POLA1, PRRT4, PSMC3IP, RAD51, RAD51AP1, RBBP8, RECQL4, RFC2, RFC3, RFC4, RFC5, RRM1, RRM2, SDC1, SLC25A19, SLC47A1, TEDC2, TMEM106C, TONSL, TYMS, UHRF1, UNG, VASN, WDR76, ZNF367, ZWINT* |
| KLF2 and KLF4 | *ANGPTL4, APOE, AQP1, AQP3, ARC, ASF1B, ASS1, ATAD2, ATAD3B, BCYRN1, BRME1, C11orf96, CA2, CCNE1, CD52, CD55, CDC25A, CDC45, CDC6, CDCA5, CDK2, CENPK, CENPM, CENPQ, CENPU, CENPV, CHAF1B, CKB, CLDN5, CLIC3, CLSPN, COL1A1, COL2A1, COL9A3, COMP, CRABP2, CRIP1, CRLF1, CSPG4, CX3CL1, DEPP1, DERL3, DNMT1, DSCC1, DSP, DTL, DUSP2, DUSP9, ECM1, EEF1A2, EMP2, ENSG00000259316, ENSG00000277605, ENSG00000279573, ENSG00000282993, ENSG00000290032, ENSG00000291224, ERFE, ESCO2, EXOSC5, FAM111B, FANCI, FEN1, FGFR3, FOXRED2, FRMD6, FSD1, GFPT2, GIMAP6, GINS2, H2BC12, HADH, HBA1, HBA2, HELLS, IFIT1, IGF2, IGFBP6, IL11, IL6, KHK, KISS1R, KLK10, LCN6, LINC01551, LINGO1, LRP3, MAD2L1, MCM2, MCM3, MCM4, MCM5, MCM6, MCM7, MCM8, MIR4300HG, MSH2, MT1A, MYBL2, MYO19, MYRF, NAT8L, NETO2, NGEF, NGFR, NOS3, NPPC, NPTX1, NR2C2AP, OLFM2, P2RX5, PCLAF, PDGFB, PI16, PLVAP, POLA1, PRRT4, PSMC3IP, PTGDS, RAD51, RAD51AP1, RBBP8, RECQL4, RFC2, RFC3, RFC4, RFC5, RRAD, RRM1, RRM2, S100P, SCNN1D, SDC1, SLC17A7, SLC25A19, SLC47A1, SLC6A8, SOX8, TEDC2, TMEM106C, TNFAIP2, TNNT1, TNXB, TONSL, TRH, TYMS, UHRF1, ULBP1, UNG, VASN, WDR76, ZNF367, ZWINT* |
| KLF2 and KLF6 | *ACKR4, ADAMTS18, ADCY3, AK4, ANGPTL4, ANKRD50, APLN, APOC1, APOE, ARHGAP18, ARRDC3, ASF1B, ATAD2, ATAD3B, BBC3, BMP4, BMX, BRME1, BTG3, CALCRL, CCDC80, CCL2, CCNE1, CCNE2, CD274, CD83, CDC25A, CDC45, CDC6, CDCA5, CDK2, CDT1, CENPK, CENPM, CENPQ, CENPU, CENPV, CERS1, CHAF1A, CHAF1B, CHRNA5, CKB, CLDN11, CLDN5, CLEC14A, CLSPN, CRABP2, CSRP2, CX3CL1, DDIT4, DERL3, DNAJB4, DNMT1, DONSON, DSCC1, DSP, DTL, DUSP2, DUSP9, E2F1, EEF1A2, EMCN, EMP2, ENO2, ENSG00000259316, ENSG00000267102, ENSG00000267583, ENSG00000277605, ENSG00000279337, ENSG00000279573, ENSG00000290032, ENSG00000291224, ERFE, ESCO2, EXOSC5, FAM111B, FANCA, FANCI, FEN1, FGFR3, FGFRL1, FILIP1L, FLRT2, FOXRED2, FRMD6, FSD1, GAL, GASK1B, GBP2, GIMAP2, GIMAP7, GINS2, GJA5, GMNN, H2BC12, HADH, HELLS, HES4, IL6, INKA1, KDR, KHK, LDB2, LINC01013, LINC01235, LINC01551, LINGO1, LIPG, LRP3, LRRC70, LYPD1, MAD2L1, MCM2, MCM3, MCM4, MCM5, MCM6, MCM7, MCM8, MIR4300HG, MSH2, MT1A, MTHFD2, MYBL2, MYO19, MYRF, NAT8L, NETO2, NPTX1, NR2C2AP, NR2F2, NREP, OLFM2, ORC6, P2RX5, PAQR4, PCLAF, PDGFB, PDXP, PHLDA1, PKMYT1, PLK2, POLA1, POLD3, PPIF, PRPS2, PRRG1, PRRT4, PSMC3IP, PTX3, RAD51, RAD51AP1, RBBP8, RECQL4, RFC2, RFC3, RFC4, RFC5, RGS3, RRM1, RRM2, SAMD1, SDC1, SH2D3C, SLC25A19, SLC39A8, SLC43A3, SLC47A1, SLC7A5, SLCO4A1, SMCO4, SOX4, SPAAR, SPRY2, TCF19, TCF7, TCIM, TEDC2, TGFB2, TGFBR2, THBS1, TK1, TM4SF18, TMEM106C, TMEM140, TMEM201, TMEM38B, TNFSF10, TNFSF4, TONSL, TSEN54, TUBGCP4, TYMS, TYRO3, UHRF1, UNG, VASN, WDR76, ZMYND8, ZNF189, ZNF367, ZNF521, ZSCAN31, ZWINT* |
| KLF4 and KLF6 | *ACKR3, ADAMTS4, ADAMTS9, ALKBH2, ANGPTL4, APOE, AQP5, ASF1B, ASNS, ATAD2, ATAD3B, BCL2L1, BRME1, C1QTNF6, CCNE1, CD82, CDC25A, CDC45, CDC6, CDCA5, CDK2, CENPH, CENPK, CENPM, CENPQ, CENPU, CENPV, CEP78, CHAF1B, CITED2, CKB, CLDN14, CLDN5, CLN6, CLSPN, CMSS1, CRABP2, CX3CL1, CXCL1, CXCL2, DERL3, DHFR, DNMT1, DPH2, DSCC1, DSP, DTL, DUSP2, DUSP9, EEF1A2, EFNB1, EMP2, ENSG00000188985, ENSG00000210049, ENSG00000256514, ENSG00000259316, ENSG00000273420, ENSG00000277605, ENSG00000279573, ENSG00000290032, ENSG00000291224, ERFE, ESCO2, EXOSC5, FAM107A, FAM111B, FAM216A, FANCG, FANCI, FEN1, FGFR3, FOS, FOSL2, FOXRED2, FRMD6, FSD1, GDF15, GINS2, H2BC12, H4C14, HADH, HAGHL, HELLS, HSPA6, HSPD1, IER5L, IFI30, IGFBP5, IL6, IMPA2, KHK, KRT17, LINC01551, LINGO1, LRP3, MAD2L1, MCM2, MCM3, MCM4, MCM5, MCM6, MCM7, MCM8, MELK, MIR22HG, MIR4300HG, MRE11, MSH2, MT1A, MYBL2, MYEOV, MYO19, MYRF, NAT8L, NCAPG2, NCKIPSD, NETO2, NPTX1, NR2C2AP, OLFM2, P2RX5, PCDH12, PCLAF, PDGFB, PIMREG, PLAU, POLA1, POLD1, PRIM1, PRRT4, PSAT1, PSMC3IP, PTPRE, RAC3, RAD51, RAD51AP1, RBBP8, RECQL4, RFC2, RFC3, RFC4, RFC5, RGCC, RGS4, RNASEH2A, RRM1, RRM2, SDC1, SKP2, SLC17A9, SLC19A1, SLC25A10, SLC25A19, SLC47A1, SLC5A6, SPNS2, SRM, TCOF1, TEDC2, TMEM106C, TMEM217, TMEM97, TONSL, TRAF3IP2, TRIP13, TSC22D1, TYMS, UHRF1, UNG, VASN, VPS9D1-AS1, WDR76, YDJC, ZNF367, ZWINT* |

**Table S3: List of differentially expressed KLF6 DEGs in PAH databases.** KLF6 DEGs were identified in the RNAseq databases of PAH HPAECs, PAH ECFCs (Ainscough et al., 2022), IPAH PAECs (Rhodes et al., 2015) and known PAH genes (Reyes-Palomares et al., 2020); FDR <0.05

| Comparison lists | Genes shared |
| --- | --- |
| KLF6 HPAECs and PAH HPAECs | *ABCD4, ABCF3, ABI3, ABLIM3, ACVRL1, ADAM10, ADAM23, ADAM9, ADAMTS1, ADAMTS4, ADAMTS9, ADD1, ADGRL2, ADGRL4, ADIPOR2, AHNAK, AHNAK2, AKR1C3, ALCAM, ALDH1A1, ALDH9A1, AMOTL2, ANGPTL4, ANKRD1, ANTXR2, ANXA11, AP1G1, AP1M1, AP3S1, ARAP1, ARHGAP18, ARHGAP24, ARHGAP31, ARHGEF15, ARL6IP5, ARMCX6, ARPC5, ARRB1, ASAP2, ASF1A, ATP2B4, ATP6V1A, ATP6V1H, ATP8B1, BAG1, BCL2L1, BET1, BLCAP, BMX, BRI3, BTN3A1, C15orf40, C16orf74, C1R, C1orf43, C1orf52, C3orf14, C8orf58, CALCRL, CALD1, CALHM2, CALU, CARD6, CASP6, CAT, CAV2, CAVIN3, CCDC50, CCDC59, CCDC85B, CCL2, CCN1, CCND1, CCNE1, CD34, CD93, CDC42EP2, CDC45, CDH11, CDH2, CDH5, CDKN1A, CDKN1B, CDKN2A, CEBPB, CENPK, CENPU, CHCHD2, CHPF2, CHST14, CHURC1, CIP2A, CLCN2, CLEC14A, CLIC4, CLUH, COL4A1, COL5A1, COPG1, COX20, CPNE7, CRABP2, CRIM1, CRIP2, CTDSP1, CYBC1, DAB2, DAG1, DAZAP2, DDAH1, DDAH2, DDX11, DDX23, DGKA, DGLUCY, DHH, DIPK1B, DKK1, DNAJB11, DNAJB4, DNAJC19, DOLK, DPY19L1, DRG2, DYNC1I2, DYNLL1, DYSF, EBNA1BP2, EDN1, EFHD2, EGLN2, EIF1AX, EIF2A, EIF3F, ELOF1, ELOVL1, EMP1, ENC1, ENDOD1, ENG, EPHB4, EPS8L1, ERCC2, ERG28, ERLEC1, EXOSC10, EXOSC5, F2R, FADD, FADS1, FAM107B, FAM124B, FAM171A1, FAM20C, FAM210B, FAM43A, FANCI, FASTKD1, FBN1, FEZ2, FHL3, FLI1, FLNB, FLNC, FOXC2, FOXRED2, FRMD4A, FST, FSTL1, FXR1, FZD4, GALNT1, GAS2L1, GASK1B, GBP3, GEMIN5, GIMAP8, GJA5, GNG11, GNG12, GOLGA2, GORASP1, GPR4, GPRC5A, GPX8, GRAP, GRB14, GTF2IRD2B, HDAC7, HDAC8, HDDC2, HDDC3, HERC5, HHIP, HHIP-AS1, HMGN3, HMGN4, HNRNPA0, HNRNPC, HNRNPLL, HNRNPM, HOMER3, HOXB4, IARS1, ICAM1, ID1, IDH1, IDH3B, IFIT3, IFNAR2, IGBP1, IGF2BP3, IKBKE, IL13RA1, IL1RL1, IL4R, ILF2, INAFM2, IQGAP1, ITGA2, ITGA5, ITGA6, ITGAV, ITGB1, ITGB3, ITGB3BP, ITGB5, JKAMP, JUP, KAT7, KCTD12, KDR, KHNYN, KHSRP, KLF2, KLF3, KLF6, KRCC1, L3HYPDH, LAMA5, LASP1, LDB2, LENG8, LEPROT, LFNG, LIPG, LMO2, LOX, LPCAT2, LRRFIP1, LSM8, LUC7L3, LYL1, LYPLA1, MACF1, MACROH2A1, MAGED1, MALL, MAN2B1, MAP3K11, MAP3K6, MAP7D1, MARCHF4, MARCKS, MARCKSL1, MARK4, MATR3, MAX, MCM7, MDFI, MDM2, MED16, MELTF, METAP2, MFSD11, MGAT2, MIR22HG, MKNK2, MLF1, MLLT1, MLLT11, MMP2, MMRN2, MPRIP, MSANTD3, MTFR1L, MTRES1, MVD, MVP, MYBL2, MYCBP2, MYCT1, MYOF, NAGK, NCAPG2, NDRG1, NEK6, NFIB, NIBAN2, NLRP1, NNMT, NOP53, NOP58, NORAD, NOS3, NRAS, NRCAM, NT5E, NTN4, NUP85, OLA1, OSGIN1, OSGIN2, OSTF1, P3H4, PABPC1, PAM, PAPSS2, PARN, PARP1, PARP4, PCLAF, PCMTD1, PDGFA, PDLIM4, PEAR1, PFN2, PGF, PHLDB1, PIAS3, PICALM, PIEZO2, PKIG, PLEKHG5, PLEKHO1, PLOD2, PLPP3, PLXNB2, PLXND1, PMP22, PNMA1, PNN, PNPLA8, PODXL, POLD4, POMT1, PPP1R37, PREX1, PRKACB, PRKAR1A, PRNP, PRPF8, PRSS23, PTGFRN, PTP4A2, PTPN12, PTPRB, PTPRE, PTTG1, PWP1, RAB32, RAB43, RAB6A, RAI14, RALB, RAMP2, RASA3, RASIP1, RBMS1, RECK, REXO2, RGS4, RHOB, RNF14, RNF141, RNF146, RNF187, RNPEPL1, ROBO4, RPF1, RPIA, RPS6KA2, RRAS, RRBP1, RRM2, RSAD2, RTN4, S1PR1, SAE1, SAFB2, SCAMP4, SCML1, SCRN1, SEC14L1, SEH1L, SEL1L3, SENCR, SEPTIN9, SERINC3, SERPINB8, SERPINE2, SET, SFXN3, SH2D3C, SH3TC1, SHC1, SHISA5, SIDT2, SLC10A3, SLC16A3, SLC19A1, SLC25A13, SLC2A3, SLC35A5, SLC35E2B, SLC38A2, SLC39A13, SLC39A14, SLC47A1, SLC9A3R2, SLFN12, SMARCD2, SMIM14, SMPD1, SMTN, SMURF2, SNHG7, SNRPF, SOX17, SOX18, SPIDR, SPOCK1, SPRY2, SPRYD3, SPTLC2, SQLE, SRD5A3, SRGN, SRRT, SRSF1, SS18, SSBP3, STAU2, STC2, STEAP3, STIMATE, STING1, STMP1, STOM, SUCLA2, SUN2, SWAP70, SYNPO, TADA3, TASOR, TAX1BP3, TBC1D15, TCEAL9, TCIM, TECPR1, TEK, TGFBR2, THBD, THBS1, THOC6, TLN1, TLR4, TM2D2, TM4SF1, TM6SF1, TM9SF1, TMBIM1, TMBIM4, TMEM14C, TMEM167B, TMEM18, TMEM250, TMEM263, TMEM30A, TMEM33, TMEM54, TMEM87B, TMSB4XP4, TNFAIP2, TNFAIP3, TNFAIP8L3, TNFRSF1A, TNFRSF21, TNFSF4, TNIK, TNS1, TNS2, TOM1, TOP2B, TPM1, TPRG1L, TRIM27, TRIM5, TRIM8, TRIP6, TRNP1, TSKU, TSPAN14, TSPAN15, TSPAN17, TSPAN31, TSPAN9, TTL, TTLL12, TTYH3, TUBA1A, U2AF1, U2AF2, UBA5, UBE2C, UBE2J1, UPF3B, USP22, USP47, UTP4, VASP, VEPH1, VIM-AS1, VPS25, VSIR, WDFY1, WDR12, WDR13, WDR6, WSB1, XIST, YWHAG, YWHAZ, ZBTB7B, ZDHHC2, ZNF480, ZNF561, ZNRF1, ZRANB2* |
| KLF6 HPAECs and PAH ECFCs | *ABCA3, ABCF1, ABLIM3, ACO1, ACOT7, ACOT8, ACTA2, ADA, ADAMTS1, ADAMTS4, ADCK2, ADCY4, ADD1, ADGRA3, ADM, ADORA2A, AEN, AGPAT2, AGPAT5, AHCY, AHSA1, AIG1, AIP, AK1, AK2, AKR1A1, AKR7A2, ALDH18A1, ALDH1B1, ALDOA, ALG1, ALKBH7, ALYREF, AMFR, AMPD2, AMZ2, ANAPC7, ANGPTL4, ANKRD9, ANLN, ANO10, ANO6, ANP32B, ANP32E, ANTKMT, ANXA11, ANXA6, AP2A1, AP2S1, AP3D1, APLN, APOBEC3C, APOBEC3G, APOE, APOL2, APOL3, APRT, ARF6, ARFGAP3, ARFRP1, ARHGAP10, ARHGDIB, ARHGEF2, ARL2, ARL4D, ARMCX2, ARPC1A, ARPC1B, ARPC4, ARRDC3, ARSJ, ASF1A, ASS1, ATIC, ATOX1, ATP13A2, ATP5F1D, ATPAF1, ATXN10, AURKAIP1, AURKB, AVEN, AXL, B4GALT2, BAIAP2, BAZ1B, BCAR1, BCAT2, BCCIP, BCL10, BCL3, BCYRN1, BEX3, BEX4, BHLHE40, BIN1, BIRC5, BLOC1S6, BNIP2, BOD1, BOK, BOP1, BPGM, BRAT1, BRD9, BTG2, BTN2A2, BTN3A2, BUD31, C19orf33, C1QBP, C1R, C1orf35, C20orf27, C4orf48, CADPS2, CALCOCO2, CALHM2, CALM3, CAMK1, CANX, CAPZA1, CARD10, CARD16, CARS2, CAV1, CAVIN1, CAVIN2, CBX6, CC2D1A, CCAR1, CCDC174, CCDC25, CCDC28B, CCDC34, CCDC50, CCDC85B, CCDC86, CCM2, CCN2, CCNA2, CCNB1, CCNI, CCNQ, CCT2, CCT3, CCT4, CCT6A, CCT7, CCT8, CD47, CD82, CDC20, CDC25A, CDC42EP2, CDC42EP4, CDCA5, CDIPT, CDK1, CDK11B, CDK2AP2, CDKN2A, CDR2L, CDT1, CDV3, CEBPB, CEBPD, CEBPZ, CENPB, CENPV, CEP131, CEP55, CGNL1, CHCHD10, CHCHD3, CHD1L, CHD3, CHD4, CHID1, CHMP4A, CHORDC1, CHP1, CHTF8, CIAO2B, CIRBP, CISD3, CIZ1, CKS1B, CKS2, CLDN11, CLEC11A, CLIC1, CLNS1A, CLPB, CLPP, CLPTM1L, CLSPN, CLTA, CLTC, CMSS1, CNN2, CNN3, COA6, COL1A2, COL6A2, COMMD9, COPB2, COPG1, COPS6, COQ5, CORO1C, CRACR2B, CRELD2, CRK, CSE1L, CSNK1A1, CTDSP1, CTNNAL1, CTNNBL1, CTR9, CTSB, CTSF, CTSK, CTU2, CUL4B, CWC27, CX3CL1, CXCL1, CXCL2, CXCL3, CXCL8, CXXC5, CYBA, CYLD, CYP51A1, DARS1, DBNDD2, DCPS, DDAH1, DDAH2, DDHD2, DDIT3, DDIT4, DDRGK1, DDX11, DDX17, DDX18, DDX21, DDX27, DDX41, DDX49, DDX54, DEAF1, DENND1A, DEPP1, DESI1, DFFA, DGCR6L, DGKZ, DHH, DHPS, DHRS3, DHX9, DIAPH1, DIPK1B, DKC1, DKK1, DNAJA1, DNAJA4, DNAJB1, DNAJB4, DNAJC1, DNAJC3, DNAJC4, DNASE1L1, DOHH, DOK1, DPP7, DPYSL2, DRAM2, DROSHA, DTL, DTX3L, DUSP1, DUSP5, DUSP7, DYM, DYNC1H1, DYNC1I2, DYNLL1, DYSF, EBP, ECH1, ECHDC2, ECI1, ECI2, ECSIT, EEF1A1, EEF1AKMT4, EEF1B2, EEF1D, EFEMP2, EGLN2, EHBP1L1, EHD1, EHD2, EIF2B3, EIF2S2, EIF2S3, EIF3B, EIF3C, EIF3I, EIF3J, EIF3K, EIF3L, EIF3M, EIF4A1, EIF4A2, EIF4A3, EIF4EBP1, EIF4G1, EIF5A, EIF5B, ELP1, ELP2, ENAH, ENDOG, ENO1, ENO2, ENTPD6, EPRS1, ERCC1, ERCC2, ERG, ESF1, ESM1, ESYT1, ETFB, ETS2, EVA1B, EXOSC3, EXOSC4, FADS3, FAF1, FAM111A, FAM120A, FAM162A, FAM168B, FAM174C, FAM50A, FANCG, FARSA, FASN, FAU, FBL, FBXL5, FBXO22, FDFT1, FDX2, FDXR, FECH, FGFR1OP2, FIS1, FKBP11, FKBP2, FKBP4, FLI1, FLII, FLNB, FLNC, FLOT1, FLOT2, FOXRED1, FSCN1, FST, FSTL3, FTH1, FTL, FTSJ1, FZD4, G6PD, GABARAPL1, GAK, GALK1, GAMT, GAPDH, GAS5, GBGT1, GBP2, GCDH, GEMIN4, GEMIN6, GFM1, GIT1, GMPPB, GNL1, GOLGA4, GPATCH4, GPI, GPR137, GPRC5A, GPS1, GRAMD1A, GRB14, GRWD1, GSDMD, GSN, GSPT1, GTF3A, GTF3C5, GTPBP2, GTPBP6, GUCD1, H1-0, H2AC6, H2AX, HAGHL, HAUS4, HAUS7, HAX1, HCFC1R1, HDDC3, HDGF, HDGFL2, HDLBP, HEBP1, HEG1, HES1, HES4, HEXIM1, HGH1, HHIP-AS1, HLA-E, HMGN2, HMOX1, HNRNPC, HNRNPDL, HNRNPU, HPRT1, HSP90AA1, HSP90AB1, HSP90B1, HSPA1A, HSPA1B, HSPA5, HSPA6, HSPA8, HSPB8, HSPD1, HSPH1, HTATIP2, HTATSF1, HYI, IARS1, ID1, ID3, IER5, IER5L, IFI27L2, IFITM3, IFT57, IGBP1, IGF2BP2, IGF2BP3, IGFBP4, IGFBP5, IGFBP6, IKBIP, IKBKE, IL33, IL6, ILF2, ILK, ILRUN, IMMT, IMP4, IMPDH1, IMPDH2, INF2, INPP5A, INPP5D, INSIG1, INTS1, INTS13, IPO11, IPO4, IPO5, ISYNA1, ITGA3, ITGAV, ITGB1BP1, ITGB4, ITPR3, ITPRIP, JAG1, JKAMP, JPT2, JTB, JUNB, KANK1, KANK3, KARS1, KCTD17, KCTD9, KDM1A, KDM5A, KIAA0930, KIF5B, KLF16, KLF2, KLF6, KLHDC2, KPNB1, KRT18, KRT19, KRT7, KRT8, LAMA5, LAMB2, LAMTOR2, LARS1, LDHA, LDHB, LDLRAP1, LETM1, LIMS2, LINC01013, LINC01235, LIPG, LMAN2, LMF2, LMNA, LONP1, LONP2, LRFN4, LRP3, LRPPRC, LRRC32, LRRC42, LRRC59, LRRC70, LSM2, LSM4, LTBP3, LTBP4, LUC7L, LUC7L3, LY6E, LYAR, LYRM1, MAD1L1, MAEA, MAGED1, MALAT1, MALL, MAOA, MAP1B, MAP2K3, MAP4, MAP4K2, MAP4K4, MAP7D1, MAP7D3, MAPK11, MAPK12, MARCHF2, MAZ, MBD3, MCM5, MCRIP2, MCTS1, MDH2, MDM2, ME2, ME3, MED16, MEPCE, MEST, METAP2, METRN, METTL26, METTL7A, MFSD10, MFSD12, MGLL, MICALL1, MIF, MIR22HG, MIR663AHG, MKNK1, MKNK2, MLKL, MLLT1, MLLT11, MMP17, MMS19, MON1A, MPRIP, MPST, MRPL11, MRPL12, MRPL17, MRPL35, MRPL37, MRPL41, MRPL53, MRPL54, MRPS15, MRPS2, MRPS24, MRPS25, MRPS34, MRPS5, MRPS6, MRPS9, MSH6, MSL3, MSMO1, MT1E, MT2A, MTCH2, MTHFD1, MTHFS, MVD, MVK, MYBL2, MYH9, MYL6, MYL6B, MYO1C, MYO1D, MYO5A, MYOF, NAA10, NAA15, NAAA, NADK, NAE1, NANS, NAT10, NCAPG, NCAPH2, NCL, NCOA4, NCOR1, NDUFB1, NDUFB5, NDUFS3, NDUFS8, NDUFV1, NEDD9, NELFB, NELFE, NEMF, NENF, NES, NEXN, NFE2L3, NFIB, NFIC, NFKB1, NFKBIA, NFYB, NGRN, NIPSNAP1, NLE1, NME2, NME3, NMT2, NNMT, NOC2L, NOP14, NOP53, NOP9, NOS3, NPAS2, NPDC1, NPM1, NPM3, NPR1, NQO1, NR2F2, NR2F6, NRARP, NRG1, NSMCE1, NSMF, NT5C2, NTHL1, NUCB2, NUCKS1, NUDC, NUDT21, NUDT5, NUP155, NUP205, NUP214, NUP37, NUP43, NUP62, NUP88, ODF2, ODF2L, OGFR, OLA1, OSGIN2, PABPC1, PABPC4, PAFAH1B2, PAICS, PALM, PARD3, PARP10, PCDH10, PCDH12, PCM1, PDCD11, PDCL, PDGFB, PDHA1, PDLIM1, PDLIM3, PDLIM7, PDXK, PELP1, PES1, PFDN4, PFDN5, PFN2, PGLS, PGM1, PGP, PHACTR2, PHETA2, PHF19, PHLDA2, PHLDA3, PHLDB2, PHYKPL, PIEZO1, PIGW, PIH1D1, PITHD1, PITPNM1, PKD1, PLEKHO2, PLIN2, PLIN3, PLOD2, PLXND1, PMVK, PNMA1, PNPLA6, PNRC1, PODXL, POLD2, POLD4, POLE4, POLR1B, POLR1E, POLR2K, POLR3D, POLR3E, POLR3GL, POLRMT, POP7, PPA1, PPAN, PPIB, PPID, PPIE, PPIF, PPM1G, PPP1CB, PPP1R12C, PPP1R16A, PPP1R35, PPP1R37, PQBP1, PRADC1, PRCC, PRDX5, PREB, PRELID1, PRELID3B, PRKAR2A, PRKCH, PRKD2, PRKDC, PRMT5, PRMT7, PRPF19, PRPF38A, PRPF40A, PRPF8, PRR7, PSMB10, PSMB9, PSMD4, PSME4, PTDSS1, PTDSS2, PTEN, PTGS1, PTMA, PTMS, PTOV1, PTP4A2, PTPRB, PTPRK, PTRH1, PTTG1, PTX3, PYCR1, PYGL, QPCT, QSOX1, QTRT1, RAB11FIP5, RAB13, RAB34, RAB8A, RABAC1, RABL6, RAC2, RAC3, RALA, RALB, RAN, RANGAP1, RAP1GDS1, RASIP1, RBM38, RCC1, RCC1L, RCN1, RDH11, REEP3, RELL1, RER1, REXO4, RFLNB, RFTN1, RGS3, RGS4, RHOC, RHOD, RNASEH2A, RNASEH2C, RNF144B, RNF167, RNF181, RNF19B, RNF213, RNPEPL1, RPE, RPL10A, RPL12, RPL13A, RPL28, RPL3, RPL31, RPL32P29, RPL34, RPL37, RPL37A, RPL4, RPL7A, RPP25, RPS3, RPS9, RPUSD4, RRAS, RRM2, RRP12, RSF1, RUFY1, RUVBL2, SAC3D1, SAMD1, SAMHD1, SAR1B, SBF1, SCAND1, SCAP, SCARB1, SCRIB, SDC1, SDHB, SEC22B, SEC23B, SELE, SELENOM, SEMA3F, SEMA4B, SEPTIN11, SEPTIN5, SEPTIN9, SERBP1, SERF2, SERP1, SERPINH1, SF3B2, SF3B5, SF3B6, SFXN4, SGTA, SH3TC1, SHC1, SHISA5, SHMT2, SIAE, SIAH2, SIL1, SLC16A1, SLC1A5, SLC20A1, SLC20A2, SLC25A3, SLC25A39, SLC25A46, SLC25A5, SLC27A3, SLC27A4, SLC35A5, SLC35E1, SLC38A2, SLC39A14, SLC43A3, SLC4A2, SLC50A1, SLC9A1, SLC9A3, SLC9A3R2, SLFN11, SLIT2, SMAGP, SMC4, SMTN, SNAP29, SNCG, SNHG29, SNHG7, SNRNP40, SNRPN, SNX5, SNX6, SOCS3, SOD1, SON, SOX12, SOX17, SOX4, SPATS2, SPATS2L, SPNS2, SPOCK1, SPOP, SPTAN1, SQLE, SREBF2, SREK1, SREK1IP1, SRP72, SRPRB, SRRM1, SRSF4, SRSF5, SRSF9, SS18, SSBP4, SSR1, SSR3, SSR4, SSRP1, SSU72, ST6GALNAC4, STARD10, STK16, STK25, STK38L, STMP1, STOML2, STUB1, SUGT1, SULT1A1, SUMF2, SUPT16H, SUPV3L1, SURF1, SURF2, SWAP70, SYMPK, SYNGR2, SYNPO, SYS1, TACC3, TADA3, TAF10, TAF13, TAOK3, TAPBP, TARS1, TASOR, TATDN2, TBC1D1, TBC1D10B, TBC1D2, TBCA, TBCD, TBL3, TCF19, TCIRG1, TCP1, TEAD4, TFE3, TFRC, TGM2, THAP4, THAP7, THBD, THG1L, THOC1, THOC5, THOP1, TIMM17B, TIMM50, TIMM9, TIMP1, TLE2, TLN1, TM2D2, TM4SF18, TMEM109, TMEM115, TMEM165, TMEM167B, TMEM219, TMEM250, TMEM30A, TMEM44, TMEM54, TMSB10, TMUB1, TNFAIP2, TNFAIP3, TNFAIP8L1, TNFRSF14, TNFRSF1A, TNFRSF6B, TNFSF18, TNIK, TNIP1, TNIP2, TOP1, TOP1MT, TOP2A, TOP2B, TOP3A, TP53I11, TP53I13, TPI1, TPP2, TPRN, TPT1, TPX2, TRABD, TRADD, TRAF7, TRAP1, TRIM28, TRIM44, TRIM5, TRIM69, TRIP12, TRIP13, TRIR, TRMT1, TRMT112, TRMT2A, TRNP1, TSC22D1, TSC22D3, TSC22D4, TSPYL1, TSR3, TST, TTC27, TTC7B, TTC9C, TTLL12, TUBA4A, TUBB4B, TUBGCP2, TUFM, TWF2, TWIST2, TXLNA, TXNL1, TXNL4A, TXNRD2, TYMP, TYMS, TYRO3, UBB, UBC, UBE2L6, UBE2M, UBE2S, UBL4A, UBQLN4, UBR4, UBTF, UFC1, UFSP2, UGDH, UGP2, UQCRC1, USO1, USP1, USP16, USP18, USP39, UTP14A, VAMP2, VAMP5, VARS1, VEGFB, VPS18, VPS51, VTI1B, WARS1, WASHC4, WDR18, WDR4, WDR54, WDR77, WTIP, WWP1, XBP1, XRCC5, YBX3, YLPM1, ZBTB7A, ZC3H15, ZC3HAV1, ZCCHC3, ZFR, ZFYVE19, ZMAT2, ZMAT3, ZNF185, ZNF358, ZNF561, ZNF581, ZNF593, ZNF702P, ZNF787, ZNHIT1, ZPR1* |
| KLF6 HPAECs and IPAH PAECs | *ABHD12, ABHD16A, ABI1, ACVR1, ADAMTSL1, AHNAK, ARIH2, ARL4A, ARMCX3, ATP6V1H, B3GALNT1, C1R, CCL2, CEBPB, CGNL1, CHRD, COL4A1, COL4A2, COL6A2, CXCL11, DARS2, DDX60, DERL1, DHH, DKK3, DNAJB9, DSP, DUSP1, EFNA1, ESRRA, ETS1, EXOSC4, FDXR, FOS, FSTL3, GOLGA7, GRAP, GTPBP2, HERC5, HERPUD1, HES1, HES4, HMGB3, HSD17B12, HSPA1A, IER5L, IFI30, IFI35, IFIT3, IGFBP7, ILF3, ITFG1, JUNB, KANK3, KIAA0930, LAMB2, LAPTM5, LOX, LPIN1, LTB, LTBP1, M6PR, MAGED2, MAP1LC3A, MAX, MCOLN1, MDM2, MMP17, MRC1, MX2, NBEAL2, NCAPD2, NDFIP2, NFE2L1, NFKBIA, NME1-NME2, NOS3, NRARP, NSG1, NSMAF, NT5C3A, OAS1, ODC1, PAM, PANX1, PFAS, PKIG, PLPP1, PLVAP, PNPT1, PTPRE, RGCC, RHOB, RND3, RNF114, RNF146, RPS27, RTP4, SAMD9, SAT1, SDF2, SERINC1, SGCE, SH2D3C, SLC25A25, SLC35A2, SLC35E1, SLU7, SMOX, SNCA, SNHG3, SP110, SPARC, SPNS2, SPSB1, STAT1, SULF2, SULT1A1, TFIP11, TMEM167B, TMEM87A, TMEM9B, TNFAIP1, TNFRSF21, TNFSF10, TSC22D1, TSG101, TSPAN15, TUBA1A, TYMP, UGCG, UNC13D, UNC93B1, YPEL5, ZNF185* |
| KLF6 HPAECs and known PAH genes | *ABCA3, ABCD4, ACKR3, ACTA2, ACVRL1, ADORA2A, AGPAT2, APOE, ARAP1, ARHGAP31, ATPAF2, BANF1, CAV1, CDIPT, CERS1, CITED2, CLCN7, COL1A2, CRTAP, CYB5D2, DCTN4, DDAH1, DOCK6, DPP9, DSP, DVL3, EDN1, ELK3, ENG, EOGT, EPAS1, EPHB4, ERAP1, ERG, FAS, FBN1, FGFR3, FOS, FOXM1, GIT1, GJA1, GNB5, HLA-DPB1, HSPG2, JAG1, KIAA0319L, KRT18, KRT8, LIPA, MGP, MLX, NFIX, NPRL3, P3H1, PAM16, PARN, PDSS1, PPIB, SARS2, SERPINE1, SERPINH1, SLC37A4, SLC9A3, SMAD4, STAT1, STN1, TCIRG1, TGFB1, THBS1, TLR4, TNFRSF1A, TOPBP1* |

**Table S4: List of KLF6-regulated DEGs shared with non-plexiform lesions- and plexiform lesions-specific DEGs.**

| Comparison List | Overlapping genes |
| --- | --- |
| KLF6 vs Non-plexiform lesions | *ABCB10, ACADM, ACSL4, ADAM9, AKT1, ALDOA, ANAPC5, ANXA5, APBB2, APOE, ARF3, ARHGAP29, ARHGEF1, ARHGEF15, ARPC1B, ATF4, ATP2A3, ATP2C1, ATP5F1B, ATP5F1D, ATP5MC3, BCAR1, BCL2L1, BCR, BOP1, BRK1, BTG1, BZW1, C17orf49, C5orf24, CCNI, CD63, CD83, CDC42EP4, CDH11, CDK11A, CDK2AP1, CHIC2, CHTOP, CLIC1, CLIP1, CLU, COA4, COL1A2, COL4A1, COL6A2, CRTAP, CST3, CTSB, CTSD, DAZAP2, DCAF11, DDX60, DHX36, DIPK2A, DNPH1, ECH1, ECHDC2, EEF1B2, EEF1D, EFEMP1, EID1, EIF3K, EIF3L, EIF4B, ELOC, EPRS1, ERCC1, ETFB, F11R, FAM111A, FAM43A, FASN, FAU, FGFRL1, FH, FKBP11, FKBP1A, FOSL2, GALNT2, GBA2, GIMAP2, GIMAP7, GMPR2, GPR108, GPR4, GRINA, GTF2I, HARS1, HBEGF, HES1, HIF1A, HIGD1A, HLA-DPA1, HLA-DPB1, HMGB1, HMGN2, HNRNPA0, HNRNPA3, HNRNPAB, HNRNPUL2, HSD17B12, HSP90AB1, HSPA8, HSPB1, HSPB8, ID1, IFITM3, IGFBP4, IGFBP7, IL13RA1, IL6ST, ILF3, ILVBL, ING5, ITGB1, ITSN1, JMJD8, KLC2, KLF2, LAMB1, LASP1, LGALS1, LMAN2, LYRM2, MALL, MAP4, MAPK9, MATR3, MAZ, MEGF6, MGLL, MKNK2, MORF4L1, MPRIP, MPZL2, MRFAP1L1, MRPS25, MT1A, MT1E, MT2A, MYCT1, MYH9, MYL6B, NASP, NDUFV2, NEXN, NFE2L1, NFKBIA, NIBAN2, NID1, NME2, NNMT, NPM1, NQO2, NT5E, OST4, P4HB, PABPC1, PABPC4, PDIA3, PDLIM1, PFDN5, PGM2, PLAT, PLEC, PLEKHG5, PLK2, PMEPA1, PMP22, POLR2E, PPP1R2, PSAT1, PSMD14, PSMD3, PTMA, PTMS, PTP4A2, PTPN1, PYM1, QSOX1, RAB1A, RAB34, RAN, RANBP2, RNASEK, RND3, RNF213, RPL10A, RPL12, RPL15, RPL28, RPL31, RPL34, RPL36AL, RPL37, RPL37A, RPL7A, RPS11, RPS18, RPS27, RPS29, RPS3, RPS6KA2, RPS9, RTN4, S1PR1, SAMD4A, SDHB, SELENOT, SEPTIN9, SERF2, SF3B6, SGK1, SH3BGRL3, SHC1, SLC25A13, SLC25A3, SLC25A5, SLC35D2, SMARCA2, SON, SRPRA, SRSF5, SRSF9, STAG1, STK25, STOML2, SUMF2, SYNGR2, SYPL1, TAPBP, TBC1D15, TFPI, TGFBI, THBS1, THOC1, THYN1, TM9SF3, TMED10, TNFSF10, TNIP1, TNKS1BP1, TNPO1, TNRC18, TNS2, TRIM22, TSN, TUBA1C, TXNIP, UBA1, UBXN8, UFC1, UFSP2, UQCRC1, USP22, VAMP2, VAPA, VCAN, VWF, XRCC6, YWHAZ, ZFAND5, ZFP36L2, ZFTA, ZYX* |
| KLF6 vs Plexiform lesions | *ACTN1, ACTR2, ACTR3, ADAM9, ADAMTS1, ADAR, ADCY3, AGPAT2, AIF1L, AKAP12, AKIRIN1, AKIRIN2, AKT1, ALDH1A1, ALDOA, AMFR, ANGPTL4, ANKH, ANP32B, ANPEP, ANTXR1, ANXA2, ANXA5, AP1M1, AP2S1, AP3S1, APOE, APP, ARF1, ARF3, ARF4, ARGLU1, ARHGDIA, ARHGDIB, ARHGEF1, ARHGEF7, ARL6IP1, ARL6IP5, ARPC1B, ARPC4, ATF4, ATL3, ATOX1, ATP2B4, ATP5F1B, ATP5F1D, ATP5MC3, ATP6AP2, AXL, BANF1, BCAR1, BCL2L1, BCL7C, BGN, BRK1, BTG1, BZW1, C1QTNF1, C1R, C1orf43, C4orf3, CALD1, CALM2, CALM3, CALU, CANX, CAVIN3, CBX6, CCNG1, CCNI, CCNY, CCT3, CCT4, CCT8, CCZ1, CD164, CD276, CD59, CD63, CD9, CD99, CDC42, CDC42BPB, CDC42EP4, CDH11, CDK11A, CDK13, CDK2AP1, CDK9, CDKN1B, CDV3, CEBPB, CENPB, CFH, CFL2, CHCHD10, CHCHD2, CHD4, CHIC2, CHSY1, CLDN5, CLIC1, CLIC4, CLIP1, CLTA, CLU, CMPK1, CNBP, CNIH1, CNN3, COL18A1, COL1A2, COL4A1, COL4A2, COL5A1, COL5A2, COL6A2, COL8A1, CORO1C, CRIM1, CRTAP, CS, CSNK2B, CSPG4, CSRP1, CST3, CTSB, CTSD, CTSK, CX3CL1, CYBRD1, DAPK3, DARS1, DAZAP1, DAZAP2, DBNDD2, DDAH2, DDIT4, DDX6, DEK, DKK3, DLGAP4, DOCK1, DOCK9, DPP7, DVL1, DYNLL1, ECE1, ECH1, EEF1B2, EEF1D, EFEMP1, EHBP1L1, EHD2, EHD4, EID1, EIF1AX, EIF3K, EIF3L, EIF4A1, EIF4B, EIF5A, ELOC, ENG, ENO1, ENO2, ESAM, ESD, EVA1B, EXOSC3, FAM20C, FAU, FBLIM1, FBN1, FBXL5, FBXO11, FIS1, FKBP11, FKBP1A, FKBP2, FKBP9, FLOT1, FN1, FNDC3B, FOSL2, FSTL1, FTH1, GABARAP, GALNT2, GATAD2A, GBP2, GET3, GIMAP7, GINM1, GJA5, GLG1, GLUD1, GNAI2, GNB1, GNG12, GNG5, GOLGB1, GOLPH3, GPI, GRB2, GRINA, GRK2, GRN, GSN, GSPT1, GSTO1, GTF2I, H2AJ, H3-3B, HDLBP, HES1, HEXB, HIF1A, HIGD1A, HLA-DPA1, HLA-DPB1, HLA-F, HMGB1, HMGB2, HMGN2, HNRNPA0, HNRNPA3, HNRNPAB, HNRNPDL, HNRNPF, HNRNPUL1, HNRNPUL2, HSD17B12, HSP90AB1, HSP90B1, HSPA8, HSPB1, HSPG2, HTRA1, ICAM2, ID1, IFITM3, IFNGR1, IGFBP4, IGFBP5, IGFBP7, IL13RA1, IL6ST, ILF3, ILK, INF2, IQGAP1, IRF2BP2, ISCU, IST1, ITGA5, ITGAV, ITGB1, ITGB5, ITPRIP, KDM2A, KHSRP, KIAA2013, KLF2, KLHDC3, KPNB1, LAMA4, LAMA5, LAMB1, LAMC1, LAPTM4A, LAPTM5, LASP1, LDHA, LGALS1, LMAN2, LRP10, LRRC32, LRRFIP1, LSM14A, LTBP1, LTBP2, LTBP3, LYRM2, MACROH2A1, MAP1B, MAP3K6, MAP4, MAP4K4, MARCKS, MATR3, MAZ, MBNL1, MED15, MEF2A, METAP2, METRNL, MGLL, MGP, MICAL2, MICALL2, MIF, MINK1, MMP14, MMP2, MORF4L1, MORF4L2, MPRIP, MPZL2, MRPL3, MRPS25, MSN, MT1A, MT1E, MT2A, MTA1, MVP, MXD4, MYH9, MYL12A, MYL6, NASP, NCL, NDRG1, NDUFA10, NDUFB4, NDUFV2, NEDD9, NEK7, NES, NEXN, NFE2L1, NFIC, NFIX, NFKBIA, NGRN, NHP2, NIBAN2, NID1, NISCH, NLRP1, NME2, NNMT, NOP53, NPDC1, NPM1, NPRL3, NPTN, NR2F2, NUCB1, NUTF2, OLA1, OS9, OST4, OTUB1, P4HB, PABPC1, PABPC4, PACSIN2, PAPSS2, PCBP1, PCDH1, PDCD5, PDCD6IP, PDIA3, PDLIM1, PDLIM3, PDLIM5, PDLIM7, PEAK1, PFDN5, PFKP, PHC2, PHLDA1, PHLDA3, PICALM, PINK1, PLEC, PLEKHO1, PLPP1, PLS3, PLTP, PLXND1, PML, PMP22, PODXL, POGLUT3, POLR2E, POSTN, PPA1, PPIB, PPP1CB, PPP1R18, PPP1R2, PRDX2, PRKACA, PRKAR1A, PRKCH, PRNP, PRRC2B, PRSS23, PRXL2B, PSAT1, PSMB1, PSMD3, PSMD8, PTK2, PTMA, PTMS, PTP4A2, PTPN1, PTTG1IP, PUF60, PXDC1, RAB13, RAB1A, RAB34, RAB6A, RABAC1, RAN, RANBP1, RANBP2, RARA, RELL1, REXO2, RGS5, RHOC, RIPOR1, RNASEH2C, RNASEK, RND3, RNF103, RNF181, RPL10A, RPL12, RPL13A, RPL15, RPL28, RPL31, RPL34, RPL36AL, RPL37, RPL37A, RPL4, RPL7A, RPS11, RPS18, RPS27, RPS27L, RPS29, RPS3, RPS6KA2, RPS9, RRAS, RRBP1, RTN4, S100A11, S100A13, S100A16, S100A6, S1PR1, SARAF, SCAMP2, SCARA3, SDHB, SDHC, SELENOM, SELENON, SENP6, SEPTIN9, SERBP1, SERF2, SERINC3, SERPINE1, SERPINE2, SF3A1, SF3B6, SFXN3, SGK1, SH3BGRL3, SH3GLB1, SH3GLB2, SHC1, SKI, SLC16A3, SLC25A3, SLC39A13, SLC39A7, SLC7A2, SLCO2A1, SLIT2, SMARCE1, SNHG32, SNRNP27, SNRPG, SNRPN, SON, SPARC, SPATS2L, SPCS3, SPTAN1, SRGN, SRP14, SRPRA, SRSF11, SRSF5, SRSF9, SSB, SSBP3, SSR3, STAB1, STMN1, STOM, SUMO2, SUN1, SYNGR2, SYNPO, SYPL1, TACSTD2, TAF15, TAGLN, TAMALIN, TAPBP, TBCA, TCEAL1, TCEAL4, TFPI, TGFB1, TGFBI, TGM2, THBS1, TIMP1, TIMP2, TINAGL1, TLK2, TLN1, TM9SF3, TMBIM1, TMED10, TMEM109, TMEM127, TMEM165, TMEM259, TMEM50A, TMEM9, TMSB10, TMSB4X, TNFSF10, TNIP1, TNKS1BP1, TNRC18, TNS1, TNS2, TPM1, TPP1, TRIM22, TRIM28, TRIM8, TRIO, TRIP12, TRIR, TRMT112, TSC22D3, TSPAN14, TUBA1C, TUBB, TUFM, TXNIP, TXNRD1, TYMP, UBA1, UBE2I, UBE2L6, UBXN6, UFC1, UGP2, UQCRC1, USP22, UTRN, VAMP2, VAPA, VCAN, VDAC1, VEGFB, VIM, VKORC1, VWF, WWTR1, XRCC6, YBX3, YIPF3, YWHAH, YWHAZ, ZBTB7A, ZC3H11A, ZFAND5, ZFP36L2, ZNHIT1, ZYX* |
| KLF6 vs all PAH vessels (Plexiform+non-plexiform) | *ABLIM3, ABR, ACTN1, ACVRL1, AIF1L, ALDOA, ANKH, ANKRD39, ANP32B, ANTXR1, AP2S1, AP3S1, ARF1, ARGLU1, ARHGDIA, ARHGEF15, ARHGEF7, ARL2, ARPC2, ARPC4, ATL3, ATP6AP2, ATP8B1, AXL, BET1L, BRK1, C5orf24, C7orf50, CALHM2, CALM3, CANX, CAPNS1, CAVIN1, CBR1, CBX6, CCNL2, CD164, CD276, CD59, CDK13, CDK5RAP3, CENPB, CLIC1, CLIC4, CLTA, CNIH1, COL18A1, COL5A2, COL8A1, COPG1, CSNK2B, CSRP2, CTNNB1, DARS1, DBNDD2, DDAH2, DIPK2A, DKK3, DPP7, DUSP7, DYNC1I2, EAPP, ECE1, EEF1D, EFTUD2, EID1, ELAVL1, ENO2, ERCC1, FBLIM1, FBN1, FHL2, FHL3, FIS1, FKBP9, FMNL3, FNDC3B, FOXC2, GADD45A, GALNT2, GBE1, GBP2, GDI1, GINM1, GJA5, GLG1, GNB1, GNPTG, GPR4, GRK2, GSN, H1-0, H2BC5, HBP1, HEG1, HES4, HIGD1A, HMGB2, HMOX1, HNRNPUL1, HTRA1, ICAM2, ICMT, ILVBL, INF2, IPO5, IST1, ITGA10, ITGA5, ITGB1, ITGB5, ITM2B, KIAA2013, KLHDC3, LAMA4, LAMA5, LAMB2, LGALS1, LIMA1, LIMS2, LOX, LRP10, LTBP1, LTBP2, LTBP3, LZTS2, MAP1B, MAP3K11, MAP4K4, MARCHF7, METRNL, MICA, MICAL2, MIF, MMP14, MOB1A, MORF4L2, MPRIP, MRPL3, MRPS21, MRPS24, MSN, MTCH2, MTHFD2, MYH9, MYL6, MYO1D, NAA10, NES, NLRP1, NME2, NPM1, NPRL3, NPTN, NSMF, NUBP2, NUTF2, PANK3, PAPSS2, PCDH1, PDLIM3, PDLIM4, PDLIM5, PGM2, PHLDA1, PHLDA3, PITRM1, PKD1, PLAT, PLEC, PLEKHG5, PLS3, PMEPA1, PPP1CB, PPP1R14B, PPP1R18, PRKACA, PRRC2B, PRSS23, PRXL2B, PTDSS1, PTK2, PTMA, PTOV1, PTP4A2, PTP4A3, PTTG1IP, RAB13, RAB6A, RANBP1, RAPGEF5, RARA, RER1, RHOB, RHOC, RING1, RIOK3, RRAGA, RRAS, RUFY1, S100A16, SARAF, SCARA3, SEC24D, SELENON, SERPINE1, SH3BGRL3, SH3BP5, SH3GLB1, SIRT2, SIVA1, SLC16A3, SLC20A1, SNHG32, SNX9, SOX18, SPARC, SPATS2L, SPTAN1, SRP14, SRSF2, SSB, SSU72, ST3GAL1, STAB1, STK25, SUN2, SYNPO, TACSTD2, TCEAL9, TGM2, TIMP2, TLE1, TLN1, TMBIM1, TPI1, TPM1, TRMT112, TSPAN14, TUBB, TUFM, TXLNA, UBA2, UBE2H, UGDH, UGP2, UROD, VIM, WBP1L, WWTR1, YWHAG, ZBTB7A, ZNF358, ZNFX1* |
| KLF6 vs [Control vs PAH] | *ABI1, ACADM, ACSL4, ACTR2, ADAM9, AKIRIN2, AKT1, ALDOA, ANXA5, APOE, ARF3, ARHGAP29, ARHGDIB, ARHGEF1, ARL6IP1, ARL6IP5, ARPC1B, ATF4, ATP5F1B, ATP5F1D, ATP5MC3, BCAR1, BCL2L1, BGN, BRK1, BTG1, BZW1, C17orf49, CCNI, CCT4, CCT8, CD63, CD83, CDC42EP4, CDH11, CDK11A, CDK2AP1, CDV3, CHCHD2, CHD4, CHIC2, CHTOP, CLIC1, CLIP1, CLTA, CLU, CNOT8, COA4, COL1A2, COL4A1, COL4A2, COL6A2, CRTAP, CSNK2B, CST3, CTSB, CTSD, DAPK3, DAZAP1, DAZAP2, DCAF11, DHX36, DIPK2A, DKK3, DLAT, DVL1, ECH1, ECHDC2, EEF1B2, EEF1D, EFEMP1, EID1, EIF3K, EIF3L, EIF4B, EIF5A, ELOC, ENG, ENO1, ERCC1, FAM43A, FAU, FKBP11, FKBP1A, FNDC3B, FOSL2, FUOM, GALNT2, GIMAP2, GIMAP7, GPI, GPR108, GRINA, GSN, GTF2I, H1-10, HES1, HIF1A, HIGD1A, HLA-DPA1, HLA-DPB1, HMGB1, HMGN2, HNRNPA0, HNRNPA3, HNRNPAB, HNRNPDL, HNRNPUL2, HSD17B12, HSP90AB1, HSPA8, HSPB1, HSPB8, ID1, IFITM3, IFNGR1, IGFBP4, IGFBP7, IL13RA1, IL6ST, ILF3, ILVBL, ING5, IRF2BP2, ITGAV, ITGB1, ITGB5, KLC2, KLF2, LAMB1, LAMC1, LASP1, LGALS1, LMAN2, LRRFIP1, LYRM2, MALL, MAP1B, MAP3K6, MAP4, MAPK9, MATR3, MAZ, MEGF6, MGLL, MGP, MICALL2, MKNK2, MORF4L1, MPRIP, MPZL2, MRFAP1L1, MRPS25, MT1A, MT1E, MT2A, MTHFS, MVP, MXD4, MYCT1, MYH9, MYL12A, MYL6B, NASP, NCL, NDUFV2, NEXN, NFE2L1, NFIX, NFKBIA, NGRN, NIBAN2, NID1, NME2, NNMT, NPM1, NUTF2, OST4, P4HB, PABPC1, PABPC4, PDIA3, PDLIM1, PFDN5, PLEC, PLK2, PLTP, PMP22, POGLUT3, POLR2E, PPA1, PPP1CB, PPP1R2, PRMT5, PSAT1, PSMD14, PSMD3, PTDSS1, PTMA, PTMS, PTP4A2, PTPN1, PXDC1, RAB14, RAB1A, RAB34, RABAC1, RAN, RANBP2, RHOC, RNASEK, RND3, RNF181, RNF213, RPL10A, RPL12, RPL15, RPL28, RPL31, RPL34, RPL36AL, RPL37, RPL37A, RPL4, RPL7A, RPS11, RPS18, RPS27, RPS29, RPS3, RPS6KA2, RPS9, RRAS, RTN4, S100A13, S1PR1, SAMD4A, SCAMP2, SDHB, SELENOM, SELENON, SELENOT, SEPTIN9, SERF2, SERPINB1, SGK1, SH3BGRL3, SHC1, SLC25A13, SLC25A5, SLIT2, SMARCA2, SNRPG, SON, SPARC, SPCS3, SRPRA, SRSF5, SRSF9, STOML2, SUMF2, SYNGR2, SYPL1, TAF15, TAPBP, TBC1D15, TFPI, TGFBI, THBS1, THOC1, TIMP1, TM9SF3, TMED10, TMSB10, TNFSF10, TNIP1, TNKS1BP1, TNRC18, TNS2, TRA2B, TRAM1, TRIM22, TRIM28, TRIO, TRIR, TSN, TUBA1C, TXNIP, U2AF2, UBA1, UBXN8, UFC1, UQCRC1, USP22, VAMP2, VAPA, VCAN, VIM, VWF, XRCC6, YBX3, YIPF3, YWHAZ, ZFAND5, ZFP36L2, ZYX* |
| KLF6 vs plexiform lesions (Tuder et al.) | *ADAMTS1, ARHGAP10, BCL3, BGN, C12orf75, CD59, CD93, CDH5, COL18A1, COL4A1, COL4A2, CTSB, DDIT4, DEPP1, EMP1, ENG, FOSL2, ICAM2, IFI27, JUNB, LMNA, MAZ, MIDN, MMRN2, MYH10, NEXN, NOS3, NPDC1, PRSS23, TIMP1, TMSB10* |

**Table S5. Differential expression of KLF6 in ACDMPV endothelial cell (EC) populations.**

*KLF6* expression in each EC subtype in ACDMPV was compared to its expression in the corresponding cell type in the 3-year-old or preterm neonate control lungs in the snRNA-seq data (PMID: 37463497). Differential expression was performed using Seurat 4 FindMarkers function with Wilcoxon rank sum test. The following criteria were used for significance: p<0.05, FC>=1.5, pct>20%
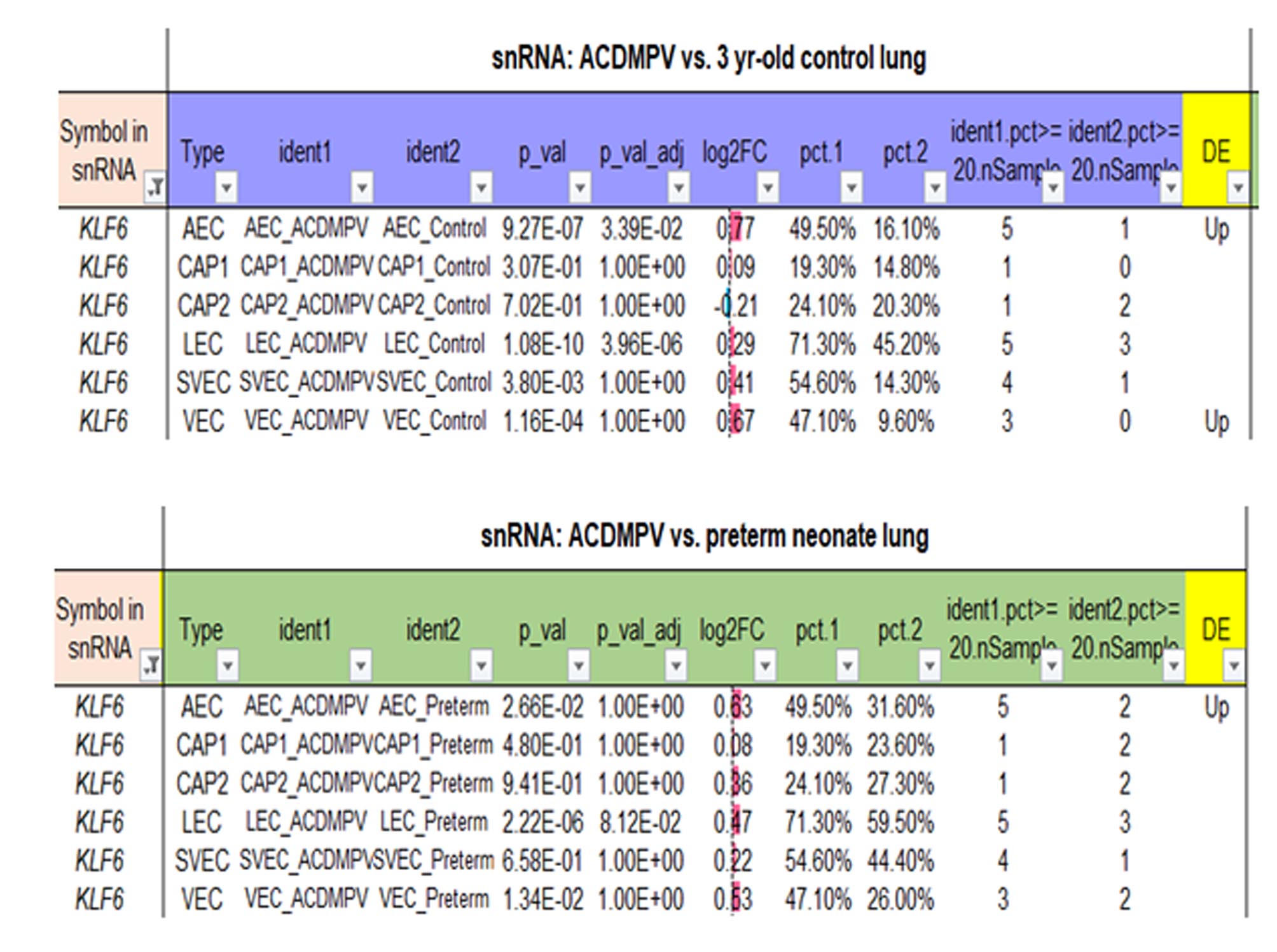
.
